# Diversity and ecology of *Caudoviricetes* phages with genome terminal repeats in fecal metagenomes from four Dutch cohorts

**DOI:** 10.1101/2022.09.02.506393

**Authors:** Anastasia Gulyaeva, Sanzhima Garmaeva, Alexander Kurilshikov, Arnau Vich Vila, Niels P. Riksen, Mihai G. Netea, Rinse K. Weersma, Jingyuan Fu, Alexandra Zhernakova

**Affiliations:** Department of Genetics, University of Groningen, University Medical Center Groningen, the Netherlands; Department of Gastroenterology and Hepatology, University Medical Center Groningen, the Netherlands; Department of Internal Medicine, Radboud University Medical Center, Nijmegen, the Netherlands; Department of Pediatrics, University of Groningen, University Medical Center Groningen, the Netherlands

## Abstract

The human gut harbors numerous viruses infecting the human host, microbes and other inhabitants of the gastrointestinal tract. Most of these viruses remain undiscovered, and their influence on human health is unknown. Here we characterize viral genomes in gut metagenomic data from 1,950 individuals from four population and patient cohorts. We focus on a subset of viruses that is highly abundant in the gut, remains largely uncharacterized, and allows confident complete genome identification – phages that belong to the class *Caudoviricetes* and possess genome terminal repeats. We detect 1,899 species-level units belonging to this subset, 19% of which do not have complete representative genomes in major public gut virome databases. These units display diverse genomic features, are predicted to infect a wide range of microbial hosts, and on average account for < 1% of metagenomic reads. Analysis of longitudinal data from 338 individuals shows that the composition of this fraction of the virome remained relatively stable over a period of 4 years. We also demonstrate that 54 species-level units are highly prevalent (detected in > 5% of individuals in a cohort). Finally, we find 34 associations between highly prevalent phages and human phenotypes, 24 of which can be explained by the relative abundance of potential hosts.

## Introduction

The human gut harbors a large and diverse collection of viruses. These include viruses that infect human cells, viruses that infect archaea and bacteria inhabiting the gut (phages), viruses that infect protists and parasites, and viruses that pass through the intestinal tract with food. The viruses of the human gut belong to diverse lineages and possess different types of genomes: single- or double-stranded (ss or ds) DNA or RNA [1, 2]. The diversity of the human gut virome is actually so large that despite the unprecedented attention to the human gut ecosystem in recent years, saturation in the number of known species of human gut viruses has not been reached [3], and a multitude of unanswered questions about their biology and links to human health remain.

One of the most abundant and diverse groups of viruses in the human gut are the tailed phages unified in the class *Caudoviricetes* [4, 5]. These phages possess dsDNA genomes that encode a distinctive major capsid protein (MCP) with the HK97 fold and a distinctive packaging enzyme, terminase, consisting of a small and a large subunit (TerL) [6, 7]. Phages belonging to class *Caudoviricetes* employ a wide range of replication mechanisms, which is reflected in their genome termini type. The virion-packaged genome molecules of all *Caudoviricetes* phages are believed to be linear. Those that replicate via a circular intermediate possess cohesive ends or direct terminal repeats (DTR) when packaged; if sequenced during replication, these genomes are also likely to produce contigs with DTR [2, 8, 9]. Genomes that replicate by transposition are flanked by random host genome fragments when packaged [8]. Genomes that employ a protein-primed replication mechanism remain linear during replication and possess inverted terminal repeats (ITR) [10]. Importantly, the presence of DTR or ITR at phage contig termini can be used in bioinformatics analysis as an indicator of phage genome sequencing completion [11]. Depending on their lifestyle, many *Caudoviricetes* phages can be referred to as virulent or temperate. Virulent phages enter a lytic state upon genome injection into a host cell: replicate, then lyse the cell to release viral progeny. Temperate phages can enter a lysogenic state (becoming dormant, for example by integrating into the host genome as a prophage) and subsequently switch to a lytic state [12]. Accumulated mutations can render a prophage incapable of switching to a lytic state, turning it into a cryptic prophage [13]. The taxonomic structure of class *Caudoviricetes* is currently undergoing a major overhaul to produce a primarily genome-based classification that accurately reflects the evolutionary relationships between member phages, and order *Caudovirales* and families *Myoviridae, Siphoviridae, Podoviridae* were recently abolished as a part of this effort [5].

During the last decade, metagenomics – analysis of nucleic acid sequences extracted from an entire ecological community – has become the primary method for studying the diversity of the human gut virome. The two popular approaches are sequencing of nucleic acid isolated either from the entire human gut community (total metagenome) or from virus-like particles (virus-enriched metagenome). Both approaches can produce a biased representation of the virome composition. For example, virus-enriched metagenomes do not include the genomes of phages in lysogenic state, while preparation of total metagenome libraries usually does not include the steps required to sequence the genomes of RNA and ssDNA viruses [14]. As metagenomics rapidly increases the amount of sequencing data available, bioinformatics analysis often becomes the bottleneck of virome discovery [11]. One promising approach to meeting this challenge is to use protein markers for virus identification (proteins encoded by viruses but not by cellular organisms, e.g. viral structural proteins) and taxonomic assignment (proteins uniquely encoded by viruses belonging to a specific lineage). Despite the challenges, recent metagenomics studies were able to uncover virome signatures associated with a number of diseases including inflammatory bowel disease (IBD), colorectal cancer and type 1 diabetes [4]. Nonetheless, the role of gut phages in relation to human diseases is still underexplored.

To improve our understanding of the human gut virome, we analyzed viruses in total fecal metagenomes from four cohorts collected in the Netherlands: the population cohorts Lifelines-DEEP (LLD) and LLD follow-up [15-17], a cohort of overweight and obese individuals with BMI > 27 kg/m^2^ (300OB) [18, 19], and a cohort of patients with IBD [20, 21]. We relied on protein markers for virus identification and taxonomic assignment. The analysis was focused primarily on viruses with genome terminal repeats belonging to the class *Caudoviricetes*, and examined their diversity, abundance, stability, predicted hosts and links to human phenotypes.

## Results

### Viral fraction of total fecal metagenomes

To identify the virus-like fraction of the total fecal metagenomes from the LLD (n = 1,135), LLD follow-up (n = 338), 300OB (n = 298) and IBD (n = 520) cohorts, we used Cenote-Taker 2, a tool relying on detection of virus marker genes for virus discovery in sequencing data [22]. We set the software to recognize contigs encoding virion proteins and to cleave off fragments of microbial genomes from these contigs (see Methods). Of the 58,776 virus-like contigs detected (Figure S1), 45% originated from LLD, 21% from LLD follow-up, 15% from 300OB and 19% from IBD. Microbial genome fragments were cleaved off from 15,570 contigs, suggesting that these contigs represent prophages. 1,613 contigs had terminal repeats: 97% had DTR and 3% had ITR. 5,706 and 21 contigs were predicted to employ alternative genetic codes with TAG and TGA stop codons recoded to amino acids, respectively.

Next, we explored the taxonomic composition of the detected virus-like contigs. Predicted proteomes of the contigs were compared to profiles of marker proteins selected to identify seven dsDNA virus groups (Table 1, Text S1, Figure S2). As a result, 39,752 contigs were assigned to class *Caudoviricetes*, one contig was predicted to belong to family *Adenoviridae* (99.9% nucleotide identity to *Human mastadenovirus D* HM770721.2 over the entire 17.4 kb contig) and the remaining 19,023 contigs did not receive any taxonomic assignment.

**Table 1.** Benchmarking of virus detection and taxonomic assignment^a^.

| Taxonomic group | Viral RefSeq genome sequences recognized by Cenote-Taker 2, % | Taxonomic assignment |  |  |
| --- | --- | --- | --- | --- |
|  |  | Marker protein profile(s)<sup>b</sup> | Sensitivity, % | Specificity, %<sup>d</sup> |
| Class *Caudoviricetes* | 99.61 | Terminase large subunit (TerL): VOG00461 and alignments from *Yutin et al. 2021* and *Benler et al. 2021*. Target proteins were required to include hits to the TerL Walker B motif and the TerL nuclease motifs I and II. Target genomes encoding the *Herpesviridae* marker protein were discarded. | 94.29 | 99.98 |
| Family *Herpesviridae* | 82.46 | Major capsid protein (MCP): PF03122 | 79.82 | 100 |
| Family *Papillomaviridae* | 0 | Capsid protein L1: VOG05075 | 99.52 | 100 |
| Family *Polyomaviridae* | 0.76 | Coat protein: PF00718 | 99.24 | 100 |
| Family *Adenoviridae* | 91.89 | Hexon protein: VOG05391 | 91.89 | 100 |
| Class *Tectiliviricetes* (excluding the *Adenoviridae* family) | 89.47 | Double Jelly-Roll MCP alignments corresponding to six prokaryotic virus groups<sup>c</sup> from *Yutin et al. 2018* | 100 | 99.99 |
| Phylum *Nucleocytoviricota* | 74.04 | Seven marker protein alignments from *Aylward et al. 2021*; each genome hit by a marker protein profile was further analyzed using ViralRecall (see Methods) | 70.19 | 99.98 |

<sup>a</sup> See Text S1.
<sup>b</sup> Marker protein alignment identifiers beginning with “VOG” and “PF” refer to the VOG and Pfam database entries, respectively.
<sup>c</sup> Two of the groups, Odin and FLiP, are divergent from the recognized members of the class *Tectiliviricetes*.
<sup>d</sup> Unclassified viruses were excluded from consideration when calculating the number of true negatives and false positives.

In order to represent the gut virome as comprehensively as possible, we incorporated genomes from existing viral databases [3, 23-31] (Table 2) into the analysis. To avoid redundancy, both the viral genomes from the databases and the virus-like contigs identified in the four Dutch cohorts were clustered into virus operational taxonomic units (vOTUs) together. As a result, 30,461 vOTUs were delineated (Figure S1). One sequence per vOTU (with terminal repeats if available) was selected as a vOTU representative.

**Table 2.**
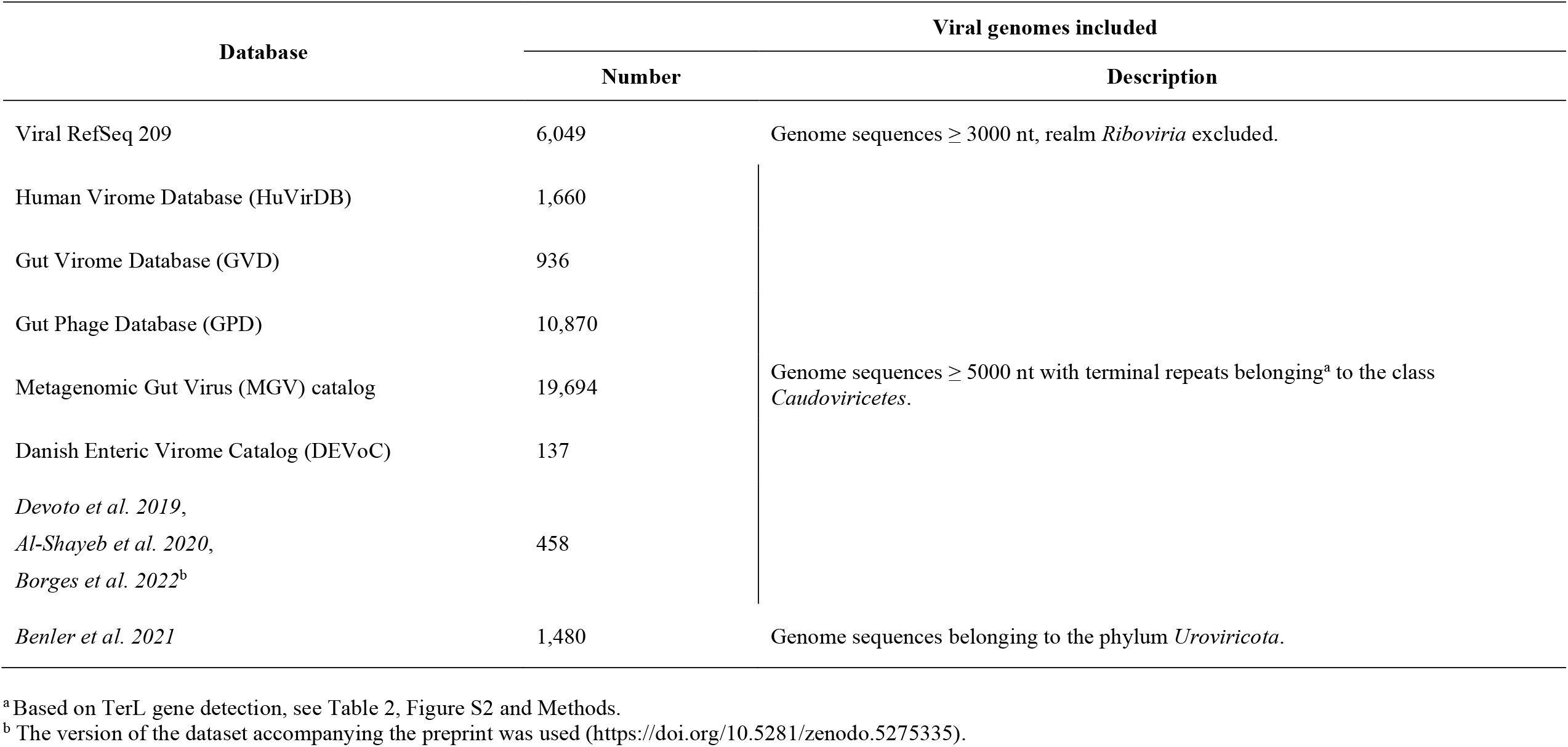
Virus genomes from databases included in the analysis.

To estimate the relative abundance of the vOTUs in metagenomic samples, sequencing reads from individual samples were mapped to the genome sequences representing the vOTUs. A vOTU was considered detected if ≥ 75% of its representative sequence length was covered by reads. In all, 15,196 vOTUs were detected in at least one sample from the four Dutch cohorts (Figure S1). Based on the congruent taxonomy of their members, 69% of these vOTUs were assigned to class *Caudoviricetes*, 31% did not receive any taxonomic assignment, and the remaining 7 vOTUs included ssDNA prokaryotic viruses from family *Microviridae* (likely sequenced while in a DNA duplex state during replication) and dsDNA human viruses from families *Papillomaviridae, Polyomaviridae* and *Adenoviridae*.

Analyzing the entire set of virus-like sequences detected in total metagenome sequencing data (Figure 1) poses a number of challenges. When analyzing virus-like genomes that lack terminal repeats, it is difficult to estimate their completeness and to distinguish between prophages that can excise from the host genome and enter a lytic state and cryptic prophages that have lost this ability. When considering virus-like genomes with DTR that did not receive any taxonomic assignment, it can be difficult to distinguish these from plasmids.

**Figure 1.**
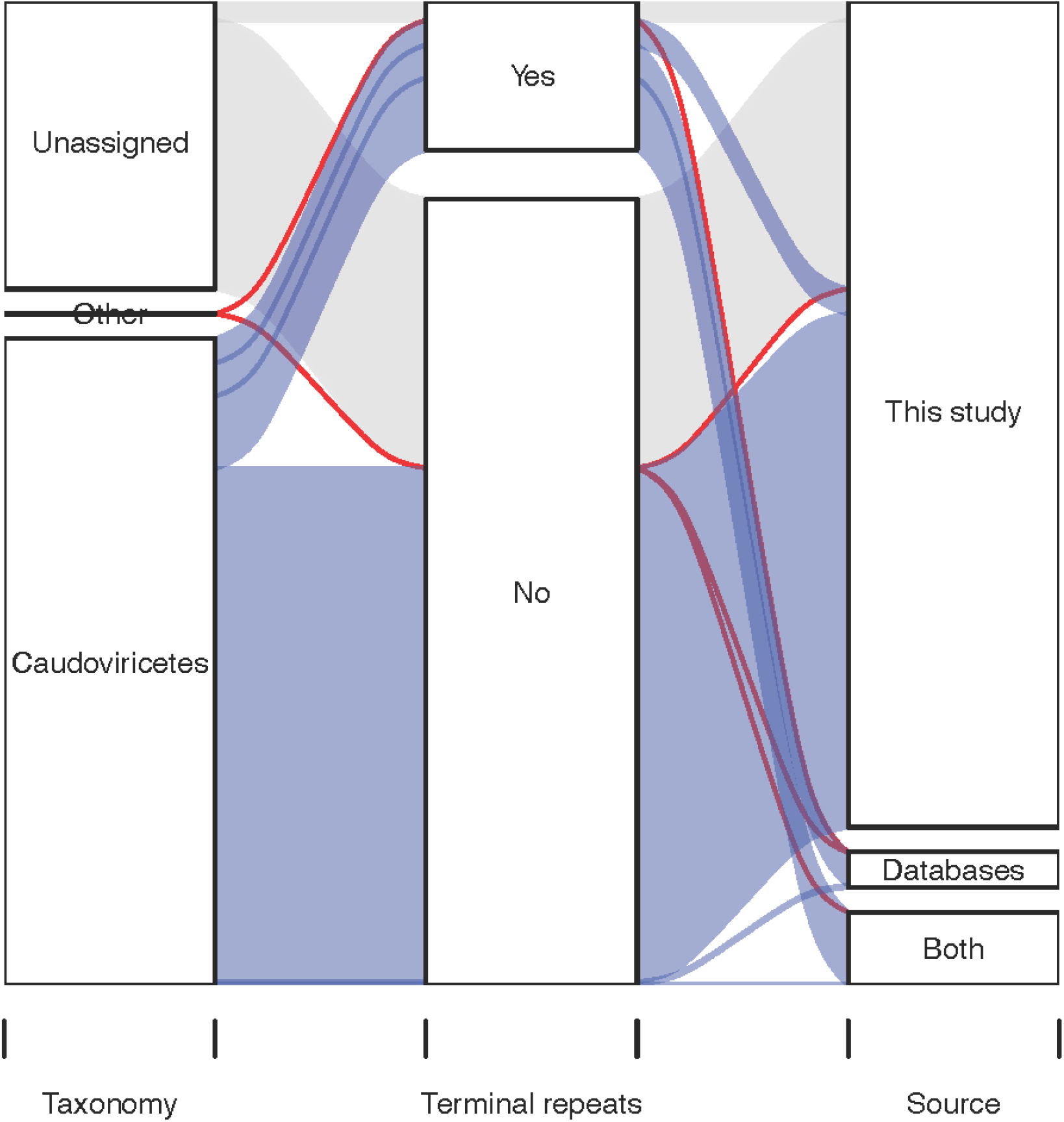
Properties of the vOTUs detected in the four Dutch cohorts. Sankey diagram shows the relationships between taxonomy, detection of terminal repeats in representative genome and source of the 15,196 vOTUs.

Notably, 20% of the taxonomically unassigned vOTUs represented by sequences with DTR were predicted to be plasmids with high confidence (PlasX score > 0.9 [32]), whereas the same was the case for only 2% of the *Caudoviricetes* vOTUs represented by sequences with DTR.

We therefore decided to focus on *Caudoviricetes* vOTUs represented by sequences with terminal repeats (Figure 1). Initially there were 2,106 such vOTUs, but after excluding a highly prevalent vOTU represented by a chimeric nucleotide sequence (NL_vir005341) and vOTUs with an undetected or dubious TerL gene in the representative genome (see Methods, Figure S2), 1,899 vOTUs remained (Table S1). Below, we refer to genome sequences belonging to these vOTUs as the Caudoviricetes Genomes with Terminal Repeats (CGTR1899) database.

### Diversity of *Caudoviricetes* phages with genome terminal repeats

We next aimed to characterize the diversity, abundance and long-term stability of the *Caudoviricetes* phages with genome terminal repeats represented by the CGTR1899 database in the four Dutch cohorts. To compare the abundance of phages in the total metagenomes from the four Dutch cohorts to estimates based on virus-enriched metagenomes, we explored a collection of 254 Danish fecal viromes [28].

The CGTR1899 database constitutes only 12% of the vOTUs detected in the four Dutch cohorts based on read alignment but encompasses 29% of all virus-like contigs assembled for these four cohorts. The CGTR1899 database includes vOTUs composed entirely of sequences from the four cohorts (19%), entirely of sequences from the databases (25%), or of a mixture of both (56%). 404 CGTR1899 vOTUs were detected in the Danish fecal viromes, providing additional confirmation for the viral nature of these vOTUs (Table S2).

We measured the abundance of CGTR1899 phages per sample both as the number of viruses detected and the percentage of recruited reads. The mean number of CGTR1899 vOTUs detected in a sample was 7 for LLD, 10 for LLD follow-up, 9 for 300OB and 5 for IBD. On average, genomes representing CGTR1899 vOTUs recruited 0.68% (LLD), 0.75% (LLD follow-up), 0.74% (300OB) and 0.60% (IBD) reads per sample (Figure 2A). When we compared the CGTR1899 phage abundances in the four Dutch cohorts to those in the Danish healthy adult viromes (n = 52), the mean number of vOTUs detected was similar (6) but the mean read recruitment rate for the Danish samples was considerably higher (6.92%), as could be expected for these virus-enriched samples.

**Figure 2.**
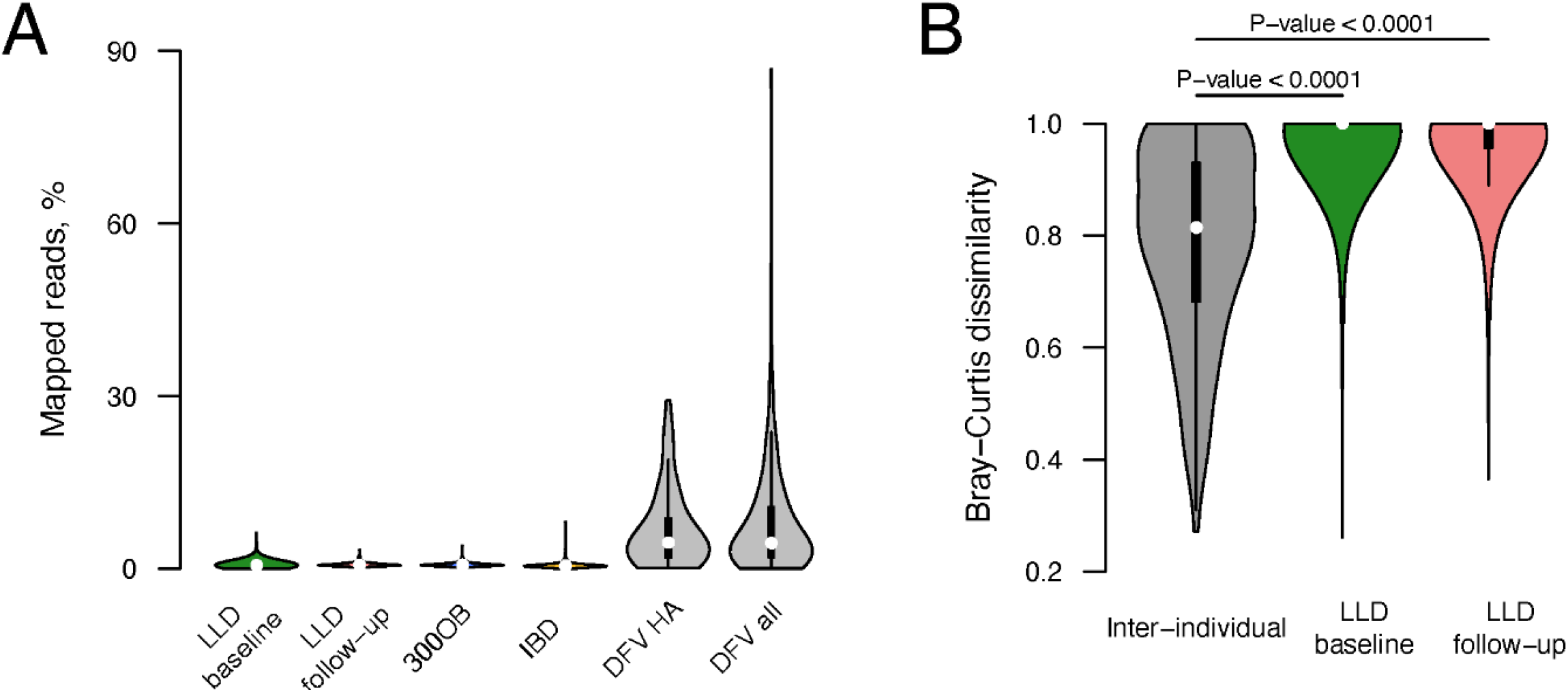
Abundance and stability of the CGTR1899 phages. (**A**) Violin plots show the percent of sample reads mapping to the genomes representing CGTR1899 vOTUs for the four Dutch cohorts (LLD baseline, LLD follow-up, 300OB and IBD), healthy adult Danish fecal viromes (DFV HA) and all Danish fecal viromes (DFV all). (**B**) Bray-Curtis dissimilarities between pairs of samples collected from the same individual 4 years apart (gray) and from different individuals at the same time point (green and red). Empirical P-values are indicated above the violin plots. Data from the 338 individuals sampled as part of the LLD and LLD follow-up cohorts were utilized.

We estimated the stability of the CGTR1899 fraction of the virome based on longitudinal data from the LLD and LLD follow-up cohorts. The Bray-Curtis dissimilarity between samples collected from the same individual 4 years apart was significantly lower than that between samples collected from different individuals at the same timepoint (P-value < 0.0001, Figure 2B), indicating relative stability. Importantly, as virus detection was based on total metagenome data, it was impossible to distinguish between phages in lytic and lysogenic states, and thus stability was estimated for phages in all states taken together.

The diversity of the CGTR1899 phages was assessed via characteristics of the genomes representing vOTUs such as genome length, GC content and genetic code. Notably, genomes with similar characteristics tended to cluster on a TerL-based phylogenetic tree (Figure 3, Table S1). The length of the genomes representing vOTUs ranged from 5,061 to 352,502 nt, mean 56,745 nt. GC content varied from 25% to 69%, mean 44%. A minority of the genomes representing vOTUs (90, 5%) possessed ITR at their termini; the rest possessed DTR. 1,608 genomes representing vOTUs were predicted to employ standard bacterial genetic code, with the remaining 285 and 6 predicted to employ alternative genetic codes with a TAG or TGA stop codon recoded to amino acid, respectively. Interestingly, transfer RNA (tRNA) genes with an anticodon matching TAG were detected in just 13% of the 285 genomes predicted to be TAG-recoded. There were 395 vOTUs containing phage genome sequences originating from prophage contigs (i.e. contigs including phage and host genome fragments) identified in the four Dutch cohorts (Table S1).

**Figure 3.**
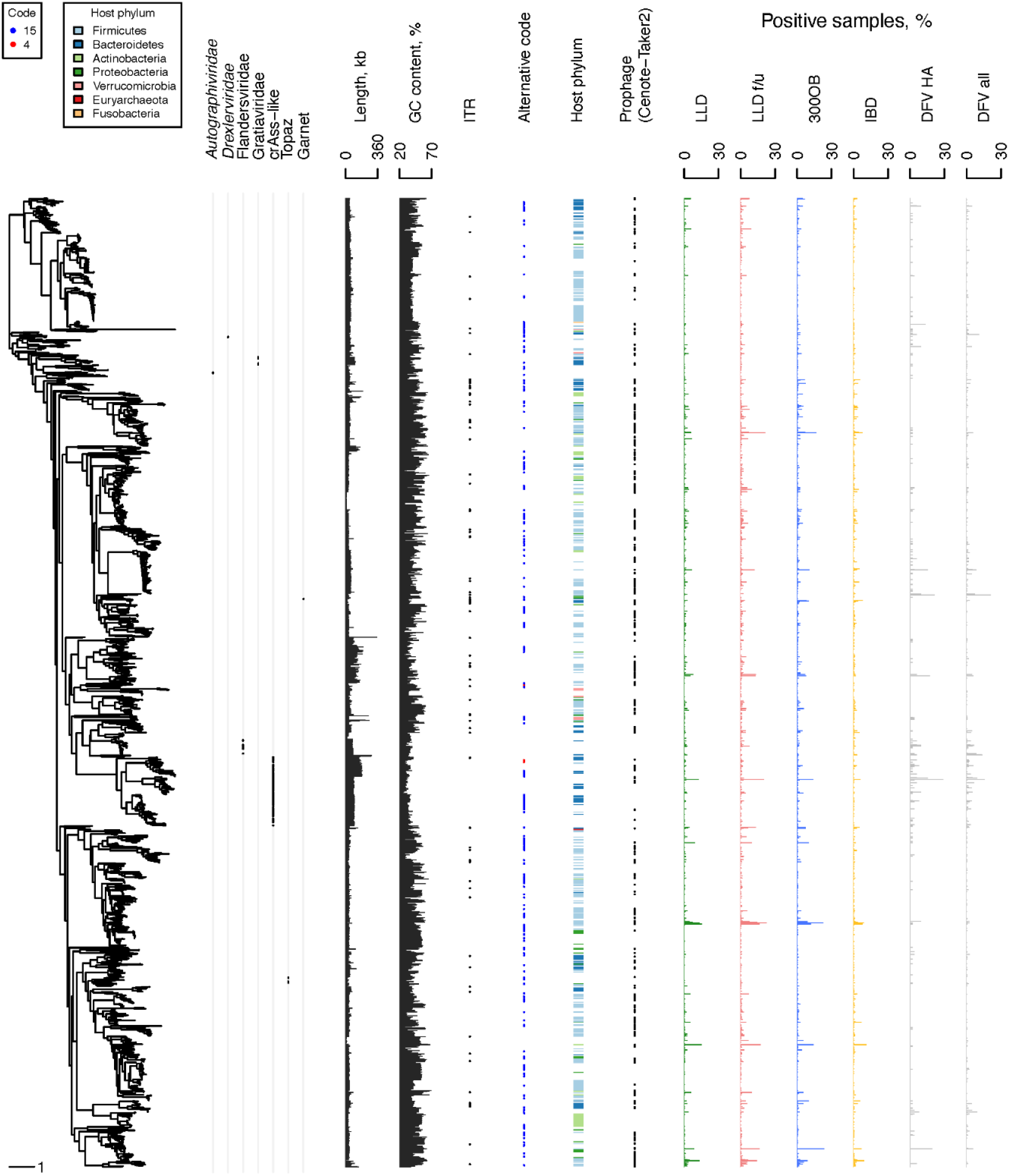
Properties of the CGTR1899 phages. Left, a phylogenetic tree reconstructed based on the TerL proteins of phages representing the CGTR1899 vOTUs. From left to right, the following genome properties are depicted per tree tip: assignment to family-level taxonomic groups, length, GC content, terminal repeats (presence or absence of a black dot indicates ITR or DTR, respectively), predicted genetic code (blue dot for code 15, red dot for code 4 or empty space for standard code 11), predicted host phyla designated by colored bars (empty space if prediction is unavailable), presence of vOTU members derived from prophage contigs identified by Cenote-Taker2, and prevalence in the four Dutch cohorts (LLD, LLD follow-up, 300OB and IBD), healthy adult Danish fecal viromes (DFV HA) and all Danish fecal viromes (DFV all).

Taxonomic assignment of the CGTR1889 phages proved to be very sparse. The majority of the CGTR1889 vOTUs could not be assigned to an established monophyletic group at the level of family or order. Only 7% of the vOTUs were assigned to such groups based on the presence of previously classified genomes within the vOTUs (Figure 3, Table S1). Extending assignment by finding the most recent common ancestor (MRCA) of all vOTUs assigned to a group on the TerL-based phylogenetic tree, and then placing all descendants of the MRCA into the group, resulted in 11% of vOTUs being taxonomically assigned: family *Autographiviridae* (4 vOTUs), family *Drexlerviridae* (2 vOTUs), Flandersviridae (also known as Gubaphages, 32 vOTUs) [2, 24, 27], Gratiaviridae (16 vOTUs) [24], crAss-like phages (135 vOTUs), group Topaz (18 vOTUs) and group Garnet (1 vOTU) [31].

Next, we predicted the hosts of the CGTR1899 phages using two methods: (1) analysis of the host genome fragments attached to prophage contigs identified in the four Dutch cohorts and (2) detection of sequence similarity between phage genomes and microbial CRISPR spacers (see Methods). The first approach yielded predictions for 228 vOTUs. The second approach yielded predictions for 578 vOTUs. In total, predictions were made for 713 vOTUs (Table S3). Predictions were made by both methods for 93 vOTUs. In 88 cases, the predictions made by both methods were identical at the host phylum level. In the remaining five cases, the predictions were incompatible at host phylum level. In these cases, we prioritized the prophage-based predictions. Predicted hosts belonged to the phyla Firmicutes (453 vOTUs), Bacteroidetes (138 vOTUs), Actinobacteria (60 vOTUs), Proteobacteria (47 vOTUs), Verrucomicrobia (13 vOTUs), Fusobacteria (1 vOTU) and Euryarchaeota (1 vOTU).

Prevalence of the individual CGTR1899 phages in the four Dutch cohorts varied depending on the phage. Some were detected in a single sample, whereas others were detected in dozens of samples (Figures 3 and 4, Table S2). The most prevalent vOTUs per cohort were NL_vir026707 in LLD (detected in 16% of samples), NL_vir053139 in LLD follow-up (detected in 22% of samples), MGV-GENOME-0279285 in 300OB (detected in 23% of samples) and OLXK01000549.1 in IBD (detected in 11% of samples). Unsurprisingly, the prevalence of the CGTR1899 phages in the Danish fecal viromes was often drastically different from that in the four Dutch cohorts. For example, the second most prevalent CGTR1899 vOTU among the Danish healthy adult viromes was MGV-GENOME-0193745 (21% positive samples), which was detected in < 1% samples in every Dutch cohort. This vOTU includes genomes of the virulent *Leuconostoc* phages ΦLN03, ΦLN04 and ΦLN12 [33, 34] that could be expected to be abundant among DNA sequences isolated from virus-like particles.

**Figure 4.**
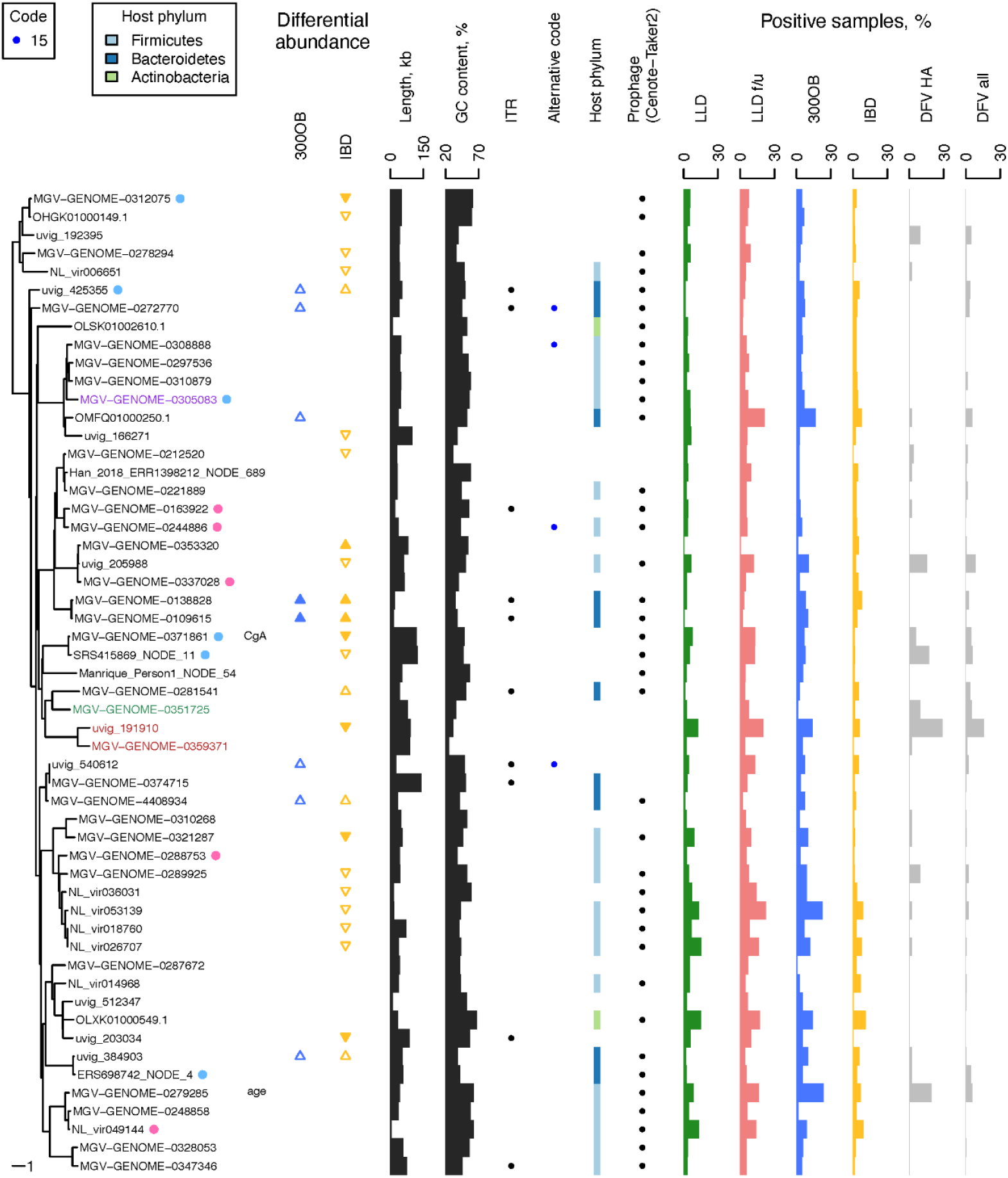
Properties of the CGTR54 phages. This figures shows a subset of the data presented in Figure 3 including only information about the 54 vOTUs detected in >5% samples of a Dutch cohort. See legend of the Figure 3 for details. The crAss-like phage, flandersvirus and *Faecalibacterium* phage Lagaffe vOTU names are written in red, green and violet font, respectively. A dot next to a vOTU indicates that a phage with a similar genome sequence was described in *Minot et al. 2012* (pink) or *Dzunkova et al. 2019* (blue) (see Figure S3). A name of a phenotype (“CgA”, “age”) next to a name of a vOTU indicates a phage–phenotype association in the LLD cohort. Statistically significant differences in prevalence of vOTUs between the population cohort LLD and patient cohort 300OB (IBD) are indicated by blue (yellow) triangles. If a vOTU was overrepresented in a patient cohort, the triangle points upward. If it was underrepresented, the triangle points downward. If the association was significant after the logistic regression was adjusted for relative abundance of the predicted host, the triangle is filled, otherwise the triangle is empty. Contig length and coverage are omitted from the phage names for brevity, where applicable.

### Diversity of the most prevalent *Caudoviricetes* phages with genome terminal repeats

In the final part of the analysis, we focused on the most prevalent CGTR1899 phages: 54 vOTUs detected in > 5% samples in at least one of the four Dutch cohorts (and referred to as the CGTR54 database below). We explored their diversity, searched for known closely related viruses and analyzed associations between the prevalence of these phages and human phenotypes. Importantly, since only total metagenome sequencing data are available for the four Dutch cohorts, we could not determine if each detected phage was in lytic or lysogenic state in a particular sample. However, it is worth noting that 35 of the 54 vOTUs were detected in the Danish fecal viromes (Figure 4), underscoring that these vOTUs can exist in the form of virus particles.

The diversity of the CGTR54 phages can be assessed through the characteristics of their representative genomes. The lengths of the representative genomes varied from 5,061 to 140,662 nt, and their GC content varied from 25% to 67%. 44 of the CGTR54 vOTUs were represented by genomes with DTR. The remaining 10 vOTUs were represented by genomes with ITR. Four representative genomes were predicted to use an alternative genetic code with TAG stop codons recoded to amino acid. Finally, 38 vOTUs included sequences originating from prophage contigs (Figure 4).

The diversity of the CGTR54 phages was also reflected in the organization of their representative genomes (Material S1). There were representative genomes with the majority of open reading frames (ORFs) positioned on a single strand, as well as genomes with ORFs occupying both the forward and reverse strand. The 54 vOTUs were represented by genomes with various shapes of AT- and GC-skew curves, including the V-shaped cumulative GC-skew curve previously suggested to be associated with bi-directional replication [35]. Some of the 54 genomes were evenly covered by reads, while in others we observed an anomaly where a specific region receives almost no coverage in a fraction of samples (Material S1), which may be a consequence of recombination [36]. Inspection of the genome maps also revealed that forward and reverse strand sequences of one of the genomes (MGV-GENOME-0281541) are completely identical, indicating that this genome might be a sequencing artifact (Material S1).

Despite the diversity of the 54 genomes, there were some shared characteristics. Genes with similar function, such as tRNA genes or genes encoding structural proteins and proteins implicated in assembly of virus particles, tended to form clusters (Material S1). Genes encoding integrases were detected in multiple genomes. Potential diversity-generating retroelements – a reverse transcriptase (RT) gene and a pair of nucleotide repeats with at least one repeat positioned in close proximity to the RT gene [37] – were detected in 21 genomes (Material S1). Interestingly, in multiple genomes there was a pair of repeats with both repeats close to the RT gene and a pair of repeats with one repeat close to the RT gene and another separated from the RT gene by a large distance (Material S1, S2).

Taxonomic assignment was only possible for 4 CGTR54 vOTUs (Figure 4). MGV-GENOME-0359371 and uvig_191910 vOTUs were recognized as crAss-like phages because of the presence of previously described crAss-like phage genomes in these vOTUs (Table S1) [36]. The MGV-GENOME-0359371 vOTU belongs to the same clade of crAss-like phages as crAssphage *sensu stricto* – clade alpha – although the nucleotide sequence similarity between them is relatively low (only 27% of the crAssphage *sensu stricto* genome was covered by hits when compared to the MGV-GENOME-0359371 genome using BLASTN with E-value threshold 0.05) [38]. The uvig_191910 vOTU belongs to the gamma clade of crAss-like phages [39, 40]. For the other two CGTR54 vOTUs, the MGV-GENOME-0351725 vOTU was recognized as belonging to the Flandersviridae group because it contains flandersvirus OJML01000036 [24] and the MGV-GENOME-0305083 vOTU was recognized as *Faecalibacterium* phage Lagaffe because it contains the genome of this phage, NC_047911 [41].

To investigate if any of the CGTR54 vOTUs had already been extensively characterized, we searched the viral fraction of GenBank for similar sequences (see Methods). We found that five viral contigs that had been revealed to contain diversity-generating retroelements in [42] exhibited various degrees of similarity to six genomes representing CGTR54 vOTUs. In addition, eight phage sequences obtained in a project where single-cell viral tagging was used to identify unknown host–phage pairs [43] demonstrated similarity to five genomes representing CGTR54 vOTUs. Finally, contig71, which was subjected to PCR amplification and Sanger sequencing in [44], displayed similarity to a fragment of the CGTR54 flandersvirus (Figure 4, Figure S3, Table S4).

Hosts of the CGTR54 phages predicted based on the analysis of prophage contigs and CRISPR spacers included phyla Firmicutes (20 vOTUs), Bacteroidetes (10 vOTU) and Actinobacteria (2 vOTUs) (Figure 4). We also predicted the hosts of the CGTR54 phages based on co-abundance with microbial taxa. The potential host of each vOTU was predicted as the microbial taxon demonstrating the most reliable (minimal false discovery rate (FDR) in meta-analysis) relative abundance correlation with the vOTU (Table S3). Notably, there were 32 vOTUs with a prophage-based and/or CRISPR-based prediction available in addition to the co-abundance-based prediction, and we observed agreement between all available predictions at host phylum level for 91% of these vOTUs (Table S3).

### Associations with human phenotypes

The availability of phenotypic data for participants of the four Dutch cohorts provides an opportunity to explore associations between detection of gut phages and human phenotypes. We explored the associations of the CGTR54 vOTUs using logistic regression where the presence of the phage represents the outcome and the phenotype represents the predictor, while adjusting for the age and sex of cohort participants. Subsequently, we conducted an additional analysis where the logistic regression was additionally adjusted for the abundance of the potential host predicted by co-abundance analysis. Associations were considered significant at a FDR < 0.05.

The association analysis based on the LLD cohort data revealed a negative association between the fecal level of the secretory protein chromogranin A (CgA) and detection of the MGV-GENOME-0371861 vOTU, and a positive association between the age of human subjects and detection of the MGV-GENOME-0279285 vOTU (Figure 4, Table S5). However, both associations were no longer significant after adjustment for the abundance of the predicted hosts.

Eight vOTUs were found to be significantly more prevalent in overweight and obese individuals (300OB cohort, BMI > 27 kg/m^2^) compared to the general population (LLD cohort). After adjustment for the abundance of the predicted hosts, only two of these associations remained statistically significant (Figure 4, Table S5).

Seven vOTUs were found to be significantly more prevalent among IBD cohort participants compared to the general population (LLD cohort), while 16 vOTUs were found to be significantly less prevalent. After the adjustment for the abundance of the predicted hosts, these numbers changed to 3 and 5 vOTUs, respectively (Figure 4, Table S5).

Overall, these results indicate that in many cases the driving force behind the association may not be the phage itself but rather its microbial host.

## Discussion

We used a marker-based bioinformatics approach to identify and classify viral genomes in fecal metagenomes from four Dutch cohorts: two population cohorts, a cohort of overweight and obese individuals and a cohort of IBD patients. Detected viruses included those belonging to class *Caudoviricetes* and families *Microviridae, Papillomaviridae, Polyomaviridae* and *Adenoviridae*, and we further focused specifically on *Caudoviricetes* phages with genome terminal repeats. We estimated the proportion of their nucleic acid in the human gut metagenomes (< 1% on average), noted the relative stability of this virome fraction over a period of 4 years and described the diversity of these viruses, including their genome characteristics, predicted hosts and prevalence in human gut metagenomes. A small fraction of the *Caudoviricetes* phages with genome terminal repeats were highly prevalent (detected in > 5% of Dutch cohort samples), allowing us to conduct a statistical analysis that identified associations between the prevalence of these phages and human phenotypes including age, fecal levels of CgA, obesity and IBD diagnosis.

Metagenomics is a powerful approach to study viruses that allows us to see the big picture of the human gut virome. However, technical challenges on each step of the study, from sample collection to bioinformatics analysis, may influence the results. Working with fecal samples, as opposed to intestinal wall biopsy samples, may affect the ratio of the number of viruses infecting microbes and human cells. Conducting total metagenome sequencing, as opposed to virus-enriched metagenome sequencing, means that viruses with dsDNA genomes will be included into the analysis even if they are in a lysogenic state, while viruses with ssDNA and RNA genomes will be excluded. Subsequent bioinformatics analysis carries its own limitations. The marker-based approach that we used to identify virus genomes and tentatively assign them to taxonomic groups is designed to be very specific as it relies on the presence of protein genes uniquely associated with a particular group of viruses. However, it also has limitations. Marker proteins can only be identified based on the known virosphere, so it is always possible that a seemingly unique association between a protein gene and a group of viruses will be disproven with the discovery of novel viruses. Likewise, the definitions of virus taxonomic groups may be shifting with the expansion of the known virosphere [5]. Another important aspect of our bioinformatics analysis was the identification of genomes with DTR and ITR, which also carries several potential pitfalls. It is possible to overlook terminal repeats if they are shorter than the threshold of 20 nt or contain a sequencing error making the 5’- and the 3’-terminal repeat sequences non-identical. On the other hand, a circular plasmid with an integrated prophage or a partial viral genome flanked by repeats can be mistakenly identified as a complete viral genome with terminal repeats. Predicting ORFs is yet another virus bioinformatics challenge that we encountered: some phages employ alternative genetic codes with stop codon reassignment, and thus their ORFs cannot be correctly predicted by standard tools [31, 39, 45]. We solved this challenge by applying an approach similar to the one described in [3, 45], which worked well on a crAss-like phage test dataset (see Methods). Notably, the percent of viral contigs from the four Dutch cohorts predicted to employ an alternative genetic code (9.74%) was higher than reported in the literature based on the gut microbiomes of people consuming a westernized diet (2.25%) [31].

This study was specifically focused on phages that belong to class *Caudoviricetes* and possess genome terminal repeats. We used *Caudoviricetes* TerL detection as an indicator that a virus belongs to this group, while requiring that the genome in question does not encode the marker of the family *Herpesviridae*, as herpesviruses possess an evolutionarily-related TerL [6]. Detection of TerL was required to involve three TerL motifs: the Walker B motif belonging to the adenosine triphosphatase domain and motifs I and II belonging to the nuclease domain (Figure S2). This approach performed well in benchmarking, reaching a sensitivity of 94.3% and a specificity of 99.9%, although with the caveat that the Viral RefSeq database used for benchmarking contained sequences employed in the TerL detection procedure, leading to a potential overestimation of the approach’s robustness (Text S1). Notably, the sensitivity did not reach 100%, and there may be several reasons for that. Not all phages belonging to the class *Caudoviricetes* encode TerL: for example, a satellite phage may lack terminase genes and employ a terminase enzyme of a helper phage instead [46]. Alternatively, a phage might possess a packaging enzyme that differs from the terminase of the majority of known *Caudoviricetes* phages [47]. The TerL gene may also be interrupted by an intron [48]. Finally, a TerL protein encoded by a divergent *Caudoviricetes* phage may fail to be recognized based on sequence similarity. On the other hand, given the diversity and mosaicism of viral genomes, it is impossible to exclude a possibility that uncharacterized viruses outside of class *Caudoviricetes* may encode close homologs of the *Caudoviricetes* TerL.

One of the most intriguing aspects of the human gut virome is its potential role in human health and disease. In this study we found a positive correlation between detection of a phage and human age, and a negative correlation between detection of a phage and fecal levels of CgA, which is a precursor to peptides with regulatory and antimicrobial activities [49]. We also found that multiple phages are underrepresented or overrepresented in metagenomes of overweight and obese people and in patients with IBD. However, the interpretation of these results is not straightforward. Although we do not know the true host of each of the viruses in question, when we adjusted our statistical model for the relative abundance of their predicted hosts, most of the associations became statistically insignificant. This strongly suggests that the microbial host, and not its phage, may be the driving force behind many of the associations we detected. This seems especially logical because we were working with total metagenome data and thus could not differentiate between detection of a phage in an actively replicating lytic state versus one in a dormant lysogenic state.

To summarize, while the big picture of the human gut virome painted with the help of metagenomics is very informative, further research and improvement of the analysis techniques in the future can help resolve remaining uncertainties.

## Methods

### Virus detection in metagenomes

Total metagenome sequencing data from 2,291 samples from four Dutch cohorts were assembled into contigs as described in [36]. To identify viral genomes, contigs from each sample were screened using Cenote-Taker 2 version 2.1.3, program *unlimited_breadsticks*.*py* with the following parameters: “--virus_domain_db ‘virion’ --minimum_length_circular 3000 --minimum_length_linear 10000 --circ_minimum_hallmark_genes 1 -- lin_minimum_hallmark_genes 2 --prune_prophage True --filter_out_plasmids True” [22]. Identified virus-like contigs were then screened for the presence of ribosomal RNA (rRNA) genes using a BLASTN 2.10.1+ [50] search in the SILVA 138.1 NR99 rRNA genes database [51] with an E-value threshold of 0.001. An rRNA gene was considered to be detected in a contig if the gene and the contig produced a hit covering > 50% of the gene length. The eight contigs with detected rRNA genes were excluded from further consideration. Importantly, 520 IBD cohort samples were screened for the presence of viral genomes, but we excluded 62 samples from individuals with stoma and ileoanal pouches, 1 duplicated sample and 2 samples without metadata from all subsequent analyses, bringing the number of IBD samples under consideration down to 455.

### Nucleotide sequence characterization

A nucleotide sequence was considered to contain a DTR or ITR if identical terminal repeats ≥ 20 nt were detected [52]. Nucleotide content, GC- and AT-skew were calculated using a 1,001 nt window sliding along the genome sequence with a 200 nt step, as described in [36]. Prediction of tRNA genes was conducted for individual genome sequences using tRNAscan-SE 2.0.9 with the “-B” parameter [53]. To search for nucleotide repeats, genome sequences were compared to themselves using BLASTN 2.12.0+ with the “-task ‘blastn’ -evalue 0.001” parameters [50], and only hits with alignment length ≥ 100 and identity ≥ 80% were considered.

### Identification of potential plasmids

Nucleotide sequences with DTR characterized by a PlasX score > 0.9 were considered potential plasmids [32].

### Genetic code prediction

ORFs were predicted using Prodigal 2.6.3 [54]. For sequences shorter than 20 kb, the prediction was made in the “meta” mode using standard bacterial genetic code 11. For each individual sequence ≥ 20 kb, the prediction was made in the “single” mode using standard bacterial genetic code 11 and alternative genetic codes 4 (TGA codon encodes tryptophan) and 15 (TAG codon encodes glutamine). If the sum of the ORF coding potential scores was higher under an alternative genetic code and exceeded the sum under the standard genetic code by 10%, the alternative genetic code was assigned to the viral genome sequence [3, 45]. We tested our method on 378 crAss-like phage genome sequences for which the genetic code had been previously predicted using manual ORF analysis [36] and obtained an identical prediction in 97% of cases.

### Proteome annotation

Individual proteomes, predicted as described above, were compared to Pfam 35.0 profiles [55] using the HMMER 3.3.2 program *hmmsearch* with the “--max -E 0.001” parameters (http://hmmer.org/). Only protein–profile pairs where ≥ 100 amino acid residues of the protein were covered by hit(s) to the profile were considered, the profile providing maximal coverage was used for annotation. Coverage was measured in HMMER envelope coordinates combined using the R package *IRanges* 2.22.2 in case of overlap [56]. Multiple sequence alignment (MSA) of proteins annotated as reverse transcriptases was generated by adding them to the Pfam 35.0 seed PF00078 alignment with the help of the R package *seqinr* 3.6-1 and MAFFT 7.453 with an “--add” parameter [57, 58].

### Taxonomic assignment based on marker genes

Taxonomic assignment was performed using the proteomes predicted as described above. Proteomes were compared to profiles of marker proteins of seven dsDNA virus groups using the HMMER 3.3.2 program *hmmsearch* with the “--max -E 0.001” parameters. The following MSAs were used to generate marker profiles (Table 1): class *Caudoviricetes –* alignment of 5130 TerL sequences from [24], alignment of 823 TerL sequences from [39], TerL MSA VOG00461 from the VOG 207 database [59]; family *Herpesviridae –* MCP MSA PF03122 from the Pfam 35.0 database [55]; family *Papillomaviridae –* capsid protein L1 MSA VOG05075 [59]; family *Polyomaviridae –* coat protein MSA PF00718 [55]; family *Adenoviridae –* hexon protein MSA VOG05391 [59]; class *Tectiliviricetes* excluding the *Adenoviridae* family *–* 6 MCP MSAs corresponding to 6 prokaryotic virus groups from [60], and phylum *Nucleocytoviricota –* 7 marker MSAs (MCP, DNA-directed RNA polymerase alpha and beta subunits, DNA polymerase family B, transcription initiation factor IIB, DNA topoisomerase II, poxvirus late transcription factor VLTF3) from [61]. The profiles were visualized using Skylign [62]. For class *Caudoviricetes*, alignments between the TerL queries and a target protein were required to span the following TerL profile residues: (1) the second conserved acidic residue of the TerL Walker B motif, (2) the first conserved acidic residue of the TerL nuclease motif I, and (3) the conserved acidic residue of the TerL nuclease motif II (Figure S2); if the genome sequence encoding the target protein also encoded a protein producing a hit with the *Herpesviridae* MCP, taxonomic assignment was based on the latter. In case of the phylum *Nucleocytoviricota*, genome sequences encoding proteins hit by any of the seven marker profiles were analyzed by ViralRecall with the “--contiglevel --evalue 1e-3” parameters [63], and an assignment to this phylum was made only if the ViralRecall score was > 2 and ViralRecall was able to detect at least one marker protein gene.

### Species-level clustering

Viral nucleotide sequences were clustered into vOTUs using the CheckV 0.7.0 script *aniclust*.*py* with the “--min_ani 95 --min_qcov 0 --min_tcov 85” parameters [64, 65]. If a vOTU contained genomes with terminal repeats, the median length genome with terminal repeats was selected as a vOTU representative. Otherwise, the longest genome without terminal repeats (encoding TerL, if available) was selected.

### Read mapping

Sequencing reads of each individual sample from the four Dutch cohorts and from the collection of 254 Danish fecal viromes [28], filtered and quality-trimmed as described in [36], were competitively mapped to a database of 30,461 virus-like genome sequences representing vOTUs using Bowtie2 2.4.4 with a “--very-sensitive” parameter [66]. Breadth of genome coverage by reads was calculated using the BEDTools 2.29.2 command *coverage* [67]. Depth of genome coverage by reads was calculated using the SAMtools 1.10 command *depth* [68]. Abundance of a vOTU in a sample was considered to be zero if the breadth of the representative genome coverage by reads was below 75% [69], otherwise it was estimated as (*N* · 10^6^)/(*L* · *S*), where *N* is the number of reads mapped to a genome, *L* is the length of a genome and *S* is the number of sample reads after filtering and quality trimming.

### Building TerL MSA

We used the HH-suite 3.3.0 command *hhalign* with the “-M 50 -mact 0 -all” parameters and script *reformat*.*pl* [70] to first combine TerL alignment VOG00461 from VOG 207 [59] and the alignment of 5130 TerL sequences from [24], and then added the alignment of 823 TerL sequences from [39] to the MSA. Next, TerL protein sequences (detected by hits to TerL profiles as described above) from the *Caudoviricetes* genomes with terminal repeats representing vOTUs in this study were added to the alignment using MAFFT 7.453 with an “--add” parameter [58]. If a genome was predicted to encode multiple copies of TerL, the one with the highest number of hit TerL motifs (Figure S2) was used. If there were multiple candidates with an equal number of hit motifs, we used a single protein characterized by the maximal length of the alignment with TerL profiles (measured in HMMER envelope coordinates combined using the R package *IRanges* 2.22.2 in case of overlap). Finally, the MSA was inspected with the help of Jalview 2.11.2.2 [71], and only the sequences containing acidic residues in the following three alignment positions were preserved in the MSA: (1) the second conserved acidic residue of the TerL Walker B motif, (2) the first conserved acidic residue of the TerL nuclease motif I and (3) the conserved acidic residue of the TerL nuclease motif II (Figure S2). MSA columns with ≥ 50% gaps were excluded from consideration using R package *Bio3D* 2.4-1 [72]. MSA conservation was estimated using the *Bio3D* 2.4-1 function *conserv* with the “method = ‘similarity’, sub.matrix = ‘blosum62’” parameters.

### Building the phylogenetic tree

The phylogenetic tree was reconstructed based on the TerL MSA using IQ-TREE 2.0.3, 1,000 replicates of ultrafast bootstrap, WAG amino acid replacement matrix [73-75]. The tree was midpoint-rooted using the R package *phangorn* 2.5.5 [76].

### Virome stability estimation

Bray-Curtis dissimilarities between samples were calculated using the function *vegdist* from the R package *vegan* 2.5-7. Data about relative abundance of the CGTR1899 vOTUs in 338 corresponding LLD and LLD follow-up samples were utilized. vOTUs that were not detected in any sample and samples without detected vOTUs were excluded from consideration. The significance of the difference between intra- and inter-individual Bray-Curtis dissimilarities was assessed using a permutation test with 10,000 iterations [17]. On each iteration, real Bray-Curtis dissimilarity values were randomly reassigned between pairs of samples, and a Wilcoxon signed-rank test comparing intra- and inter-individual dissimilarities was performed using the R function *wilcox*.*test* with the “alternative = ‘two.sided’, paired = F” parameters [77]. The empirical P-value was calculated as the proportion of P-values that were obtained based on permuted data and lower than the P-value obtained based on real data.

### Prophage-based host prediction

The analysis was based on contigs from the four Dutch cohorts that were predicted to contain both host and prophage genome fragments by Cenote-Taker2. If the prophage fragment of such a contig belonged to the CGTR1899 database, the host fragment(s) of the contig ≥ 1,000 nt were extracted for the analysis. If the number of extracted host fragments per vOTU exceeded 100, we considered a subset of 100 randomly selected host fragments. Host fragments of each vOTU were compared to the bacterial and archaeal genome sequences from the NCBI “nt” database (downloaded on 23.05.2022) using BLASTN 2.12.0+ with the “-task ‘blastn’ -perc_identity 95” parameters [50, 78]. Only hits characterized by alignment length ≥ 1,000 were considered; if multiple targets produced hits with a query, the one associated with the maximal query coverage by BLASTN alignments was used for taxonomic assignment. When multiple predictions made for a vOTU were incompatible at the host phylum level, they were disregarded, but this only occurred in one case.

### CRISPR-based host prediction

The database of CRISPR spacers published in [79] and the CRISPR-Cas++ spacers database (21.01.2021) [80] were independently compared to the representatives of the CGTR1899 vOTUs using BLASTN 2.12.0+ with the parameters “-task blastn-short -dust no -evalue 1 - max_target_seqs 1000000” [50]. A phage genome was linked to a host if there was a spacer– protospacer match characterized by ≥ 95% identity over the length of the spacer, or multiple spacer–protospacer matches characterized by ≥ 80% identity over the length of each spacer [81]. If multiple spacers of a host matched exactly the same region of a phage genome, or if multiple regions of a phage genome matched the same spacer of a host, a single spacer– protospacer match characterized by the highest bit-score was considered. Host taxonomy was retrieved from GenBank using the EFetch utility [78]. When multiple predictions made for a vOTU were incompatible at host phylum level, they were disregarded (observed in 10 cases).

### Co-abundance-based host prediction

The relative abundance of microbial taxa was estimated using MetaPhlAn 3.0.7 [82]. Correlations between the relative abundances of microbial taxa (from kingdoms to species) and CGTR54 vOTUs were assessed using the R function *cor*.*test* with the “method = ‘spearman’” parameter [77] for each cohort independently. Only taxa present in > 10 samples in a given cohort were considered. Meta-analysis of the results obtained for the independent LLD, 300OB and IBD cohorts was conducted using the R package *meta* 5.1-1 [83], function *metacor* with the “sm = ‘ZCOR’, method.tau = ‘SJ’” parameters. Multiple testing correction was performed by the R function *p*.*adjust* using the Benjamini-Hochberg procedure [84]. The host of each vOTU was predicted based on a correlation characterized by the minimal FDR obtained for this vOTU in meta-analysis.

### Finding similar extensively characterized phages

To identify extensively characterized genome sequences similar to the CGTR54 vOTU representatives, each of the 54 vOTU representatives was compared to the viral genome sequences from the NCBI “nt” database (downloaded on 23.05.2022) using BLASTN 2.12.0+ with the “-task ‘blastn’ -evalue 0.001 -perc_identity 50” parameters [50, 78]. Only query–target pairs characterized by ≥ 10% query and ≥ 50% target length coverage by the query–target BLASTN alignments were considered. Coverage was calculated with the help of R package *IRanges* 2.22.2 [56]. Information about the publications associated with the target sequences was obtained using the EFetch utility [78]. Sequence similarity within the query–target pairs was visualized as dot plots. The underlying matching words data were generated using the EMBOSS 6.6.0 function *polydot* with the “-wordsize 12” parameter [85].

### Associations with human phenotypes

Association analysis was carried out for 1135 LLD cohort samples, 207 phenotypes (missing values imputed) and the CGTR54 vOTUs. Separately, prevalence of the 54 vOTUs was compared between the following groups: (1) LLD vs. IBD, (2) LLD vs. 300OB, (3) within the IBD cohort: CD vs. UC, (4) within the IBD cohort: exclusively colonic vs. ileal-inclusive disease location and (5) within the 300OB cohort: absence vs. presence of metabolic syndrome. All analyses were conducted using logistic regression adjusted for (1) the age and sex of the cohort participants and (2) the age and sex of the participants and the log-transformed abundance of the host predicted as described above. The corresponding phenotype was used as a predictor, and the detection of a phage was an outcome in the logistic regression. Logistic regression was fitted using the R 4.0.3 function *glm* with the “family = ‘binomial’” parameter [77]. Multiple testing correction was conducted using the R function *p*.*adjust* employing the Benjamini-Hochberg procedure [84]. A significance threshold of FDR < 0.05 was used.

### Visualization

The Sankey diagram was prepared using R package *alluvial* v0.1-2. Sequence logos were constructed using R package *ggseqlogo* 0.1 [86]. The phylogenetic tree was visualized using R package *ape* 5.4-1 [87]. Boxplots were plotted using R package *vioplot* 0.3.7. Colors designating host phyla were selected using R package *RColorBrewer* 1.1-2. Genome annotation labels in Material S1 were positioned with the help of R package *TeachingDemos* 2.12. RT MSA was visualized using ESPript 3.0 [88].

## Data and code availability

The 17,193 phage genomes and genome fragments identified in this study and belonging to the CGTR1899, as well as sequence of the contig NL_vir005341, will be published in a Figshare repository and are currently available from the corresponding authors. The code used to conduct the analysis was deposited to the GitHub repository https://github.com/aag1/NL_vir_analysis/.

## Acknowledgments

We would like to thank Kate Mc Intyre for editing the manuscript, the Center for Information Technology of the University of Groningen for their support and for providing access to the Peregrine high performance computing cluster and the volunteers of the LifeLines-DEEP, 300OB and IBD cohorts for their participation. S.G. holds a scholarship from the Graduate School of Medical Sciences, University of Groningen. A.Z. is supported by European Research Council (ERC) Starting Grant 715772, Netherlands Organization for Scientific Research (NWO) VIDI grant 016.178.056 and NWO Gravitation grant ExposomeNL 024.004.017. J.F. is supported by NWO Gravitation grant Netherlands Organ-on-Chip Initiative 024.003.001, ERC Consolidator grant 101001678 and NWO VICI grant VI.C.202.022. N.P.R., M.G.N., J.F. and A.Z. are supported by the Netherlands Heart Foundation CVON grant 2018-27. R.K.W. is supported by the Seerave Foundation and the Dutch Digestive Foundation (16-14).

## Author contributions

A.G. designed the study, performed data analysis and wrote the paper. A.V.V., N.P.R., M.G.N., R.K.W., J.F. and A.Z. provided the data. A.G., S.G., A.K. and A.Z. discussed the analysis. All authors reviewed and edited the manuscript.

**Figure S1. Workflow leading to delineation of the CGTR1899 and CGTR54 databases**.

**Figure S2. TerL alignment conservation**. Mean conservation of the multiple sequence alignment (MSA) of TerL proteins encoded by the CGTR1899 vOTU representatives (Table S2) was calculated using an 11-column window sliding along the MSA with a 1-column step. MSA columns with ≥ 50% gaps were excluded from consideration. Adenosine triphosphatase motifs Walker A and B and nuclease motifs I, II and III are each highlighted by a distinct color. The sequence logo of each motif is presented below the conservation profile, and the color of the frame designates the motif.

**Figure S3. Sequence similarity between genomes representing the CGTR54 vOTUs and extensively characterized phages**. Each dot plot illustrates similarity between a pair of sequences, X-axis corresponds to a sequence from the literature, Y-axis corresponds to a CGTR54 sequence. Coordinates are indicated in kilobases. Every 12-letter word shared by a pair of sequences is presented as a black dot on a dot plot. If a reverse complement of a sequence was analyzed, the letters “RC” are added to the sequence identifier. Contig length and coverage are omitted from the sequence identifiers for brevity, where applicable. The color of the dot plot frame points to a publication about the X-axis phage: blue, *Minot et al. 2012*; green, *Ly et al. 2016*; pink, *Dzunkova et al. 2019*. See Table S4.

**Table S1. Properties of the CGTR1899 phage genome sequences**. vOTUs are ordered according to their position on the TerL-based phylogenetic tree (Figure 3). Properties of the phage genome sequences identified in the form of prophage contigs were assessed after the fragments of microbial genomes were cleaved off.

**Table S2. Properties of the CGTR1899 vOTU representatives**. TerL gene coordinates are specified by indicating the genome strand (f, forward or r, reverse) and coordinates separated by semicolons. The number of positive samples is indicated for the four Dutch cohorts and for all and healthy adult (HA) Danish fecal viromes (DFV).

**Table S3. Predicted phage hosts**. First page, results of the prophage-based host prediction conducted for the CGTR1899 database. Second page, results of the CRISPR-based host prediction conducted for the representatives of the CGTR1899 vOTUs. Third page, associations between the relative abundances of the CGTR54 vOTUs and microbial taxa. Microbial taxa are designated using the MetaPhlAn software format, with “k_”, “p_”, “c_”, “o_”, “f_”, “g_” and “s_” standing for kingdom, phylum, class, order, family, genus and species, respectively. Fourth page, comparison between host predictions made by the different methods. Data are shown for those CGTR54 vOTUs for which predictions by multiple methods were available. Prophage-, CRISPR- and co-abundance-based predictions are shown with blue, orange and white backgrounds, respectively. Co-abundance-based predictions that diverge from other predictions at phylum level are highlighted by red font.

**Table S4. Experimentally characterized phage sequences similar to the CGTR54 vOTU representatives**.

**Table S5. Associations with human phenotypes**. First page, associations between the 207 LLD cohort phenotypes and prevalence of the CGTR54 vOTUs. Second page, definition of the 207 LLD cohort phenotypes considered in the analysis. Subsequent pages, results of the analyses comparing the prevalence of the 54 vOTUs between the following groups: (1) LLD vs. 300OB cohort, (2) LLD vs. IBD cohort, (3) within 300OB cohort: absence vs. presence of metabolic syndrome, (4) within IBD cohort: CD vs. UC, and (5) within IBD cohort: exclusively colonic vs. ileal-inclusive disease location. Significant associations (FDR < 0.05) are highlighted by green background.

**Test S1. Benchmarking of virus detection and taxonomic assignment**.

**Material S1. Characteristics of the CGTR54 genomes**. Each page corresponds to a genome representing a CGTR54 vOTU. Genome name, predicted genetic code and type of terminal repeats are indicated at the top of the page. *Top panel*, coverage by reads from the four Dutch cohorts: each line corresponds to a sample and represents mean coverage depth in a sliding window. The color of the line indicates cohort. *Second panel from top*, genome map. Genome is represented by a white bar in black frame. Three forward and three reverse frames are shown. ORFs are represented by light gray bars in black frames. Red frame denotes the TerL ORF key to recognizing the genome as belonging to the class *Caudoviricetes*. Regions of an ORF matching the PFAM profile used for its annotation are indicated by color: blue – structural protein profiles and profiles of proteins implicated in assembly of virus particles, pink – DNA polymerase profiles, green – integrase profiles, orange – reverse transcriptase profiles, and dark gray – all other profiles. Names of the profiles are indicated above the genome map (note that in some cases subdomains justifying the name of the profile may not be part of the alignment between the profile and the ORF product). Predicted tRNA genes are shown by dark red bars, and their names include the tRNA isotype and anticodon. Position of nucleotide repeats is indicated by vertical orange lines below the genome map, with pairs of repeats connected by horizontal orange lines. If one of the two repeats lies in the reverse strand, the orange line is dashed. *Third panel from top*, average content of the four nucleotides along the genome. *Bottom panels*, regular and cumulative GC- and AT-skew. All curves were generated using a 1,001 nt window sliding with a 200 nt step.

**Material S2. MSA of RTs from the CGTR54 genomes representing vOTUs**. Absolutely conserved residues are shown on red background and partially conserved residues in red font. Conserved motifs are underlined in green. Their names are indicated underneath. Name of each protein sequence consists of a genome name followed by strand (f, forward or r, reverse) and coordinates of the RT gene separated by semicolons. Contig length and coverage are omitted from the genome names for brevity, where applicable.

