## Supplementary figures and images for "Diversity and ecology of *Caudoviricetes* phages with genome terminal repeats in fecal metagenomes from four Dutch cohorts"

### Figure S1

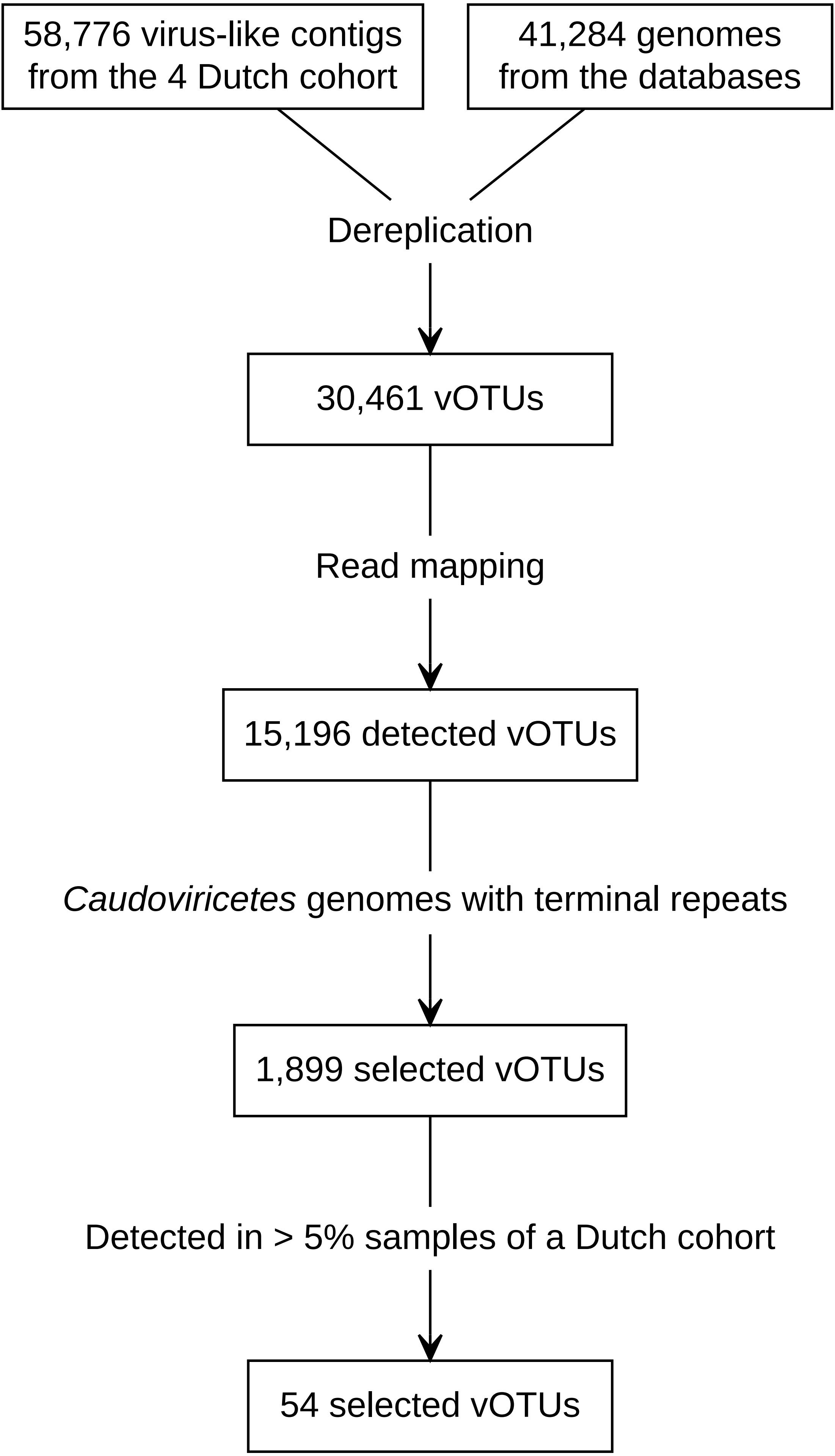

### Figure S2

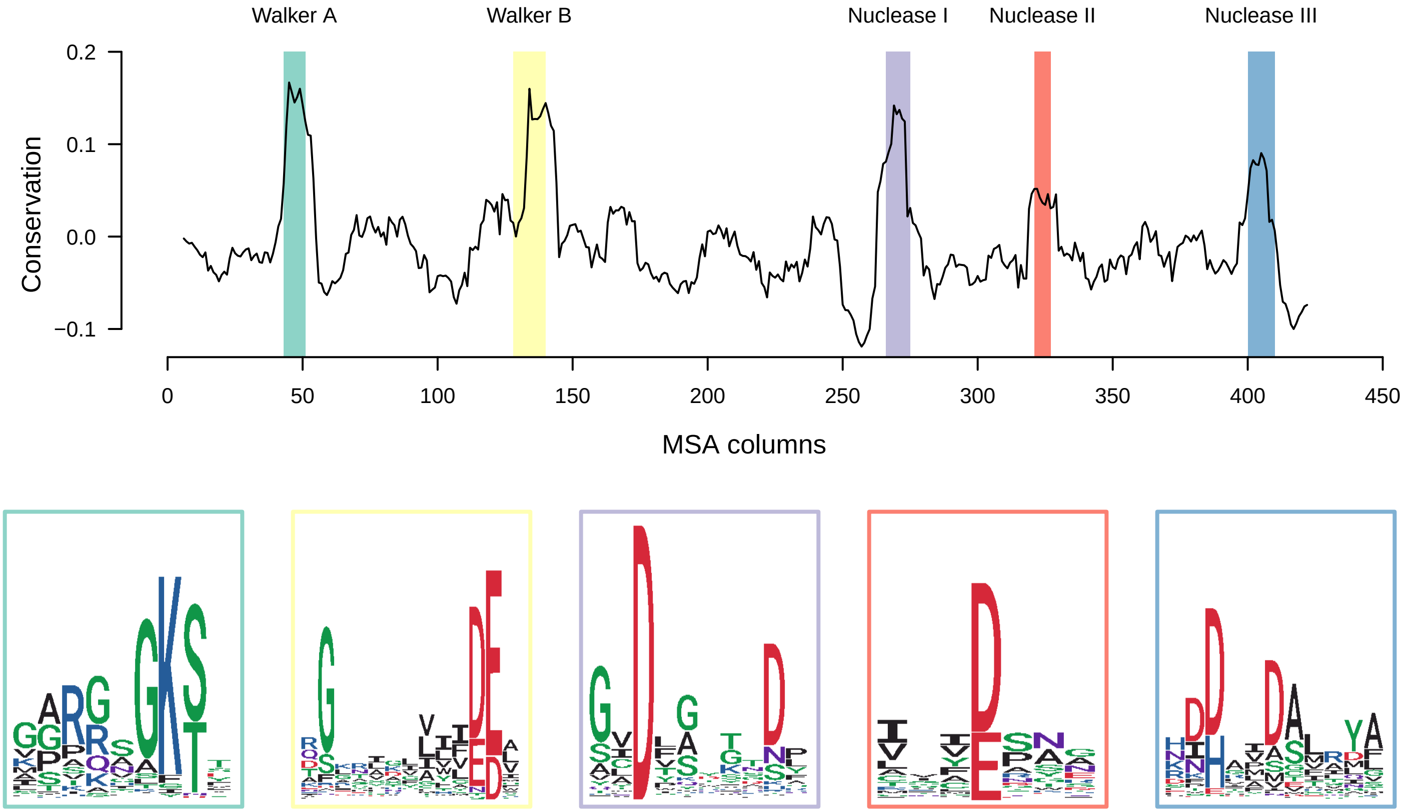

### Figure S3

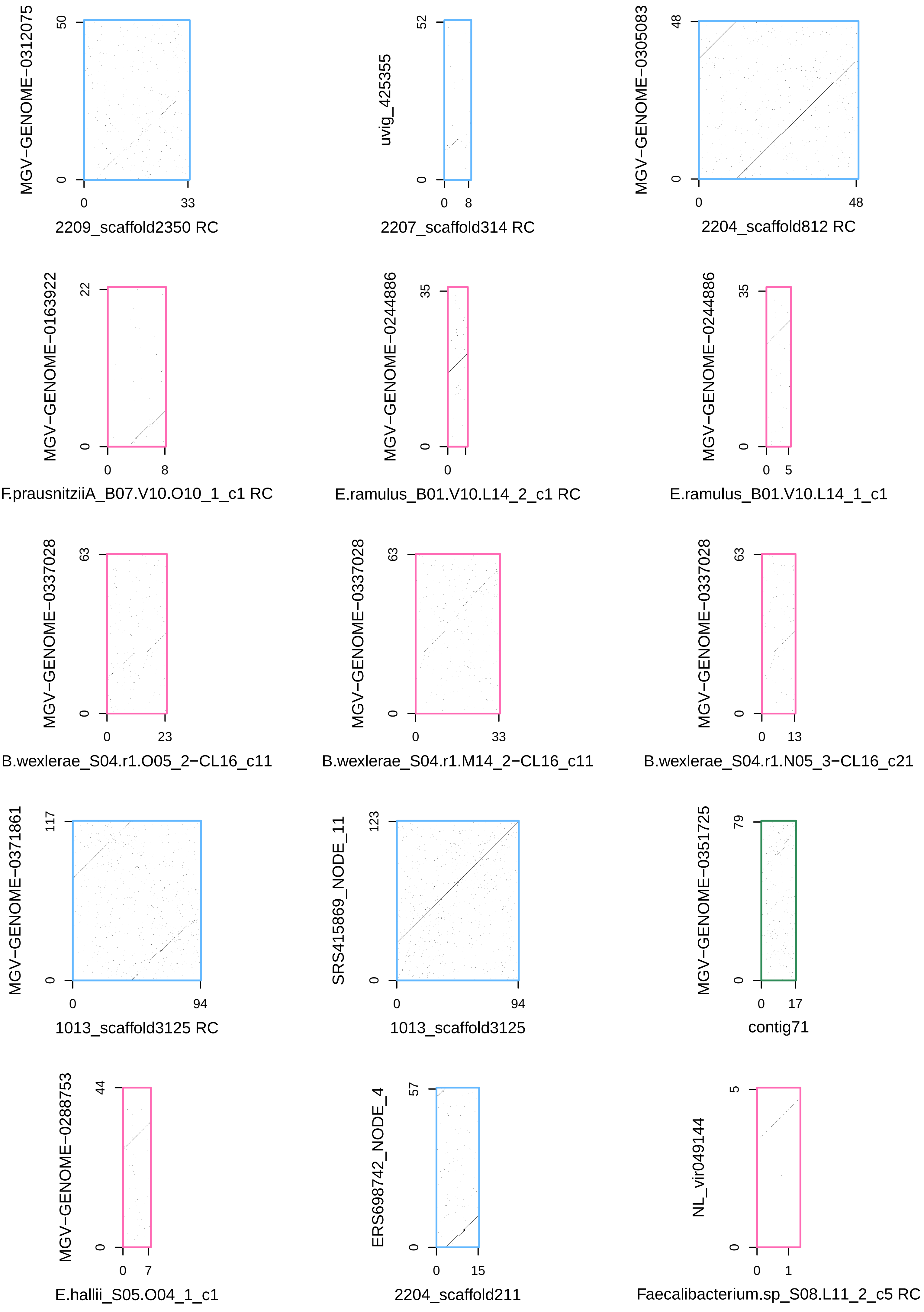
