## Supplementary material for "Diversity and ecology of *Caudoviricetes* phages with genome terminal repeats in fecal metagenomes from four Dutch cohorts": Material S1

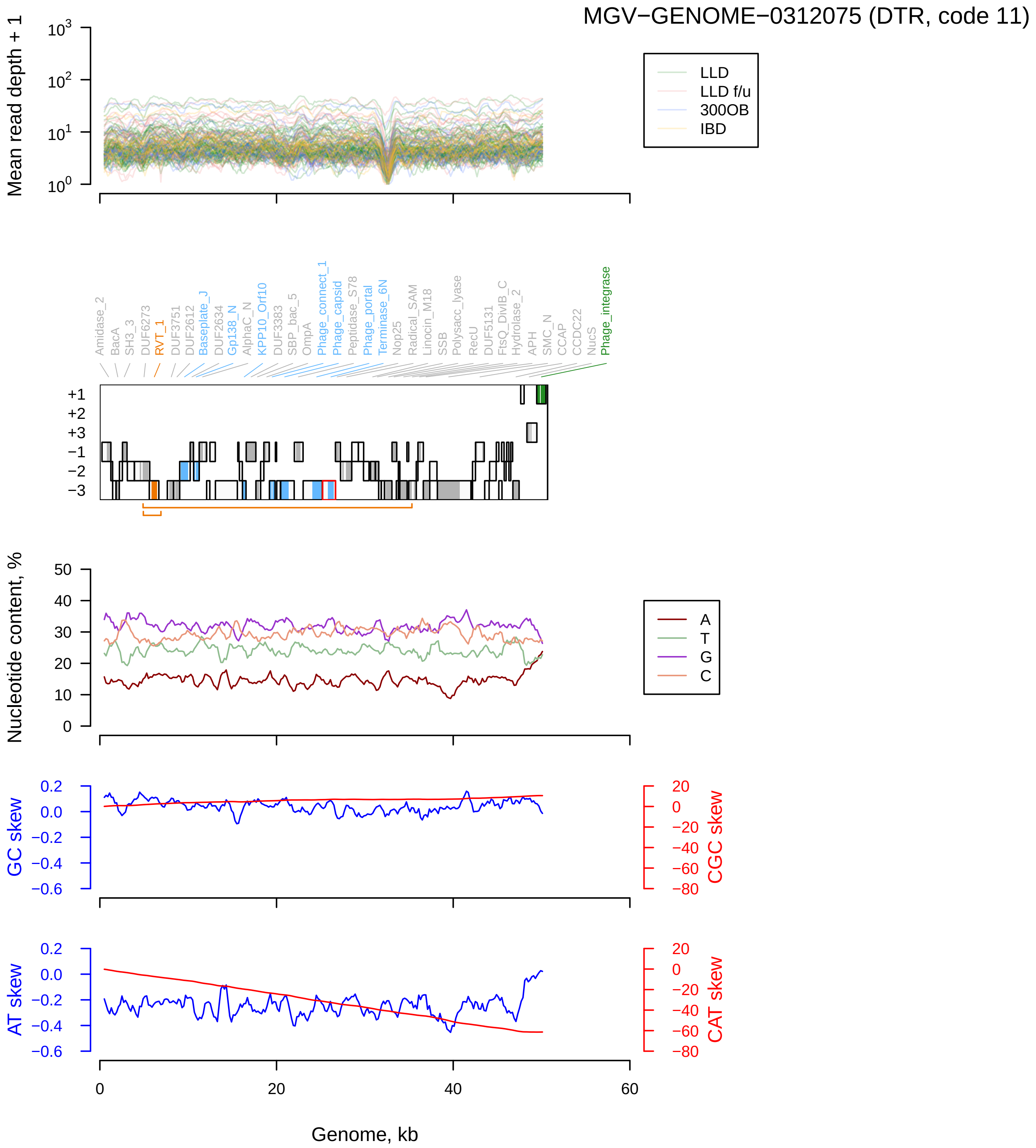

MGV−GENOME−0312075 (DTR, code 11)
Mean read depth + 1
$10^0$
$10^1$
$10^2$
$10^3$
LLD
LLD f/u
300OB
IBD
Amidase_2
BacA
SH3_3
DUF6273
RVT_1
DUF3751
DUF2612
Baseplate_J
DUF2634
Gp138_N
AlphaC_N
KPP10_Orf10
DUF3383
SBP_bac_5
OmpA
Phage_connect_1
Phage_capsid
Peptidase_S78
Phage_portal
Terminase_6N
Nop25
Radical_SAM
Linocin_M18
SSB
Polysacc_lyase
RecU
DUF5131
FtsQ_DivIB_C
Hydrolase_2
APH
SMC_N
CCAP
CCDC22
NucS
Phage_integrase
+1
+2
+3
−1
−2
−3
Nucleotide content, %
0
10
20
30
40
50
A
T
G
C
GC skew
0.2
0.0
−0.2
−0.4
−0.6
CGC skew
20
0
−20
−40
−60
−80
AT skew
CAT skew
0
20
40
60
Genome, kb

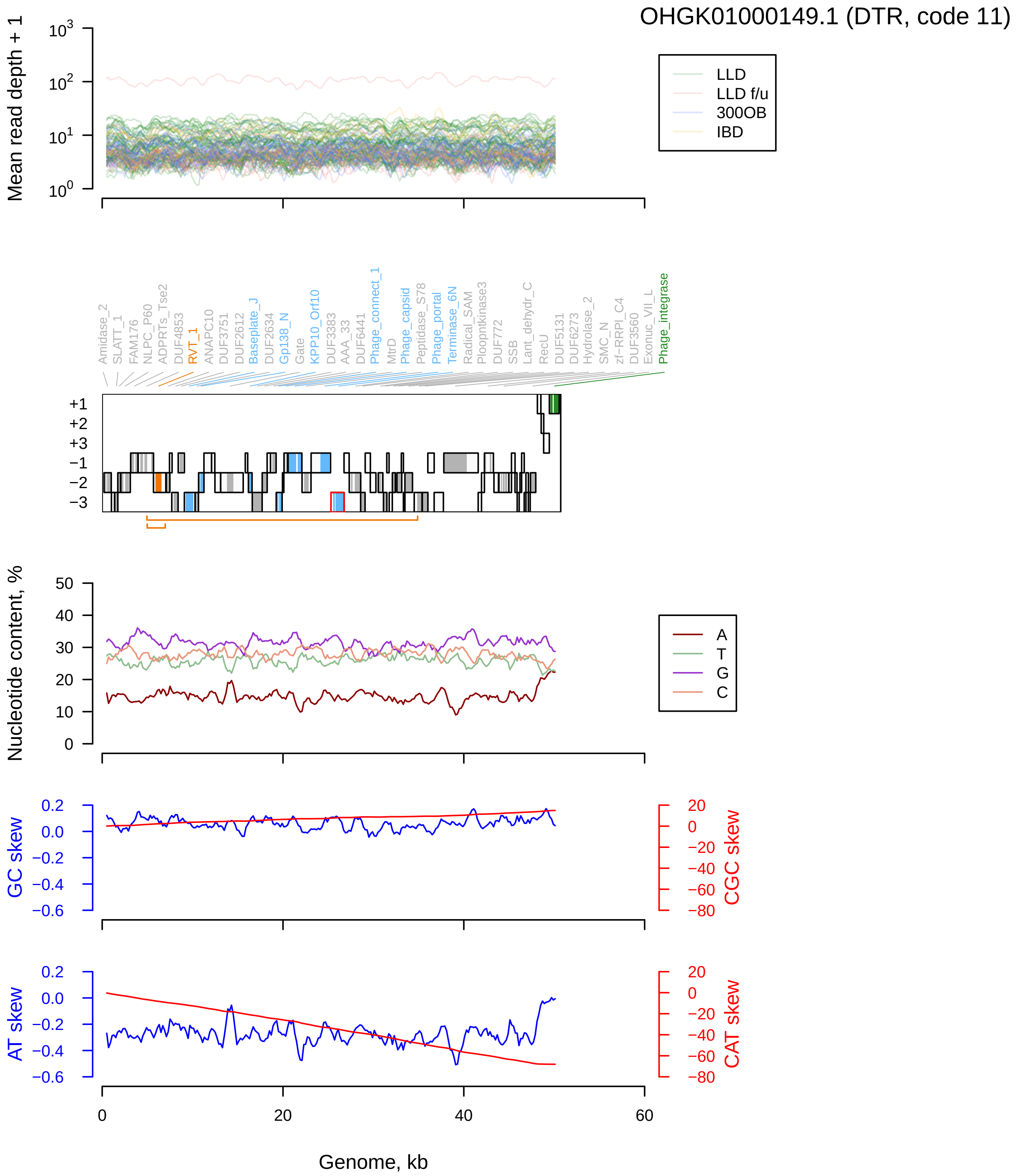

OHGK01000149.1 (DTR, code 11)
Mean read depth + 1
LLD
LLD f/u
300OB
IBD
Amidase_2
SLATT_1
FAM176
NLPC_P60
ADPRTs_Tse2
DUF4853
RVT_1
ANAPC10
DUF3751
DUF2612
Baseplate_J
DUF2634
Gp138_N
Gate
KPP10_Orf10
DUF3383
AAA_33
DUF6441
Phage_connect_1
MtrD
Phage_capsid
Peptidase_S78
Phage_portal
Terminase_6N
Radical_SAM
Ploopntkinase3
DUF772
SSB
Lant_dehydr_C
RecU
DUF5131
DUF6273
Hydrolase_2
SMC_N
zf-RRPI_C4
DUF3560
Exonuc_VII_L
Phage_integrase
+1
+2
+3
−1
−2
−3
Nucleotide content, %
A
T
G
C
GC skew
CGC skew
AT skew
CAT skew
Genome, kb

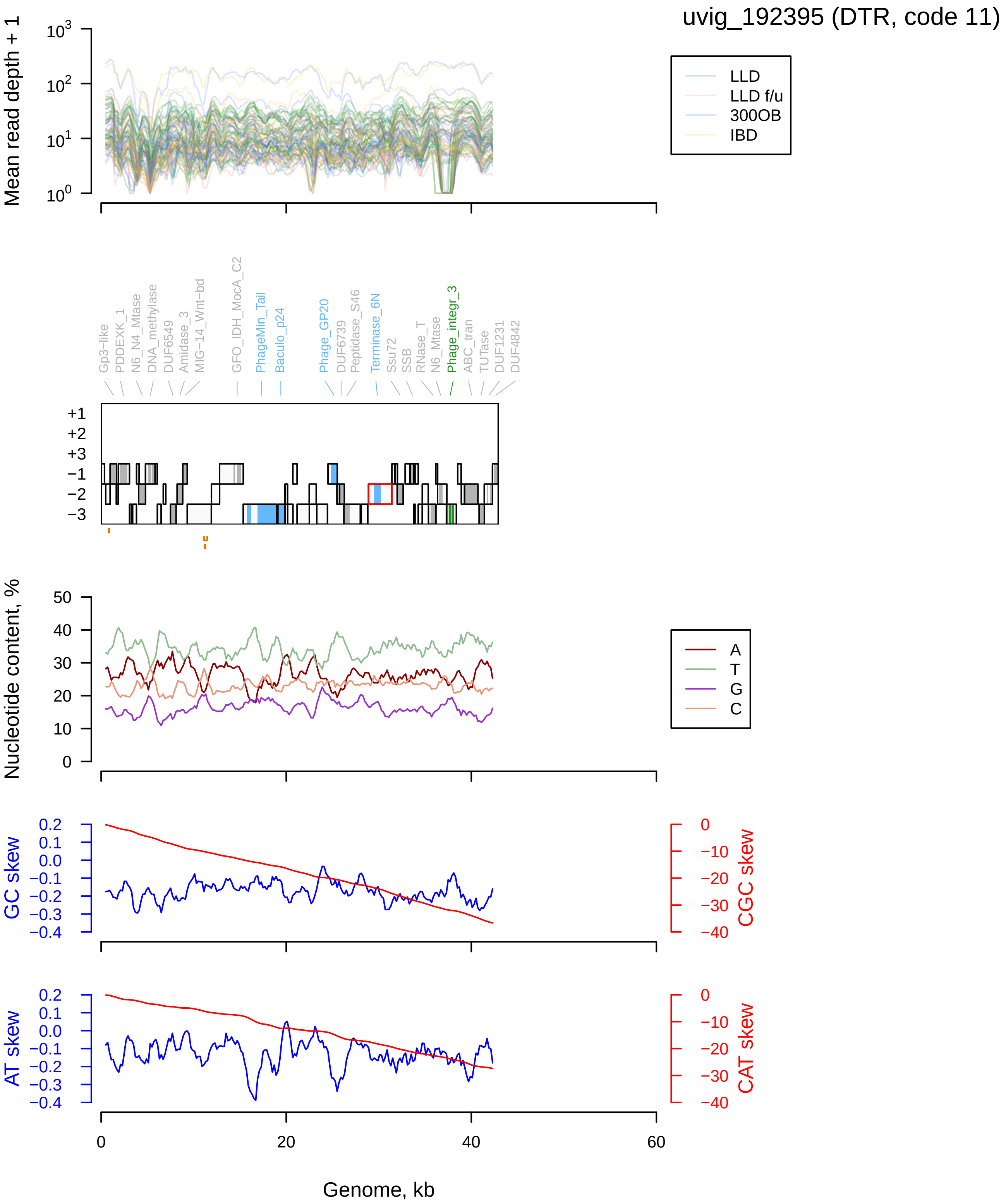
uvig_192395 (DTR, code 11)
Mean read depth + 1
LLD
LLD f/u
300OB
IBD
Gp3−like
PDDEXK_1
N6_N4_Mtase
DNA_methylase
DUF6549
Amidase_3
MIG−14_Wnt−bd
GFO_IDH_MocA_C2
PhageMin_Tail
Baculo_p24
Phage_GP20
DUF6739
Peptidase_S46
Terminase_6N
Ssu72
SSB
RNase_T
N6_Mtase
Phage_integr_3
ABC_tran
TUTase
DUF1231
DUF4842
Nucleotide content, %
A
T
G
C
GC skew
CGC skew
AT skew
CAT skew
Genome, kb

### MGV−GENOME−0278294 (DTR, code 11)

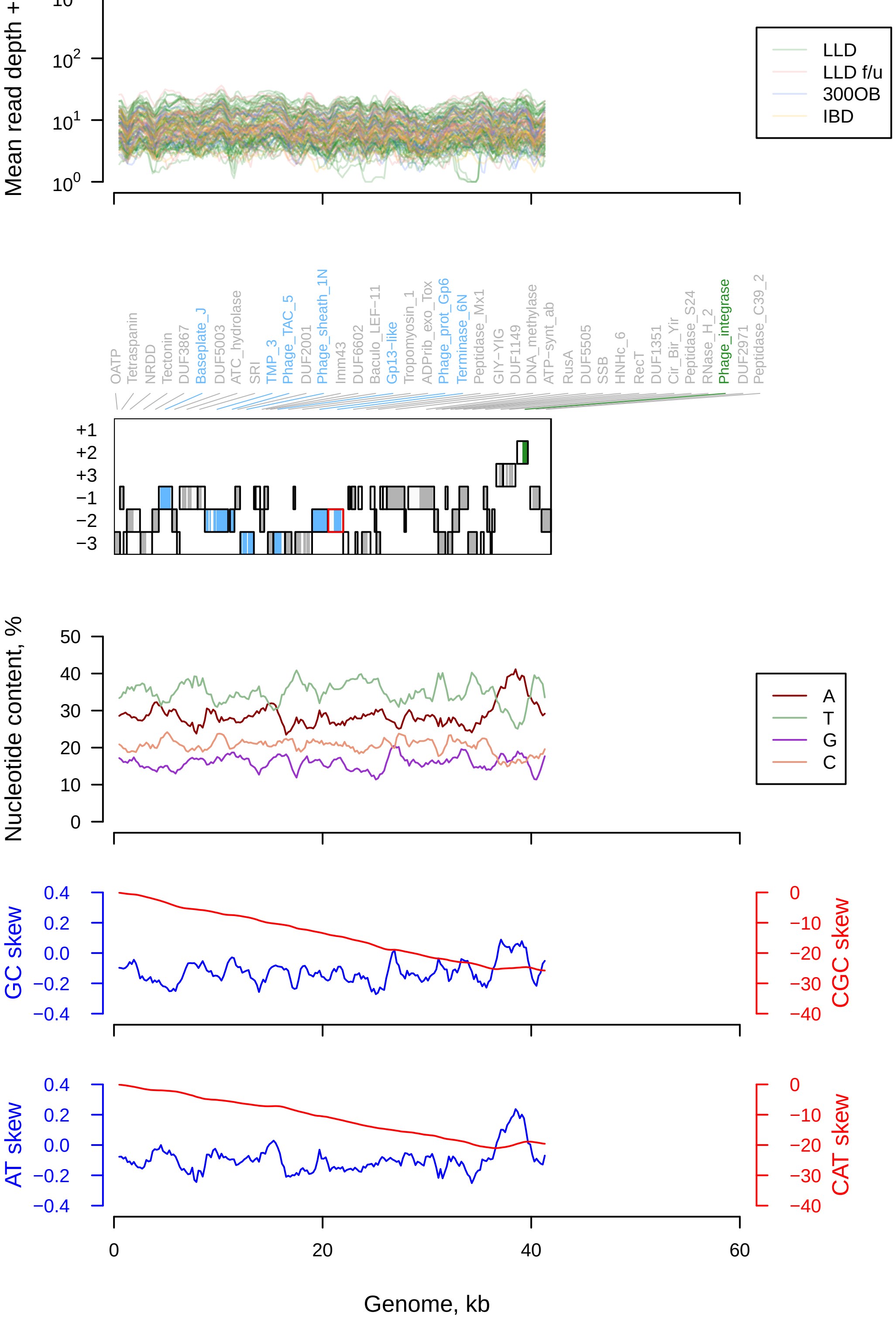

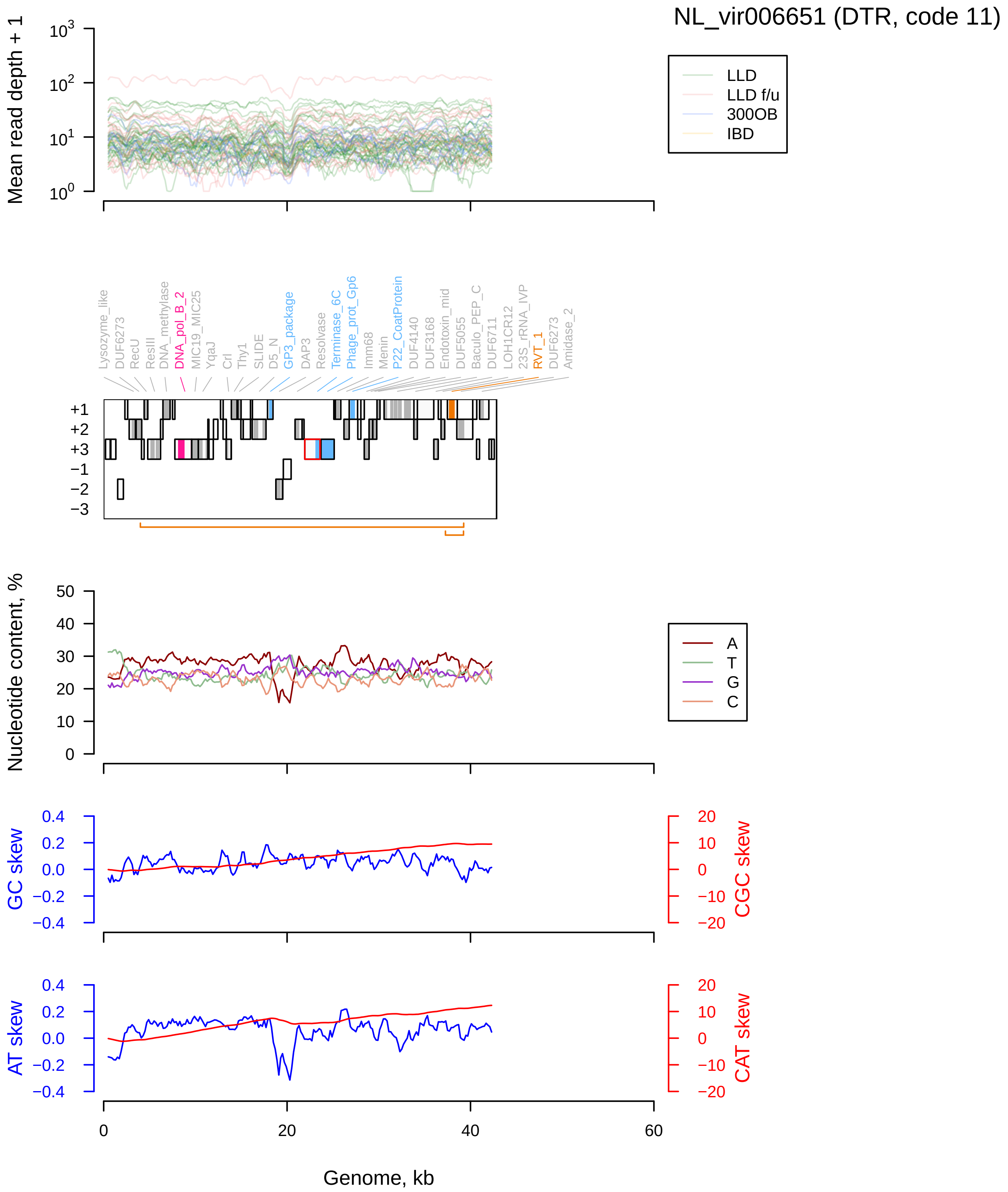

NL_vir006651 (DTR, code 11)
Mean read depth + 1
LLD
LLD f/u
300OB
IBD
Lysozyme_like
DUF6273
RecU
ResIII
DNA_methylase
DNA_pol_B_2
MIC19_MIC25
YqaJ
Crl
Thy1
SLIDE
D5_N
GP3_package
DAP3
Resolvase
Terminase_6C
Phage_prot_Gp6
Imm68
Menin
P22_CoatProtein
DUF4140
DUF3168
Endotoxin_mid
DUF5055
Baculo_PEP_C
DUF6711
LOH1CR12
23S_rRNA_IVP
RVT_1
DUF6273
Amidase_2
Nucleotide content, %
A
T
G
C
GC skew
CGC skew
AT skew
CAT skew
Genome, kb

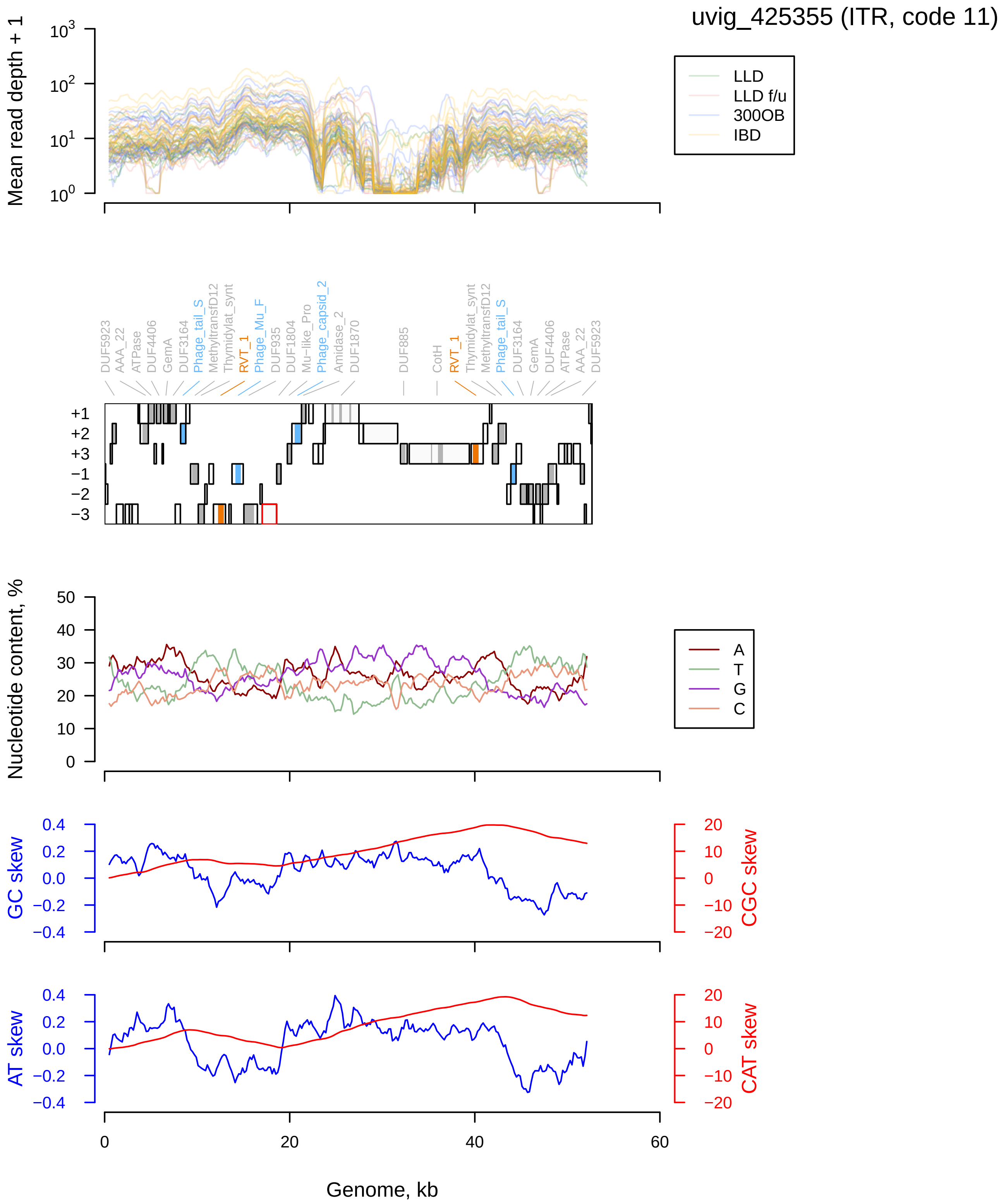

uvig_425355 (ITR, code 11)
Mean read depth + 1
LLD
LLD f/u
300OB
IBD
DUF5923
AAA_22
ATPase
DUF4406
GemA
DUF3164
Phage_tail_S
MethyltransfD12
Thymidylat_synt
RVT_1
Phage_Mu_F
DUF935
DUF1804
Mu-like_Pro
Phage_capsid_2
Amidase_2
DUF1870
DUF885
CotH
Nucleotide content, %
A
T
G
C
GC skew
CGC skew
AT skew
CAT skew
Genome, kb

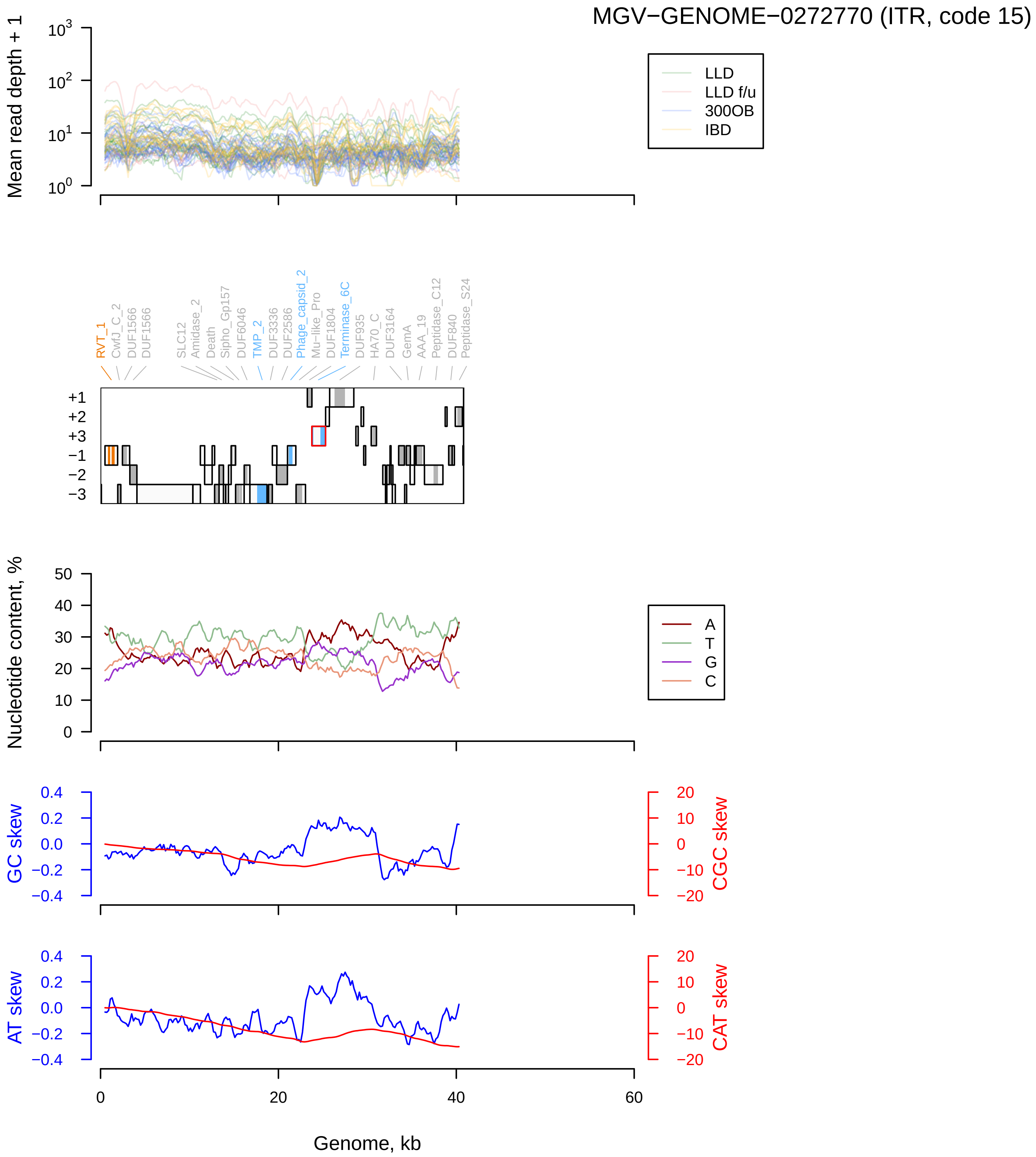
MGV−GENOME−0272770 (ITR, code 15)
Mean read depth + 1
LLD
LLD f/u
300OB
IBD
RVT_1
CwfJ_C_2
DUF1566
DUF1566
SLC12
Amidase_2
Death
Sipho_Gp157
DUF6046
TMP_2
DUF3336
DUF2586
Phage_capsid_2
Mu−like_Pro
DUF1804
Terminase_6C
DUF935
HA70_C
DUF3164
GemA
AAA_19
Peptidase_C12
DUF840
Peptidase_S24
Nucleotide content, %
A
T
G
C
GC skew
CGC skew
AT skew
CAT skew
Genome, kb

OLSK01002610.1 (DTR, code 11)

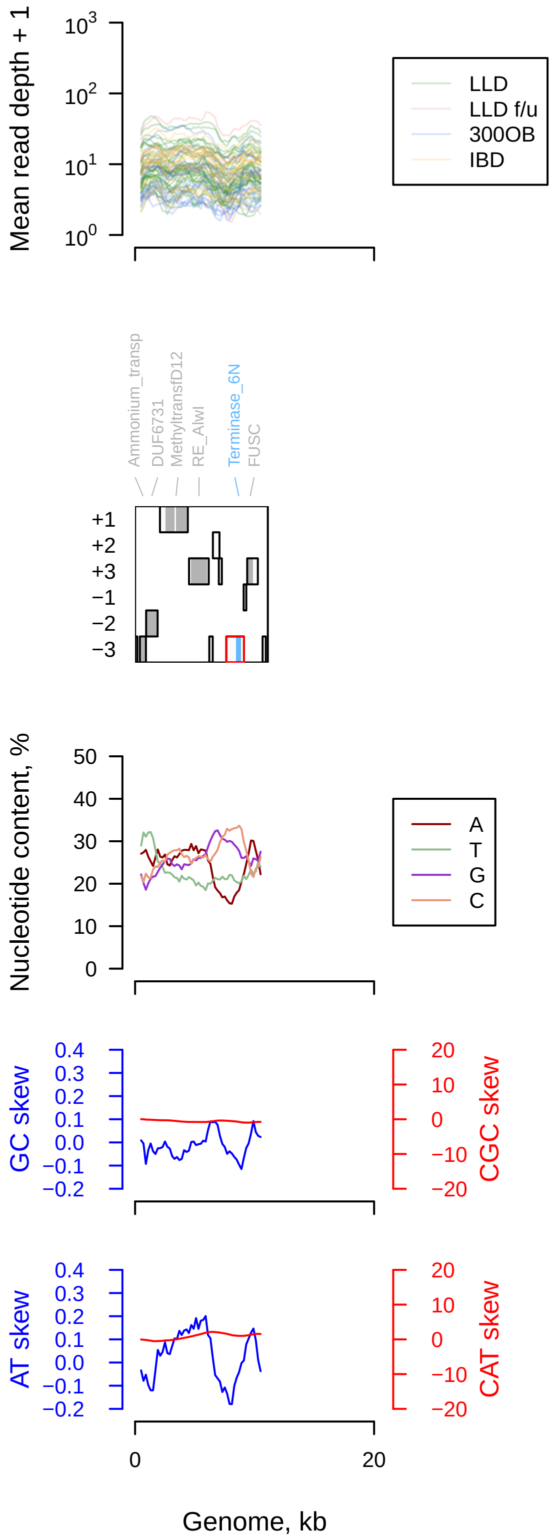

MGV−GENOME−0308888 (DTR, code 15)

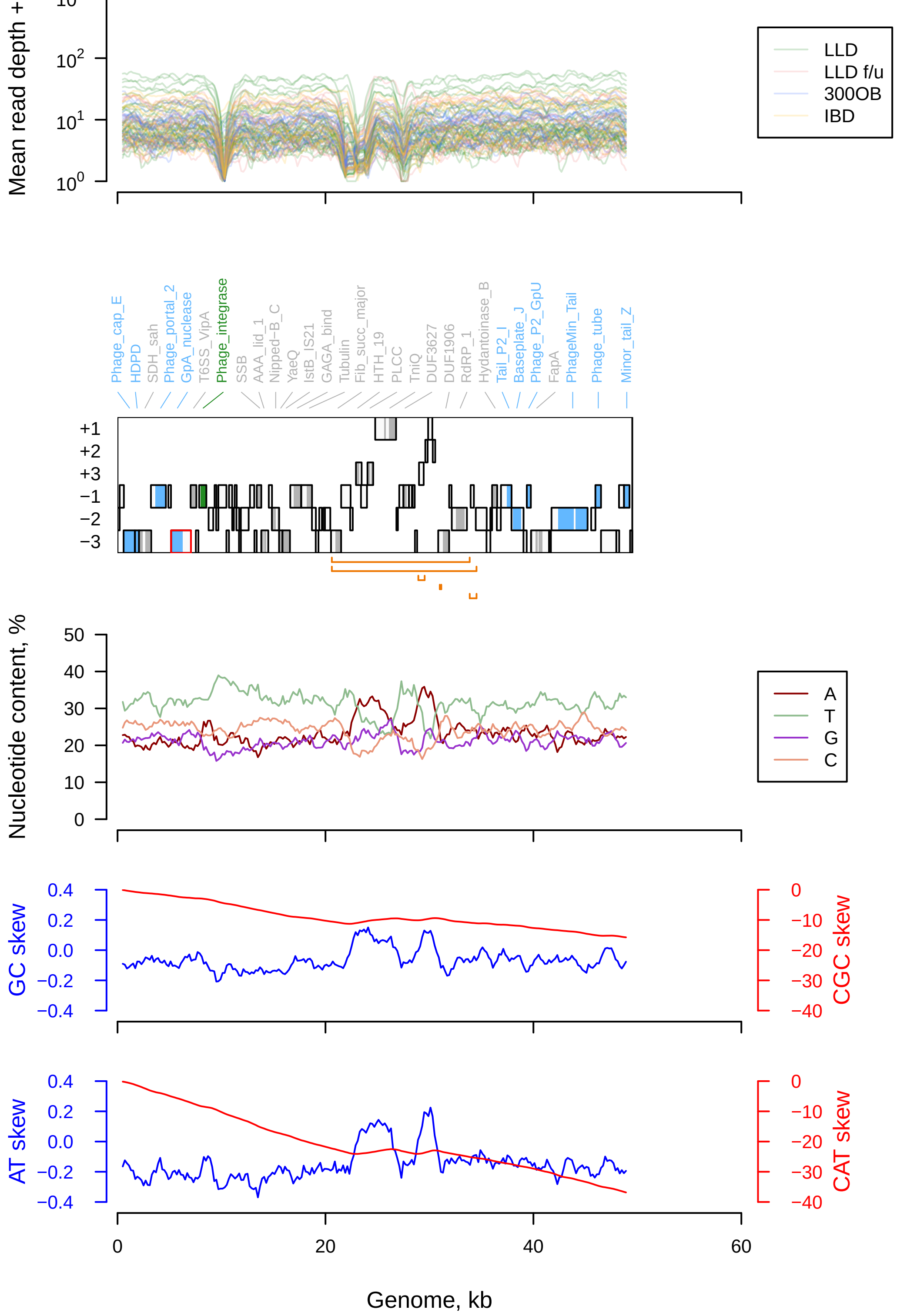

MGV−GENOME−0297536 (DTR, code 11)

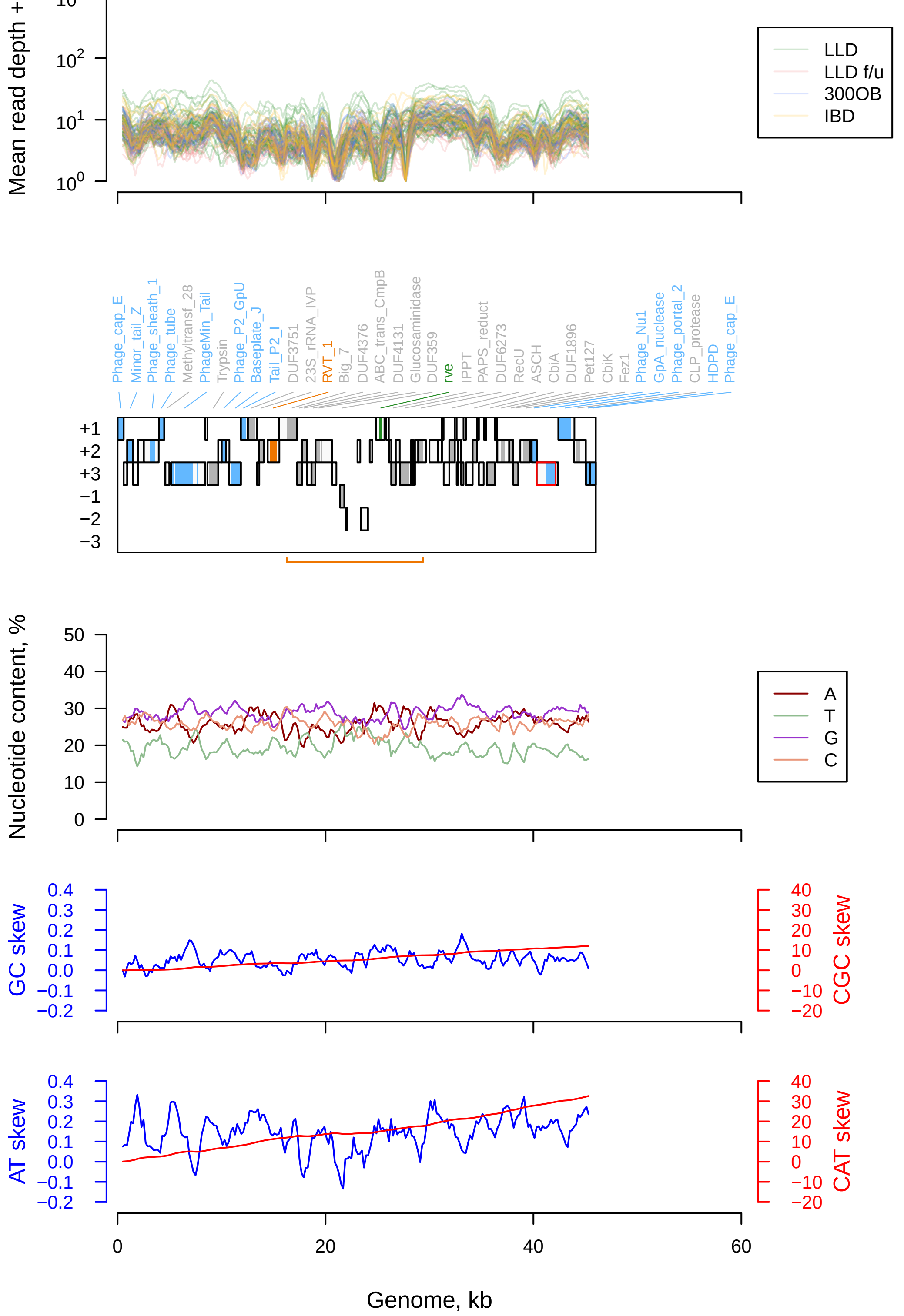

MGV−GENOME−0310879 (DTR, code 11)

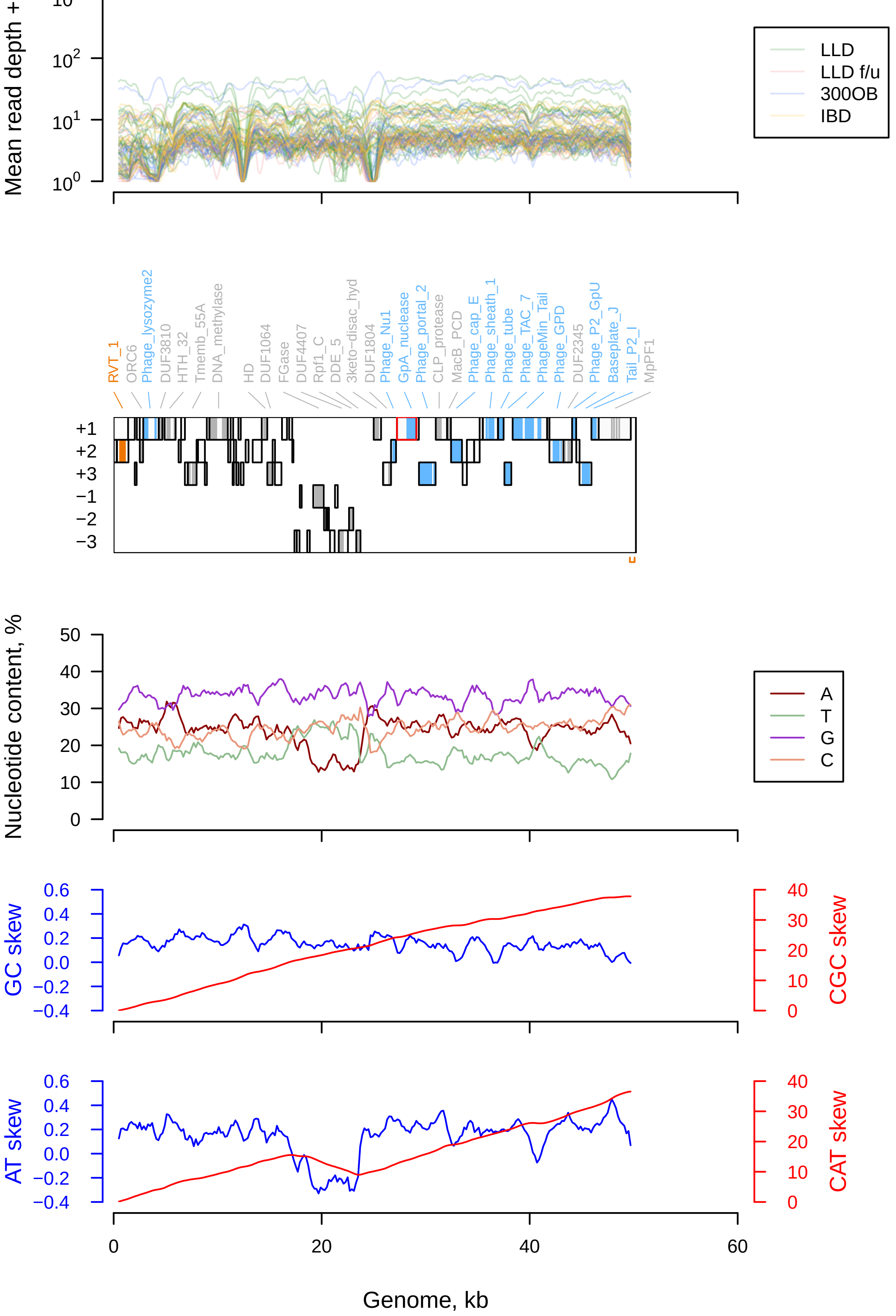

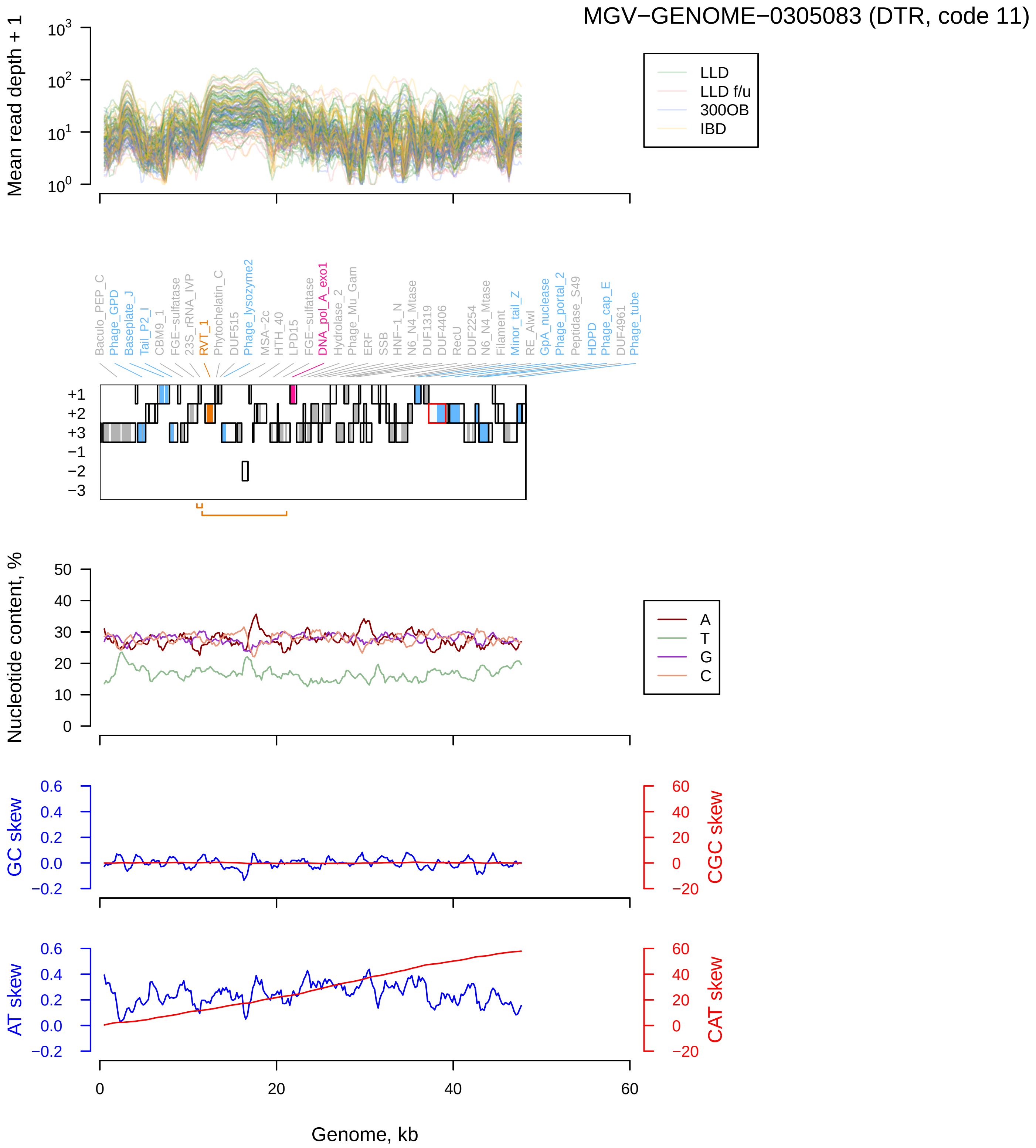

MGV−GENOME−0305083 (DTR, code 11)
Mean read depth + 1
LLD
LLD f/u
300OB
IBD
Baculo_PEP_C
Phage_GPD
Baseplate_J
Tail_P2_I
CBM9_1
FGE−sulfatase
23S_rRNA_IVP
RVT_1
Phytochelatin_C
DUF515
Phage_lysozyme2
MSA−2c
HTH_40
LPD15
FGE−sulfatase
DNA_pol_A_exo1
Hydrolase_2
Phage_Mu_Gam
ERF
SSB
HNF−1_N
N6_N4_Mtase
DUF1319
DUF4406
RecU
DUF2254
N6_N4_Mtase
Filament
Minor_tail_Z
RE_AlwI
GpA_nuclease
Phage_portal_2
Peptidase_S49
HDPD
Phage_cap_E
DUF4961
Phage_tube
Nucleotide content, %
A
T
G
C
GC skew
CGC skew
AT skew
CAT skew
Genome, kb

### OMFQ01000250.1 (DTR, code 11)

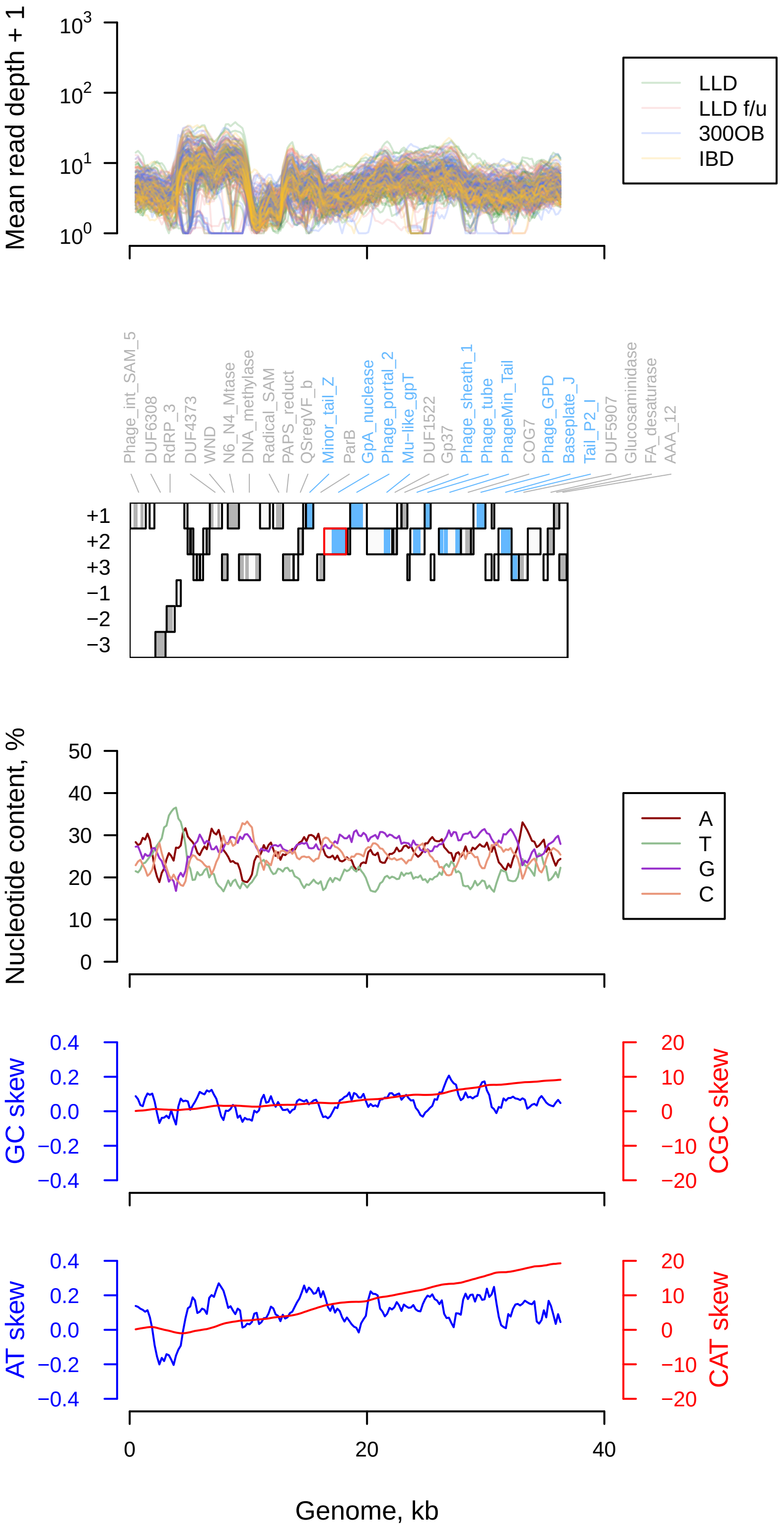

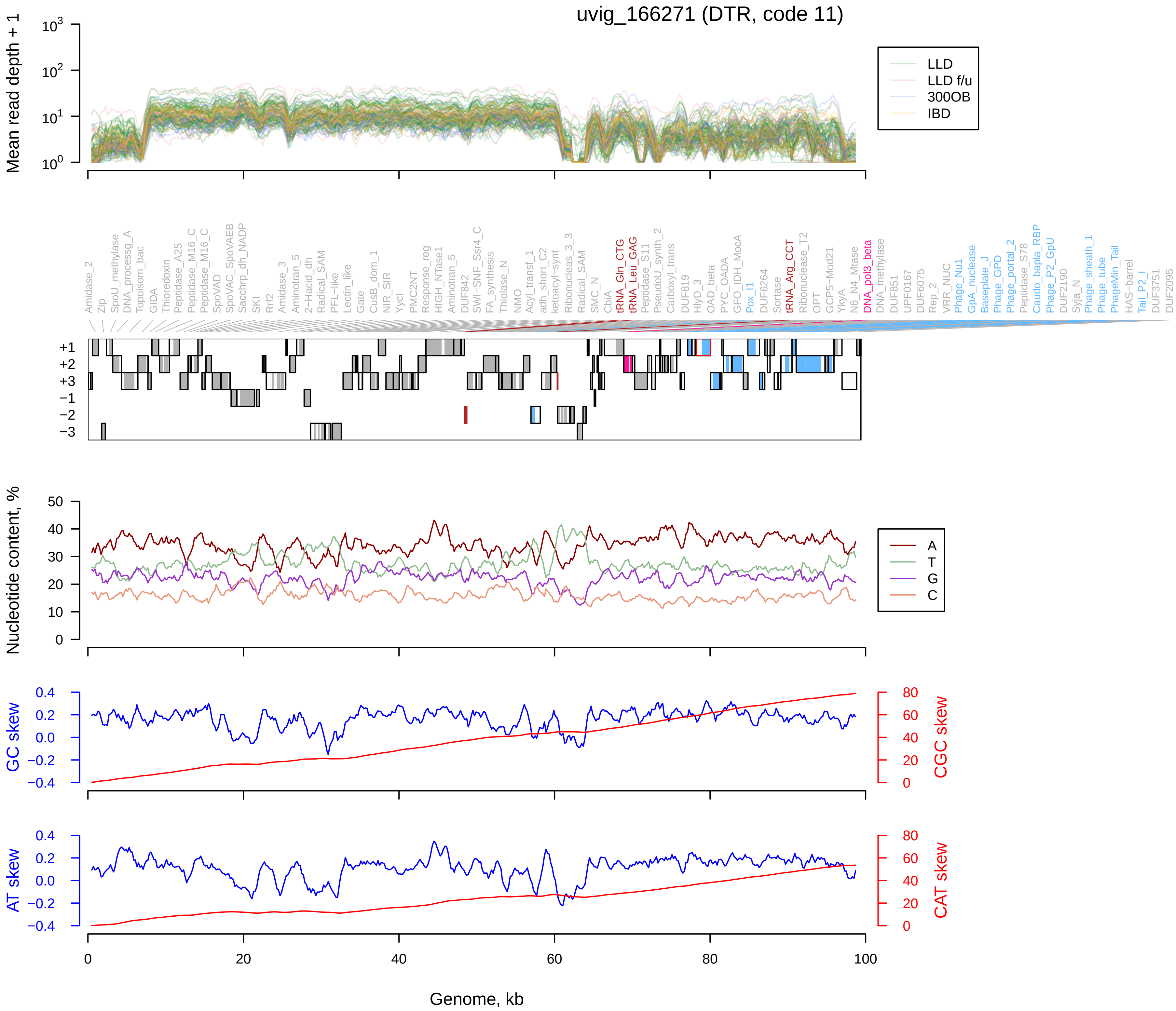

uvig_166271 (DTR, code 11)
Mean read depth + 1
LLD
LLD f/u
300OB
IBD
Amidase_2
Zip
SpoU_methylase
DNA_processg_A
Topoisom_bac
GIDA
Thioredoxin
Peptidase_A25
Peptidase_M16_C
Peptidase_M16_C
SpoVAD
SpoVAC_SpoVAEB
Sacchrp_dh_NADP
SKI
Rrf2
Amidase_3
Aminotran_5
2-Hacid_dh
Radical_SAM
PFL-like
Lectin_like
Gate
CusB_dom_1
NIR_SIR
YycI
PMC2NT
Response_reg
HIGH_NTase1
Aminotran_5
DUF842
SWI-SNF_Ssr4_C
FA_synthesis
Thiolase_N
NMO
Acyl_transf_1
adh_short_C2
ketoacyl-synt
Ribonucleas_3_3
Radical_SAM
SMC_N
CbiA
tRNA_Gln_CTG
tRNA_Leu_GAG
Peptidase_S11
PseudoU_synth_2
Carboxyl_trans
DUF819
HlyD_3
OAD_beta
PYC_OADA
GFO_IDH_MocA
Pox_I1
DUF6264
Sortase
tRNA_Arg_CCT
Ribonuclease_T2
OPT
GCP5-Mod21
YkyA
N6_N4_Mtase
DNA_pol3_beta
DNA_methylase
DUF851
UPF0167
DUF6075
Rep_2
VRR_NUC
Phage_Nu1
GpA_nuclease
Baseplate_J
Phage_GPD
Phage_portal_2
Peptidase_S78
Caudo_bapla_RBP
Phage_P2_GpU
DUF2190
Syja_N
Phage_sheath_1
Phage_tube
PhageMin_Tail
HAS-barrel
Tail_P2_I
DUF3751
DUF2095
+1
+2
+3
−1
−2
−3
Nucleotide content, %
A
T
G
C
GC skew
CGC skew
AT skew
CAT skew
Genome, kb

### MGV−GENOME−0212520 (DTR, code 11)

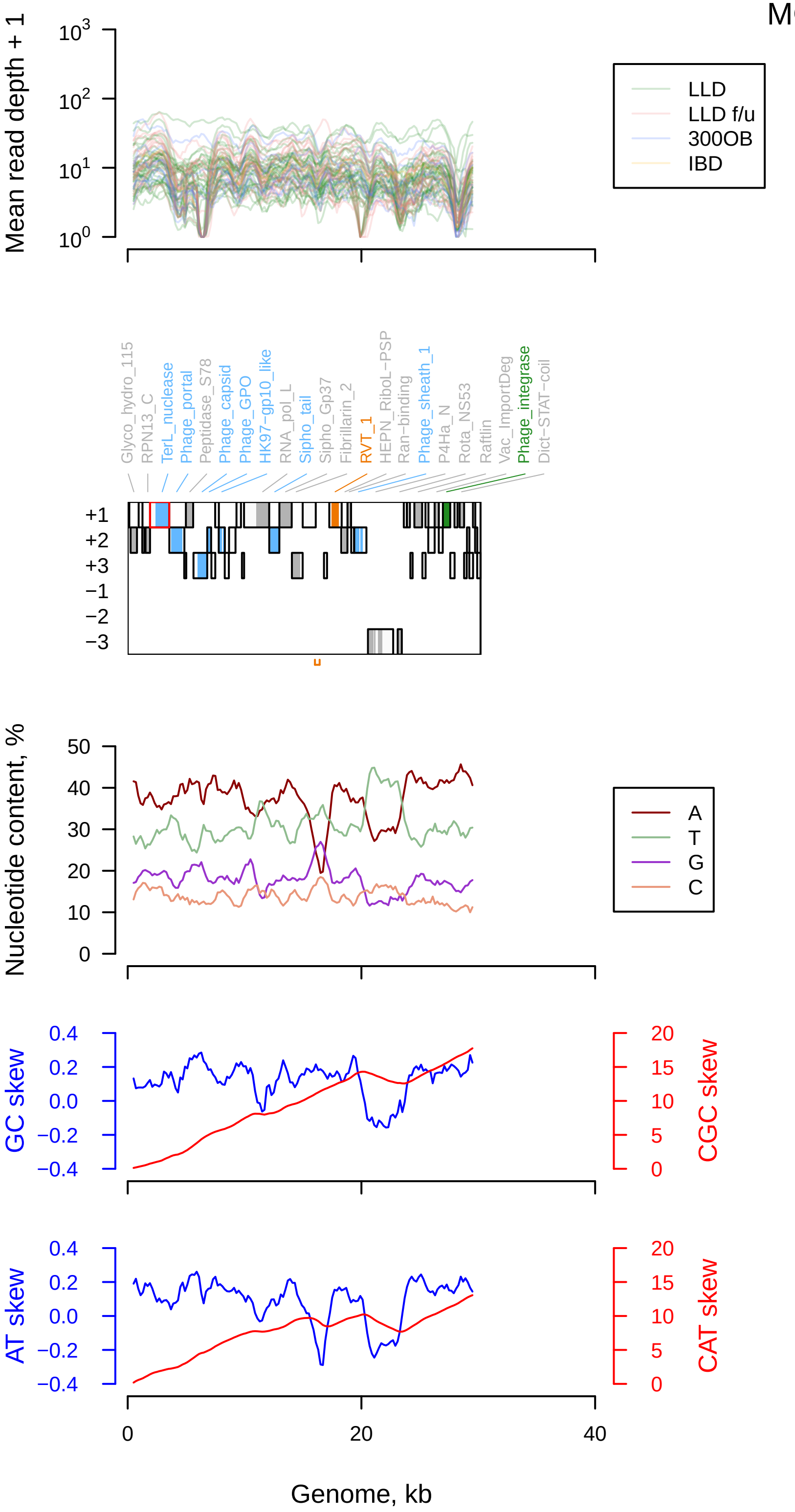

Han_2018_ERR1398212_NODE_689_length_32728_cov_10.495210 (DTR, code 11)

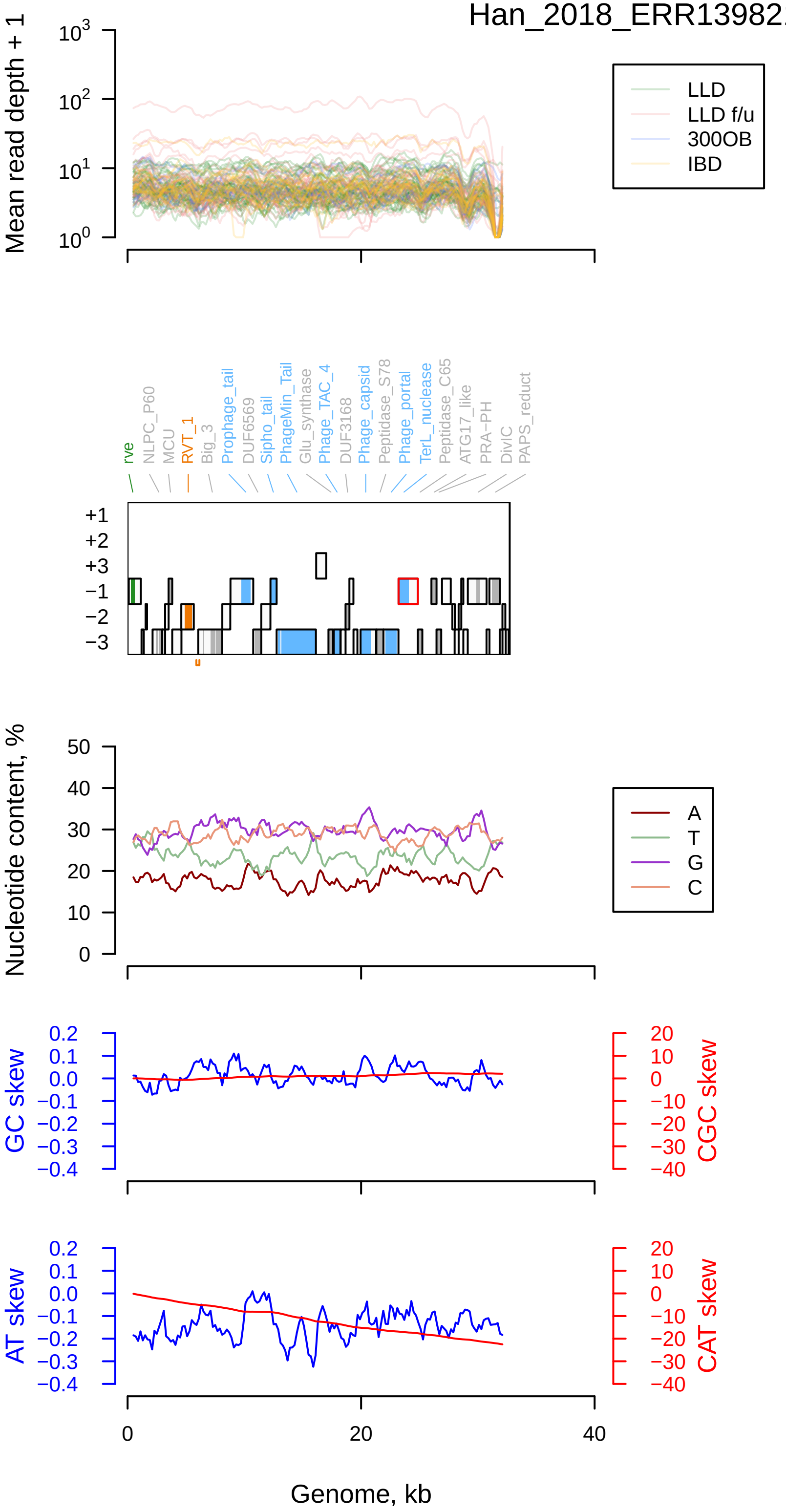

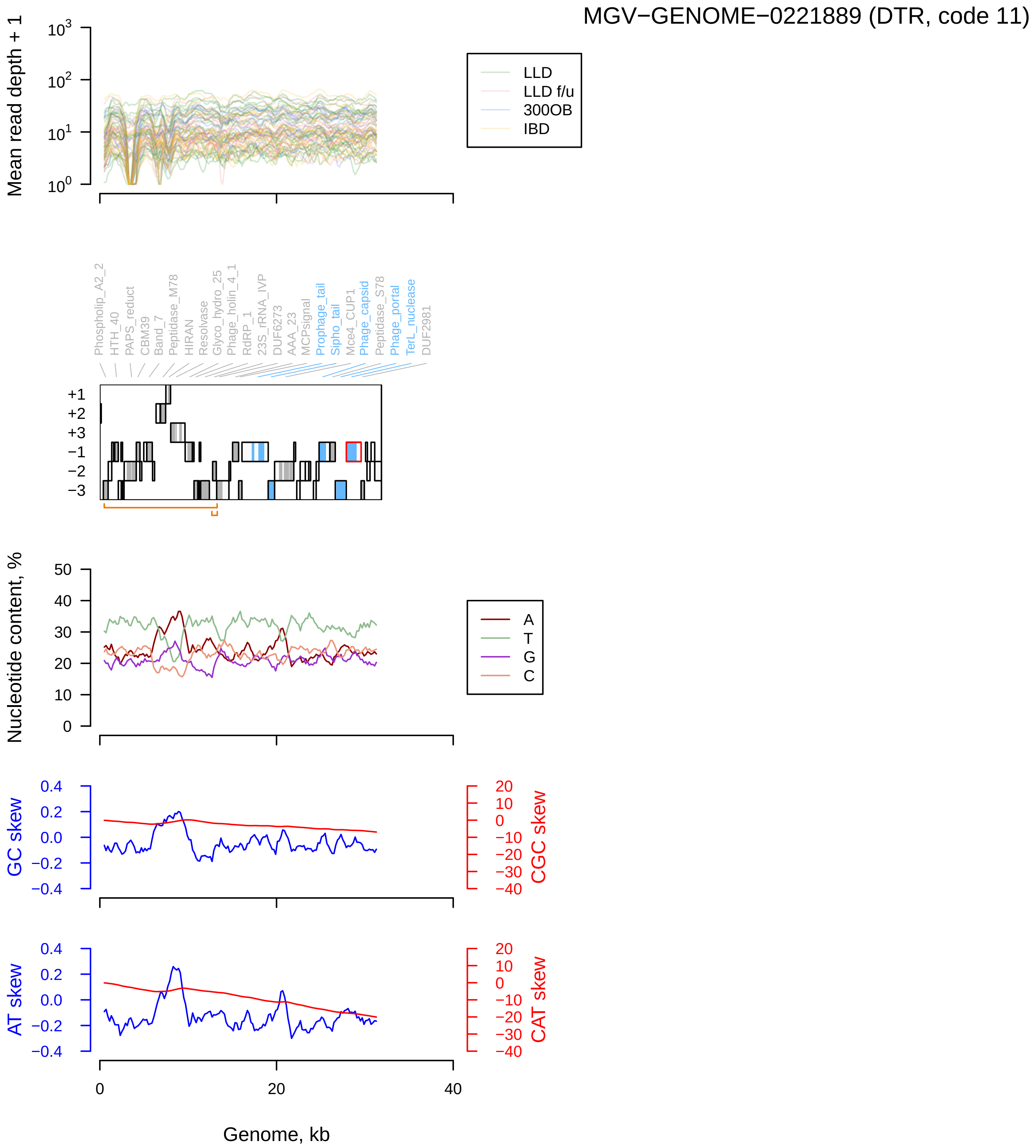

MGV−GENOME−0221889 (DTR, code 11)
Mean read depth + 1
LLD
LLD f/u
300OB
IBD
Phospholip_A2_2
HTH_40
PAPS_reduct
CBM39
Band_7
Peptidase_M78
HIRAN
Resolvase
Glyco_hydro_25
Phage_holin_4_1
RdRP_1
23S_rRNA_IVP
DUF6273
AAA_23
MCPsignal
Prophage_tail
Sipho_tail
Mce4_CUP1
Phage_capsid
Peptidase_S78
Phage_portal
TerL_nuclease
DUF2981
Nucleotide content, %
A
T
G
C
GC skew
CGC skew
AT skew
CAT skew
Genome, kb

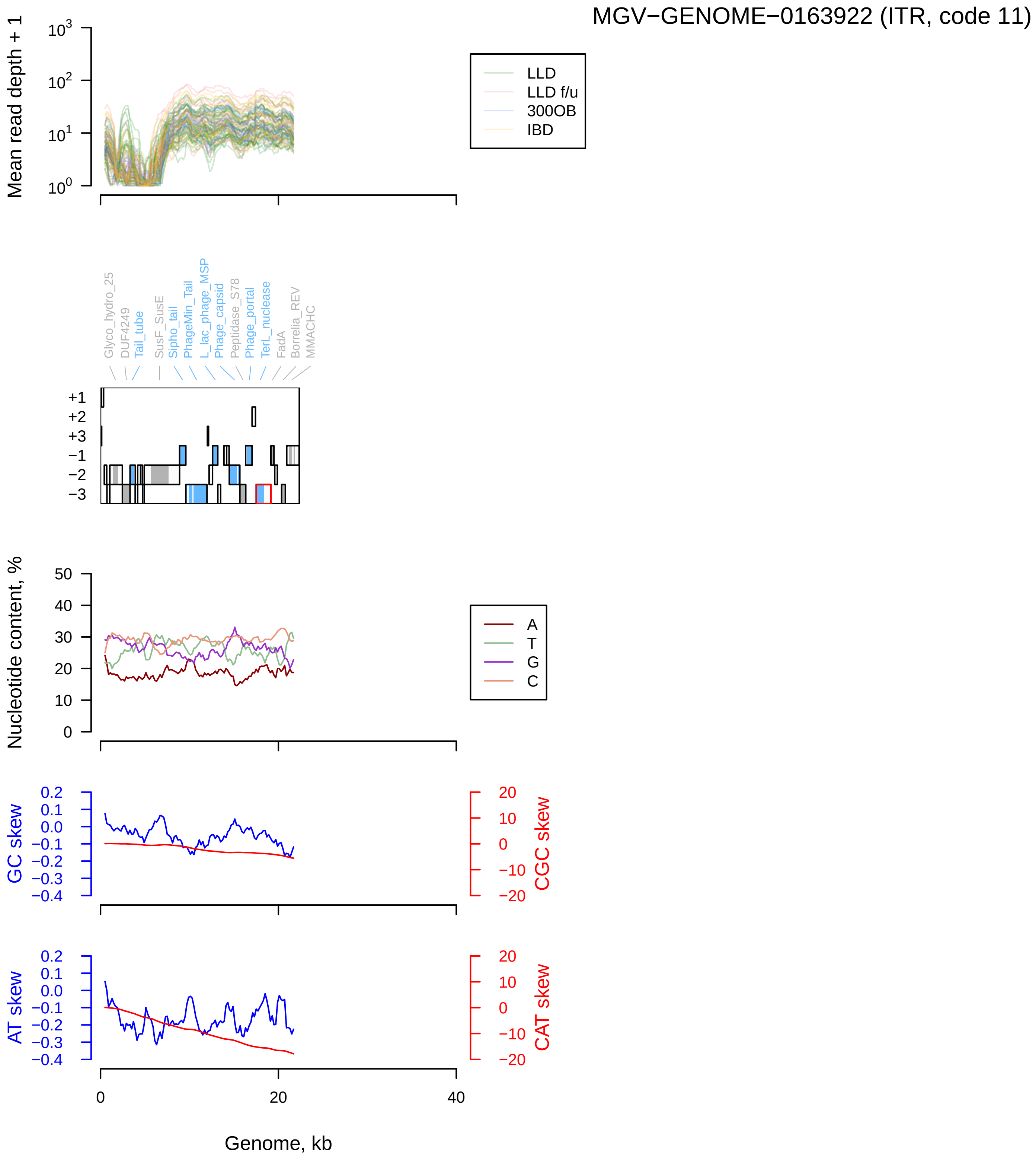
MGV−GENOME−0163922 (ITR, code 11)
Mean read depth + 1
LLD
LLD f/u
300OB
IBD
Glyco_hydro_25
DUF4249
Tail_tube
SusF_SusE
Sipho_tail
PhageMin_Tail
L_lac_phage_MSP
Phage_capsid
Peptidase_S78
Phage_portal
TerL_nuclease
FadA
Borrelia_REV
MMACHC
Nucleotide content, %
A
T
G
C
GC skew
CGC skew
AT skew
CAT skew
Genome, kb

### MGV−GENOME−0244886 (DTR, code 15)

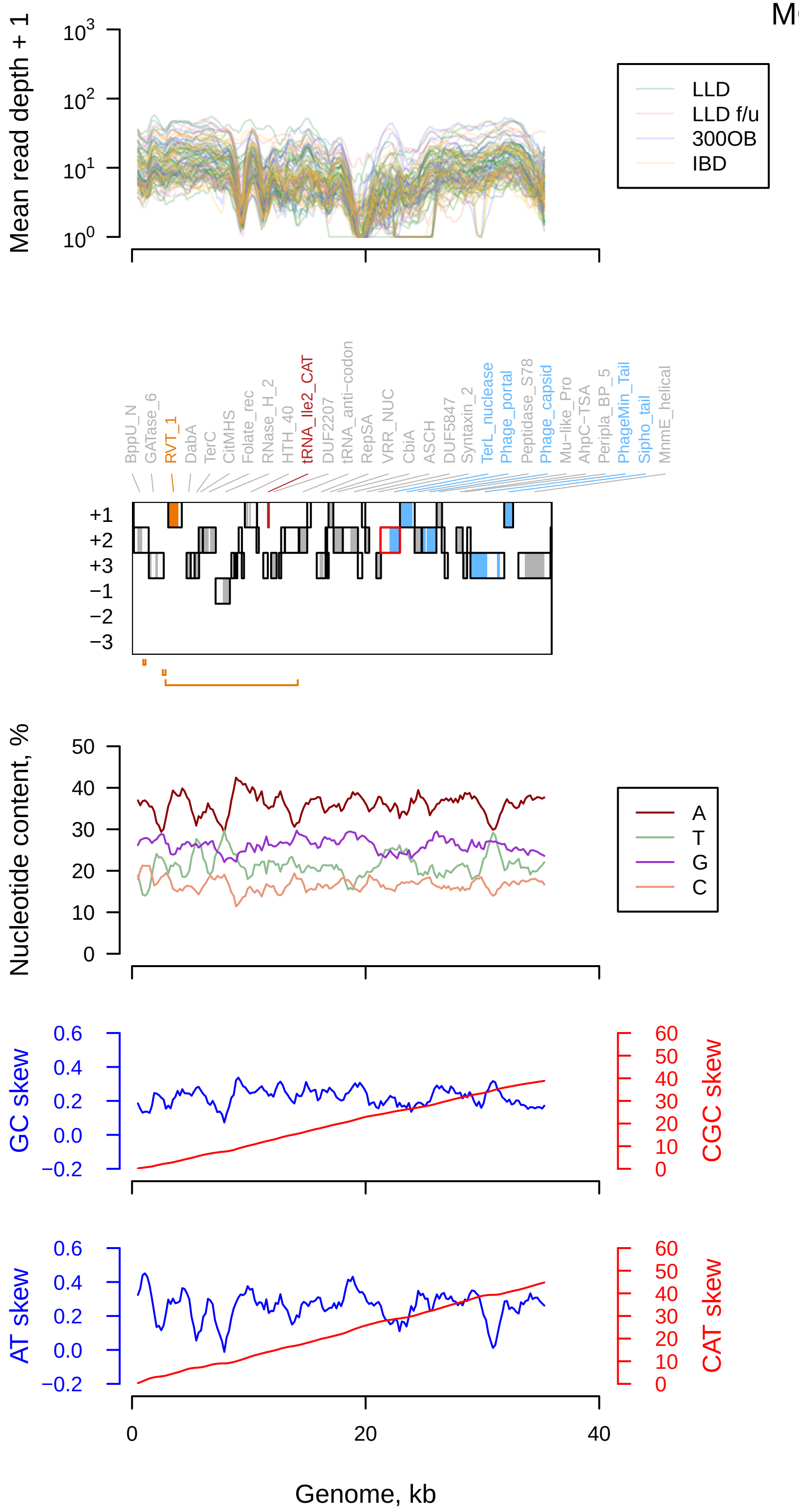

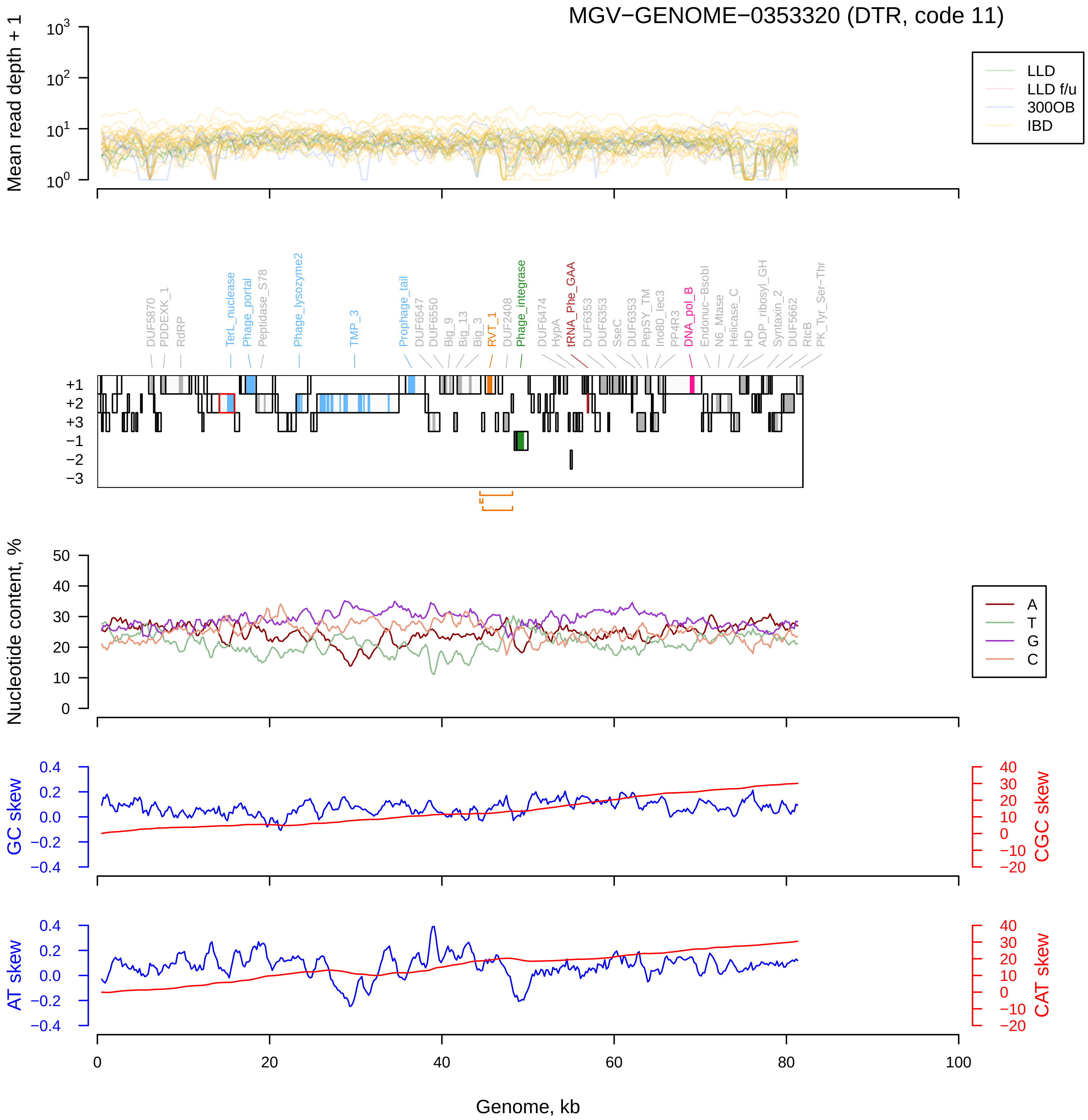

MGV−GENOME−0353320 (DTR, code 11)
Mean read depth + 1
LLD
LLD f/u
300OB
IBD
DUF5870
PDDEXK_1
RdRP
TerL_nuclease
Phage_portal
Peptidase_S78
Phage_lysozyme2
TMP_3
Prophage_tail
DUF6547
DUF6550
Big_9
Big_13
Big_3
RVT_1
DUF2408
Phage_integrase
DUF6474
HypA
tRNA_Phe_GAA
DUF6353
DUF6353
SseC
DUF6353
PepSY_TM
Ino80_Iec3
PP4R3
DNA_pol_B
Endonuc−BsobI
N6_Mtase
Helicase_C
HD
ADP_ribosyl_GH
Syntaxin_2
DUF5662
RtcB
PK_Tyr_Ser−Thr
Nucleotide content, %
A
T
G
C
GC skew
CGC skew
AT skew
CAT skew
Genome, kb

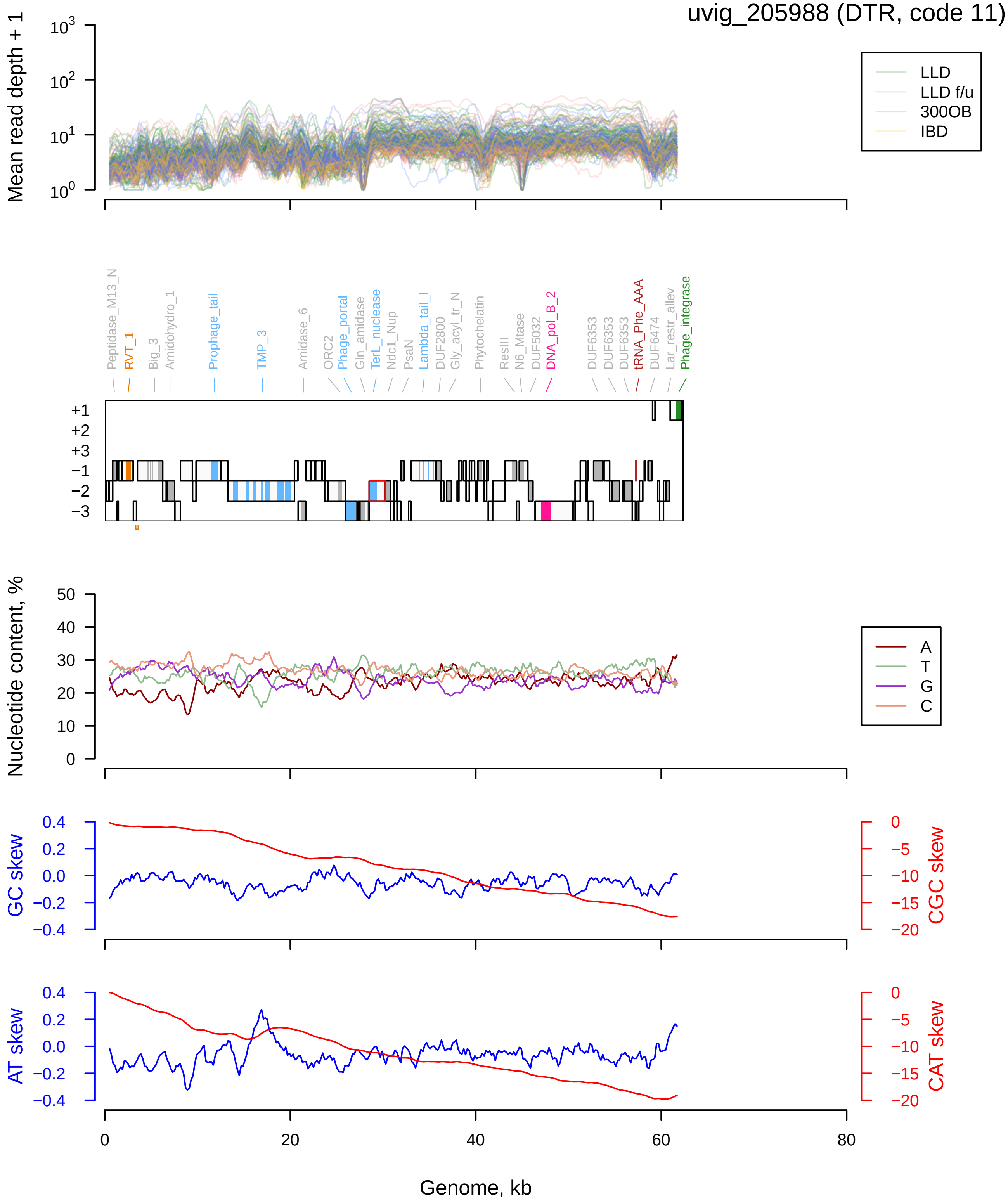

uvig_205988 (DTR, code 11)
Mean read depth + 1
LLD
LLD f/u
300OB
IBD
Peptidase_M13_N
RVT_1
Big_3
Amidohydro_1
Prophage_tail
TMP_3
Amidase_6
ORC2
Phage_portal
Gln_amidase
TerL_nuclease
Ndc1_Nup
PsaN
Lambda_tail_I
DUF2800
Gly_acyl_tr_N
Phytochelatin
ResIII
N6_Mtase
DUF5032
DNA_pol_B_2
DUF6353
DUF6353
DUF6353
tRNA_Phe_AAA
DUF6474
Lar_restr_allev
Phage_integrase
Nucleotide content, %
A
T
G
C
GC skew
CGC skew
AT skew
CAT skew
Genome, kb

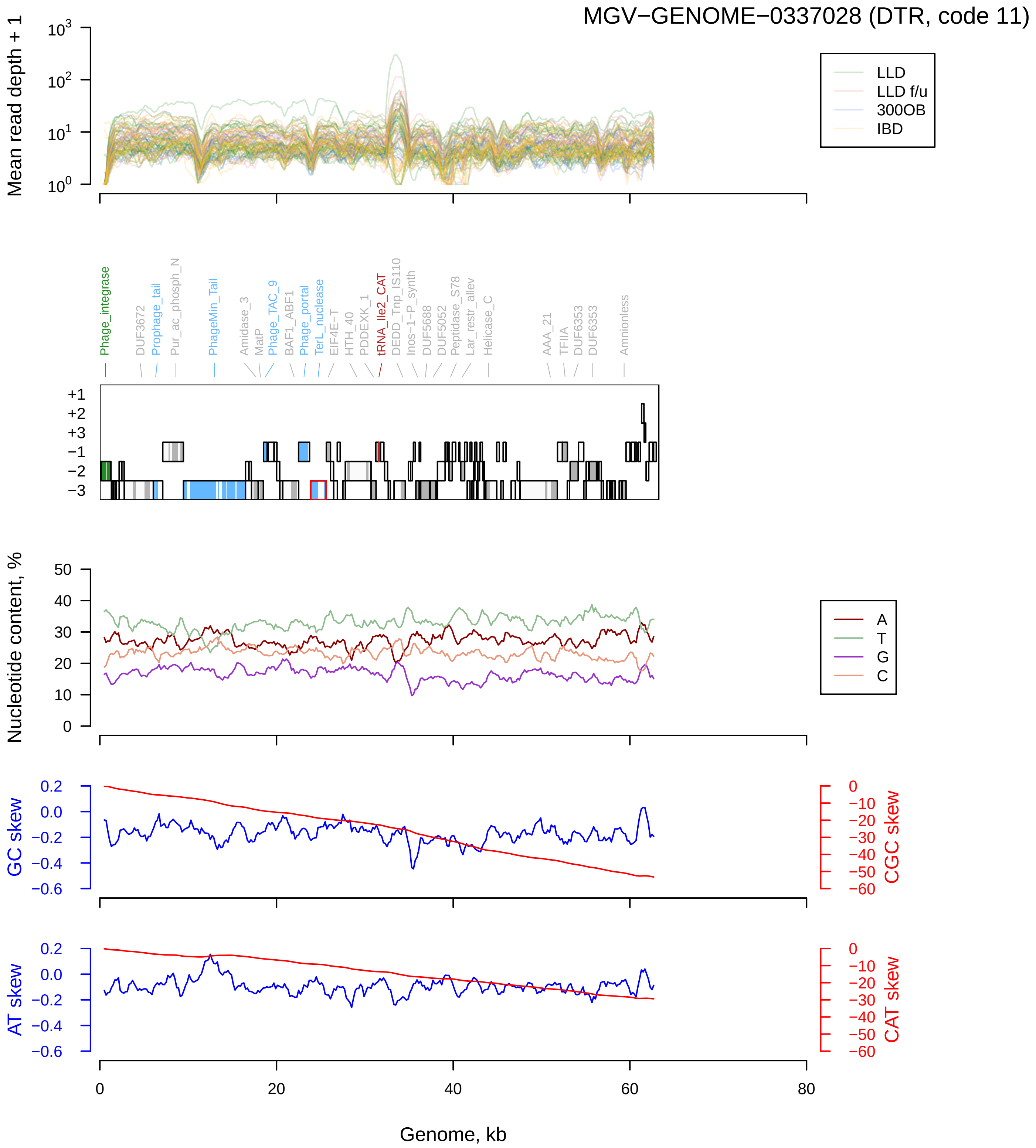

MGV−GENOME−0337028 (DTR, code 11)
Mean read depth + 1
LLD
LLD f/u
300OB
IBD
Phage_integrase
DUF3672
Prophage_tail
Pur_ac_phosph_N
PhageMin_Tail
Amidase_3
MatP
Phage_TAC_9
BAF1_ABF1
Phage_portal
TerL_nuclease
EIF4E-T
HTH_40
PDDEXK_1
tRNA_Ile2_CAT
DEDD_Tnp_IS110
Inos-1-P_synth
DUF5688
DUF5052
Peptidase_S78
Lar_restr_allev
Helicase_C
AAA_21
TFIIA
DUF6353
DUF6353
Amnionless
Nucleotide content, %
A
T
G
C
GC skew
CGC skew
AT skew
CAT skew
Genome, kb

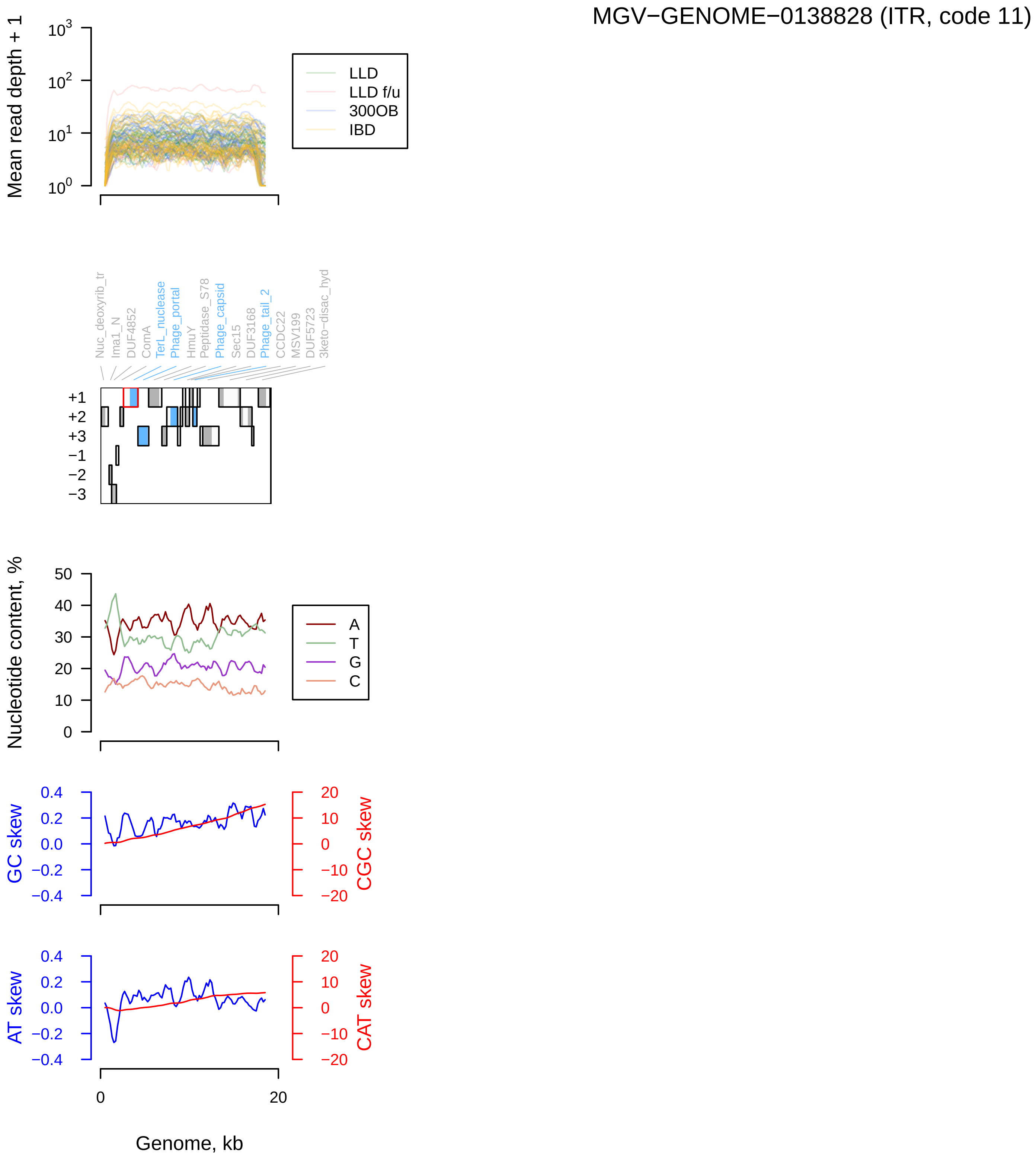
MGV−GENOME−0138828 (ITR, code 11)
Mean read depth + 1
LLD
LLD f/u
300OB
IBD
Nuc_deoxyrib_tr
Ima1_N
DUF4852
ComA
TerL_nuclease
Phage_portal
HmuY
Peptidase_S78
Phage_capsid
Sec15
DUF3168
Phage_tail_2
CCDC22
MSV199
DUF5723
3keto−disac_hyd
Nucleotide content, %
A
T
G
C
GC skew
CGC skew
AT skew
CAT skew
Genome, kb

MGV−GENOME−0109615 (ITR, code 11)

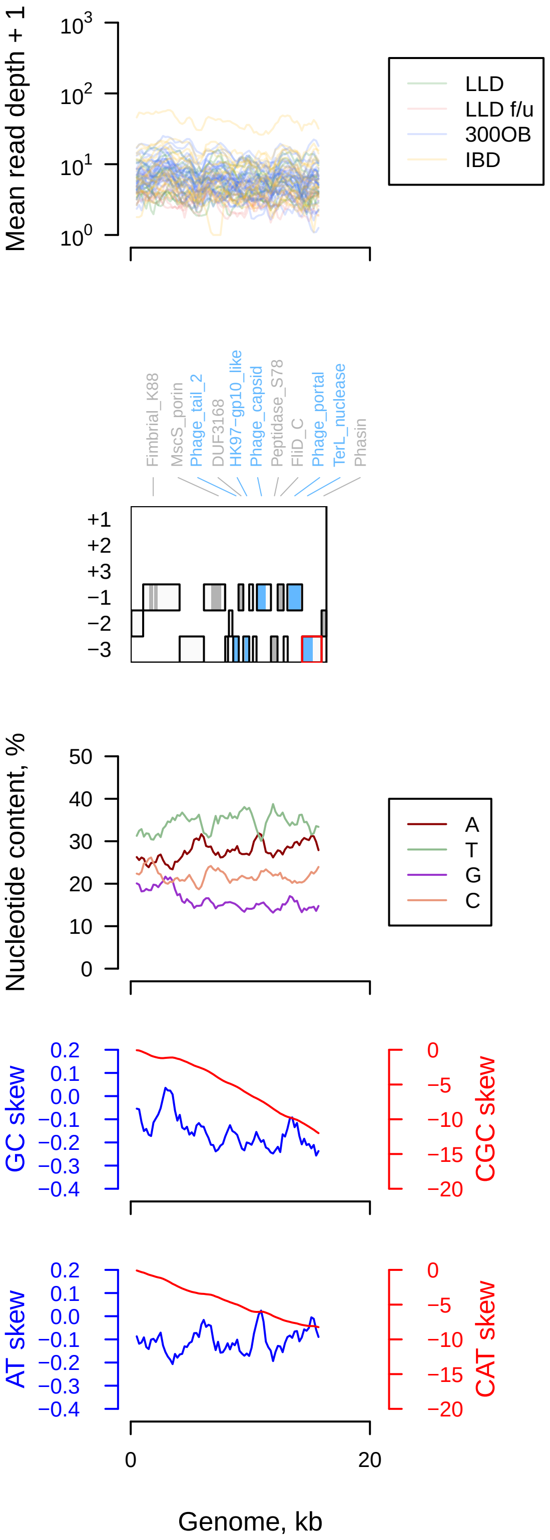

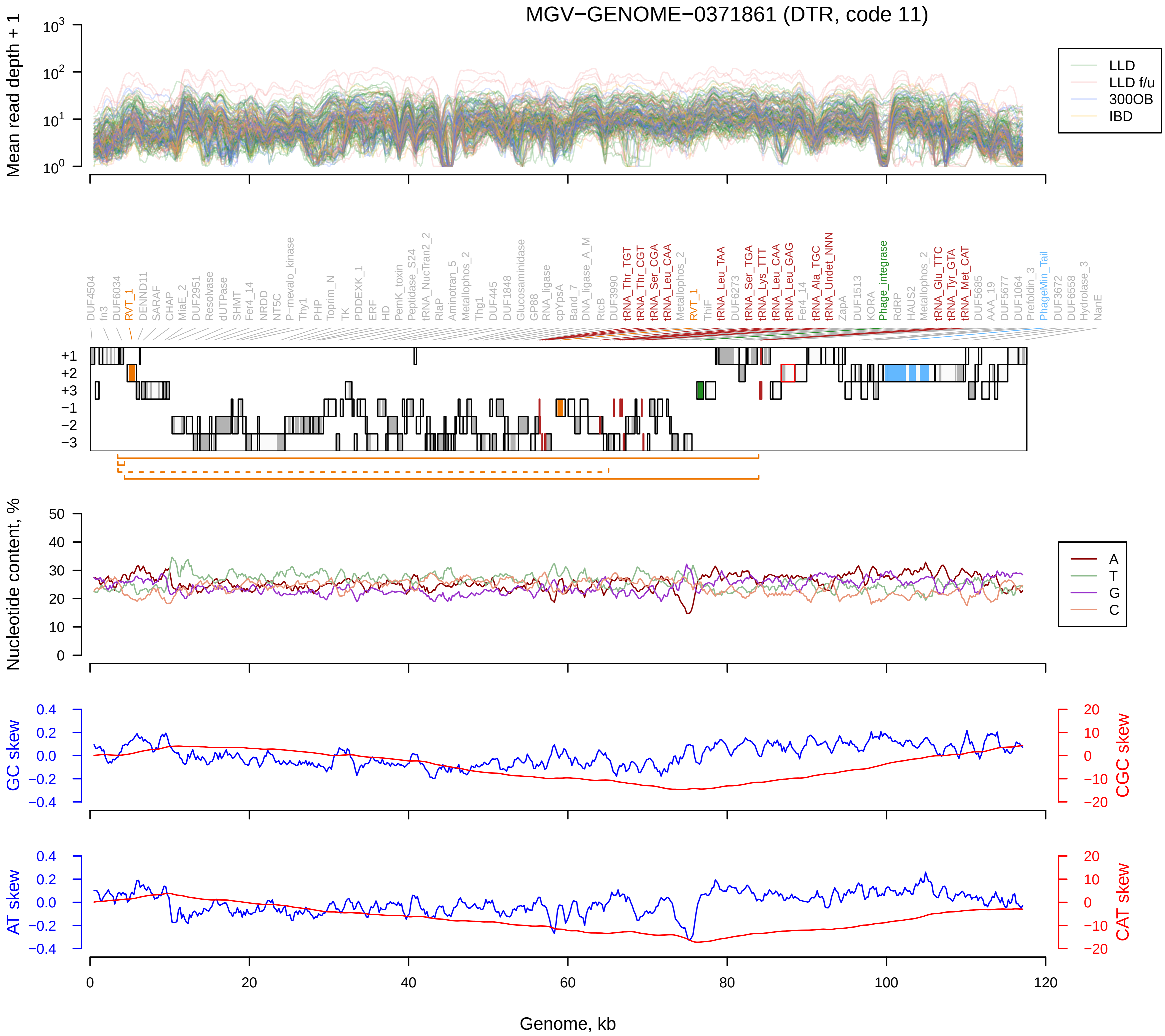
MGV−GENOME−0371861 (DTR, code 11)
Mean read depth + 1
LLD
LLD f/u
300OB
IBD
DUF4504
fn3
DUF6034
RVT_1
DENND11
SARAF
CHAP
MlaE_2
DUF2951
Resolvase
dUTPase
SHMT
Fer4_14
NRDD
NT5C
P−mevalo_kinase
Thy1
PHP
Toprim_N
TK
PDDEXK_1
ERF
HD
PemK_toxin
Peptidase_S24
tRNA_NucTran2_2
RlaP
Aminotran_5
Metallophos_2
Thg1
DUF445
DUF1848
Glucosaminidase
GP88
RNA_ligase
cpYpsA
Band_7
DNA_ligase_A_M
RtcB
DUF3990
tRNA_Thr_TGT
tRNA_Thr_CGT
tRNA_Ser_CGA
tRNA_Leu_CAA
Metallophos_2
RVT_1
ThiF
tRNA_Leu_TAA
DUF6273
tRNA_Ser_TGA
tRNA_Lys_TTT
tRNA_Leu_CAA
tRNA_Leu_GAG
Fer4_14
tRNA_Ala_TGC
tRNA_Undet_NNN
ZapA
DUF1513
KORA
Phage_integrase
RdRP
HAUS2
Metallophos_2
tRNA_Glu_TTC
tRNA_Tyr_GTA
tRNA_Met_CAT
DUF5685
AAA_19
DUF5677
DUF1064
Prefoldin_3
PhageMin_Tail
DUF3672
DUF6558
Hydrolase_3
NanE
+1
+2
+3
−1
−2
−3
Nucleotide content, %
A
T
G
C
GC skew
CGC skew
AT skew
CAT skew
Genome, kb

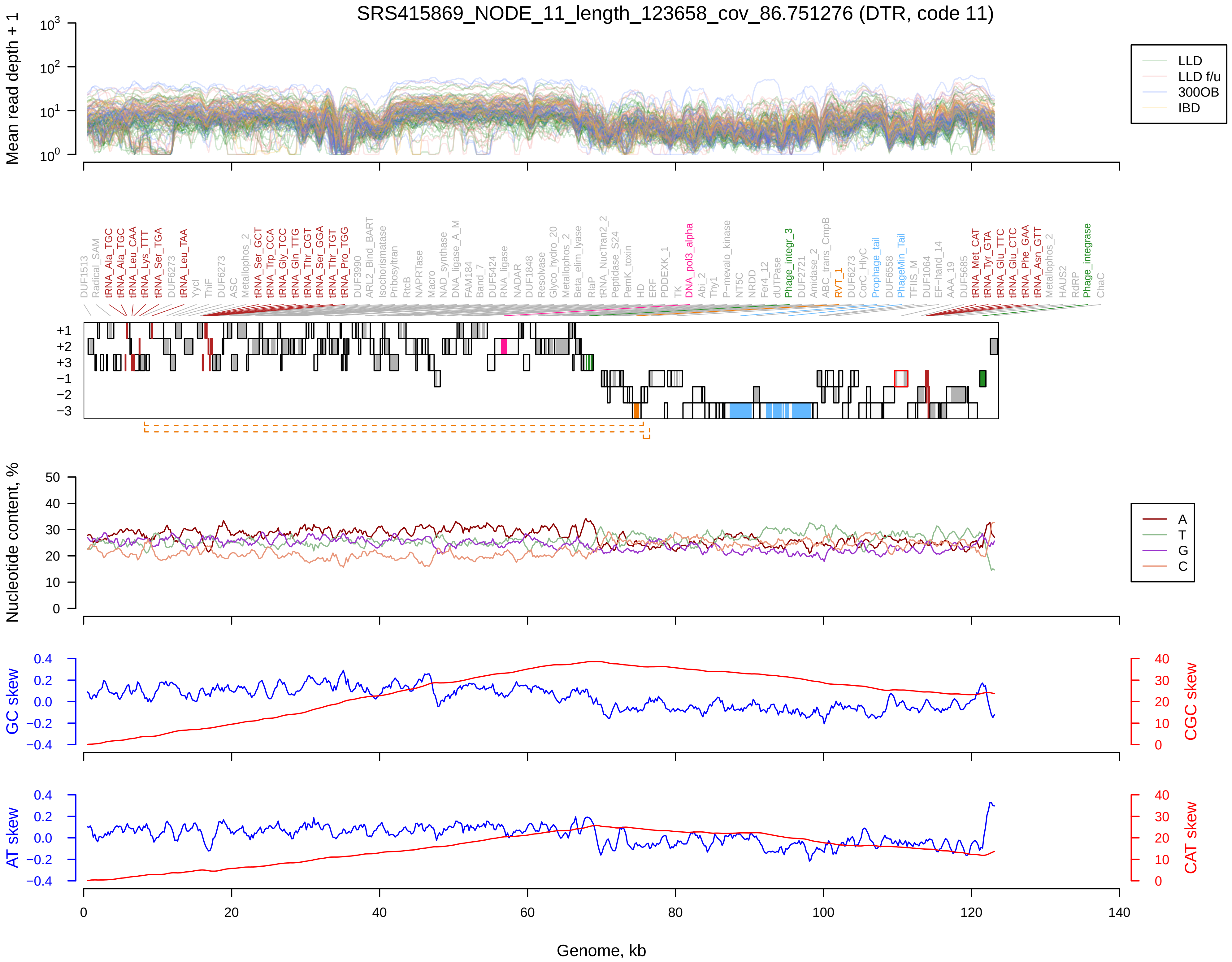
SRS415869_NODE_11_length_123658_cov_86.751276 (DTR, code 11)
Mean read depth + 1
LLD
LLD f/u
300OB
IBD
DUF1513
Radical_SAM
tRNA_Ala_TGC
tRNA_Ala_TGC
tRNA_Leu_CAA
tRNA_Lys_TTT
tRNA_Ser_TGA
DUF6273
tRNA_Leu_TAA
YcjI
ThiF
DUF6273
ASC
Metallophos_2
tRNA_Ser_GCT
tRNA_Trp_CCA
tRNA_Gly_TCC
tRNA_Gln_TTG
tRNA_Thr_CGT
tRNA_Ser_GGA
tRNA_Thr_TGT
tRNA_Pro_TGG
DUF3990
ARL2_Bind_BART
Isochorismatase
Pribosyltran
RtcB
NAPRTase
Macro
NAD_synthase
DNA_ligase_A_M
FAM184
Band_7
DUF5424
RNA_ligase
NADAR
DUF1848
Resolvase
Glyco_hydro_20
Metallophos_2
Beta_elim_lyase
RlaP
tRNA_NucTran2_2
Peptidase_S24
PemK_toxin
HD
ERF
PDDEXK_1
TK
DNA_pol3_alpha
Abi_2
Thy1
P-mevalo_kinase
NT5C
NRDD
Fer4_12
dUTPase
Phage_integr_3
DUF2721
Amidase_2
ABC_trans_CmpB
RVT_1
DUF6273
CorC_HlyC
Prophage_tail
DUF6558
PhageMin_Tail
TFIIS_M
DUF1064
EF-hand_14
AAA_19
DUF5685
tRNA_Met_CAT
tRNA_Tyr_GTA
tRNA_Glu_TTC
tRNA_Glu_CTC
tRNA_Phe_GAA
tRNA_Asn_GTT
Metallophos_2
HAUS2
RdRP
Phage_integrase
ChaC
Nucleotide content, %
A
T
G
C
GC skew
CGC skew
AT skew
CAT skew
Genome, kb

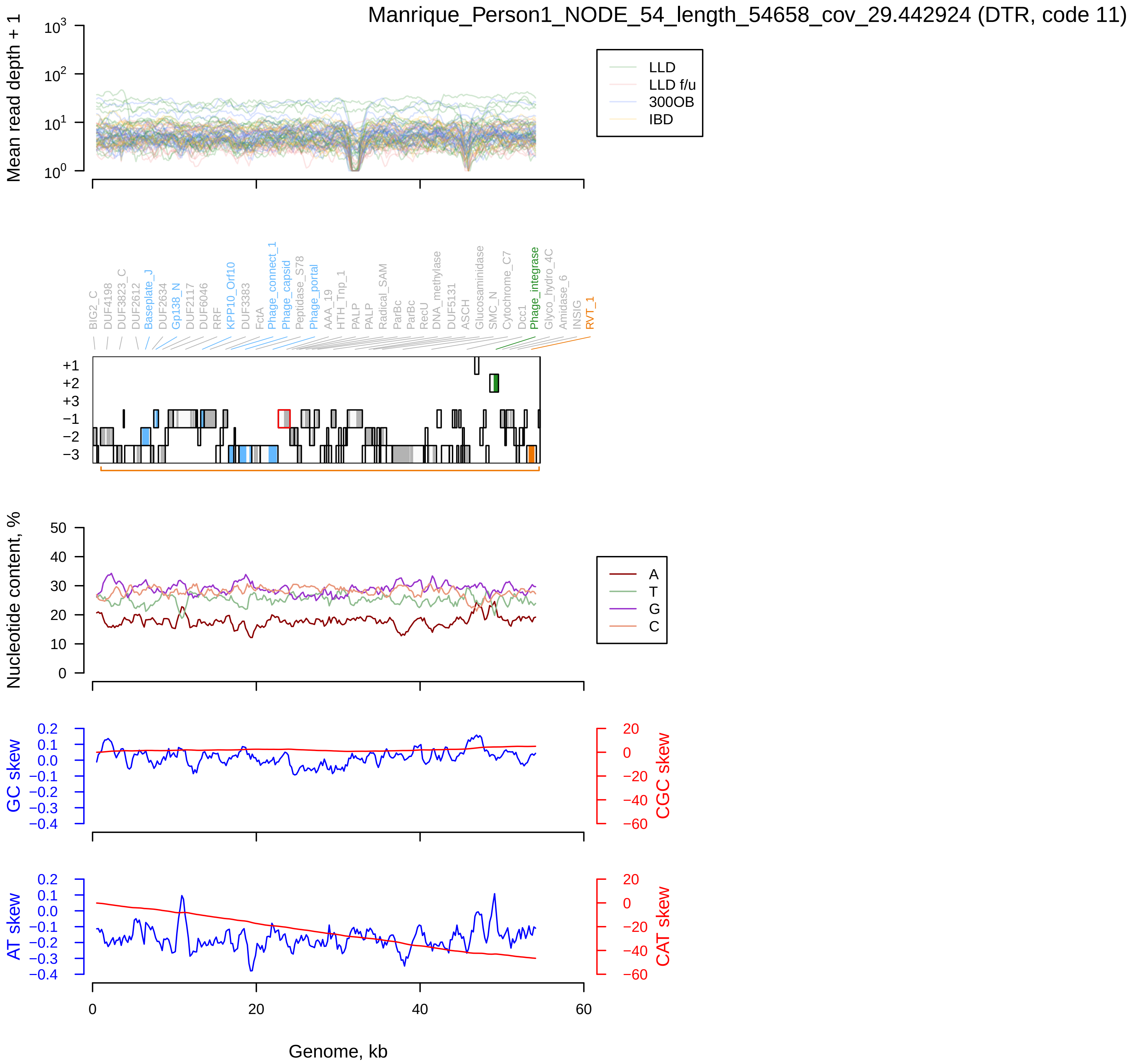

Manrique_Person1_NODE_54_length_54658_cov_29.442924 (DTR, code 11)
Mean read depth + 1
LLD
LLD f/u
300OB
IBD
BIG2_C
DUF4198
DUF3823_C
DUF2612
Baseplate_J
DUF2634
Gp138_N
DUF2117
DUF6046
RRF
KPP10_Orf10
DUF3383
FctA
Phage_connect_1
Phage_capsid
Peptidase_S78
Phage_portal
AAA_19
HTH_Tnp_1
PALP
PALP
Radical_SAM
ParBc
ParBc
RecU
DNA_methylase
DUF5131
ASCH
Glucosaminidase
SMC_N
Cytochrome_C7
Dcc1
Phage_integrase
Glyco_hydro_4C
Amidase_6
INSIG
RVT_1
Nucleotide content, %
A
T
G
C
GC skew
CGC skew
AT skew
CAT skew
Genome, kb

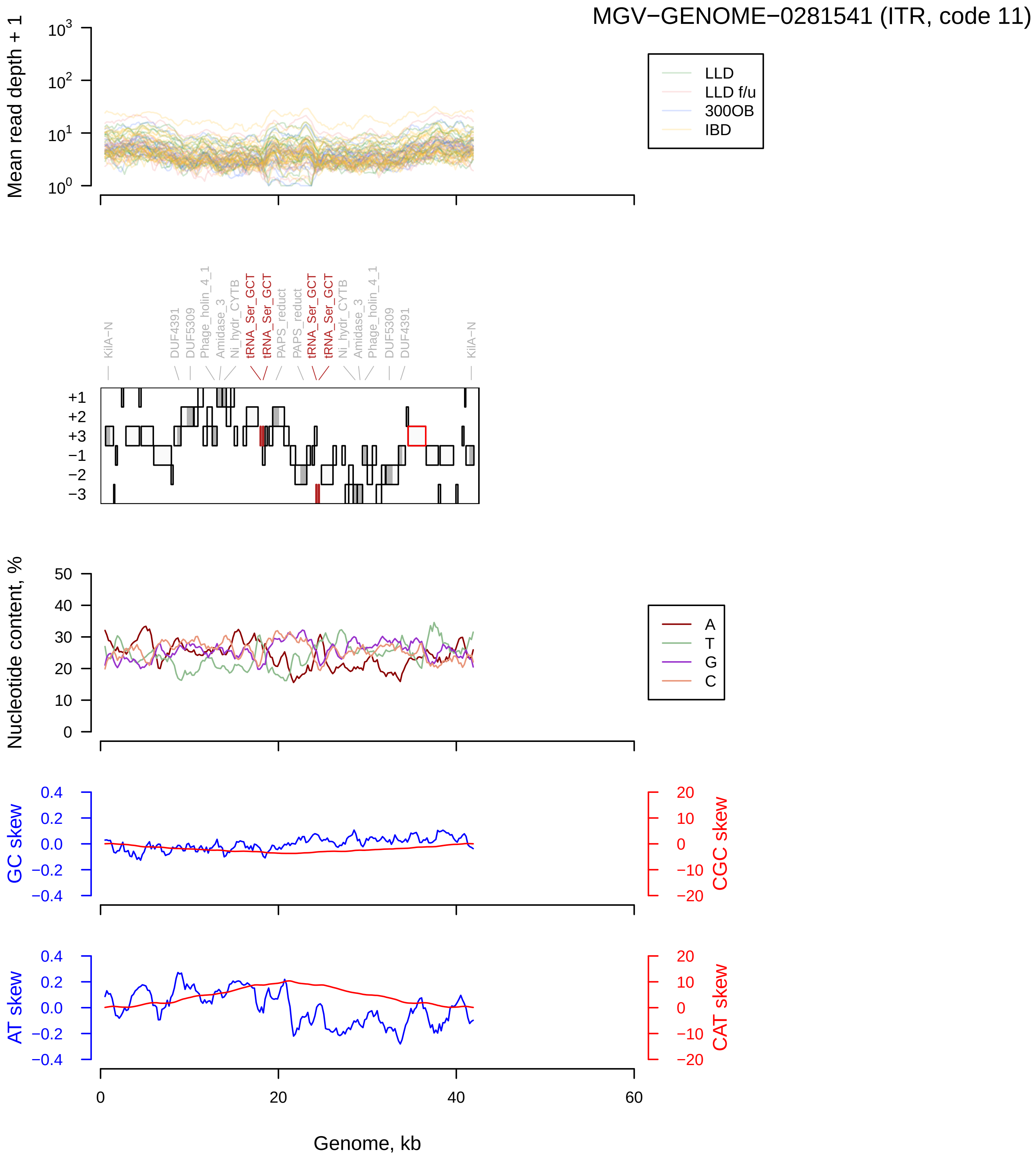

MGV−GENOME−0281541 (ITR, code 11)
Mean read depth + 1
LLD
LLD f/u
300OB
IBD
KilA−N
DUF4391
DUF5309
Phage_holin_4_1
Amidase_3
Ni_hydr_CYTB
tRNA_Ser_GCT
tRNA_Ser_GCT
PAPS_reduct
PAPS_reduct
tRNA_Ser_GCT
tRNA_Ser_GCT
Ni_hydr_CYTB
Amidase_3
Phage_holin_4_1
DUF5309
DUF4391
KilA−N
Nucleotide content, %
A
T
G
C
GC skew
CGC skew
AT skew
CAT skew
Genome, kb

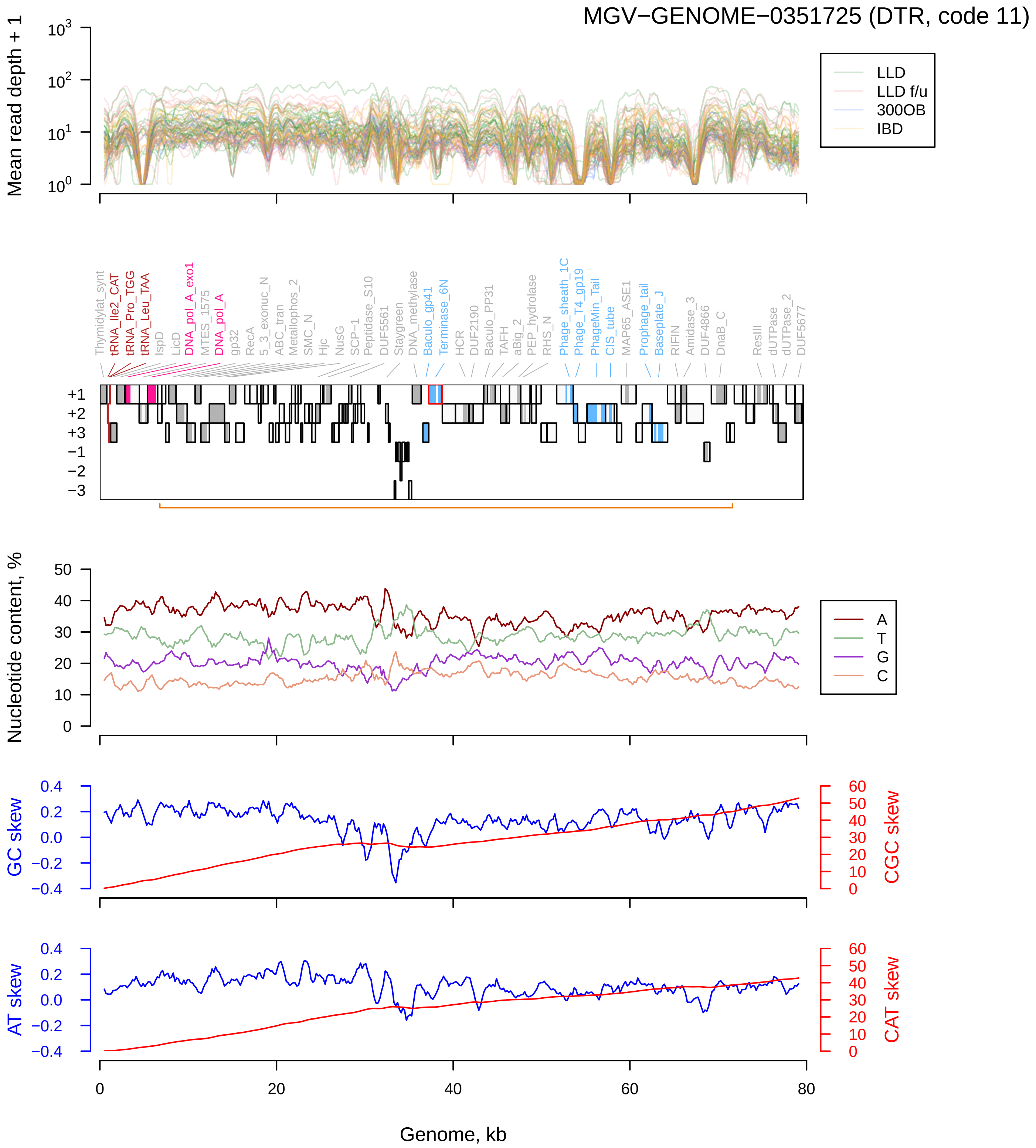
MGV−GENOME−0351725 (DTR, code 11)
Mean read depth + 1
LLD
LLD f/u
300OB
IBD
Thymidylat_synt
tRNA_Ile2_CAT
tRNA_Pro_TGG
tRNA_Leu_TAA
IspD
LicD
DNA_pol_A_exo1
MTES_1575
DNA_pol_A
gp32
RecA
5_3_exonuc_N
ABC_tran
Metallophos_2
SMC_N
Hjc
NusG
SCP-1
Peptidase_S10
DUF5561
Staygreen
DNA_methylase
Baculo_gp41
Terminase_6N
HCR
DUF2190
Baculo_PP31
TAFH
aBig_2
PEP_hydrolase
RHS_N
Phage_sheath_1C
Phage_T4_gp19
PhageMin_Tail
CIS_tube
MAP65_ASE1
Prophage_tail
Baseplate_J
RIFIN
Amidase_3
DUF4866
DnaB_C
ResIII
dUTPase
dUTPase_2
DUF5677
+1
+2
+3
−1
−2
−3
Nucleotide content, %
A
T
G
C
GC skew
CGC skew
AT skew
CAT skew
Genome, kb

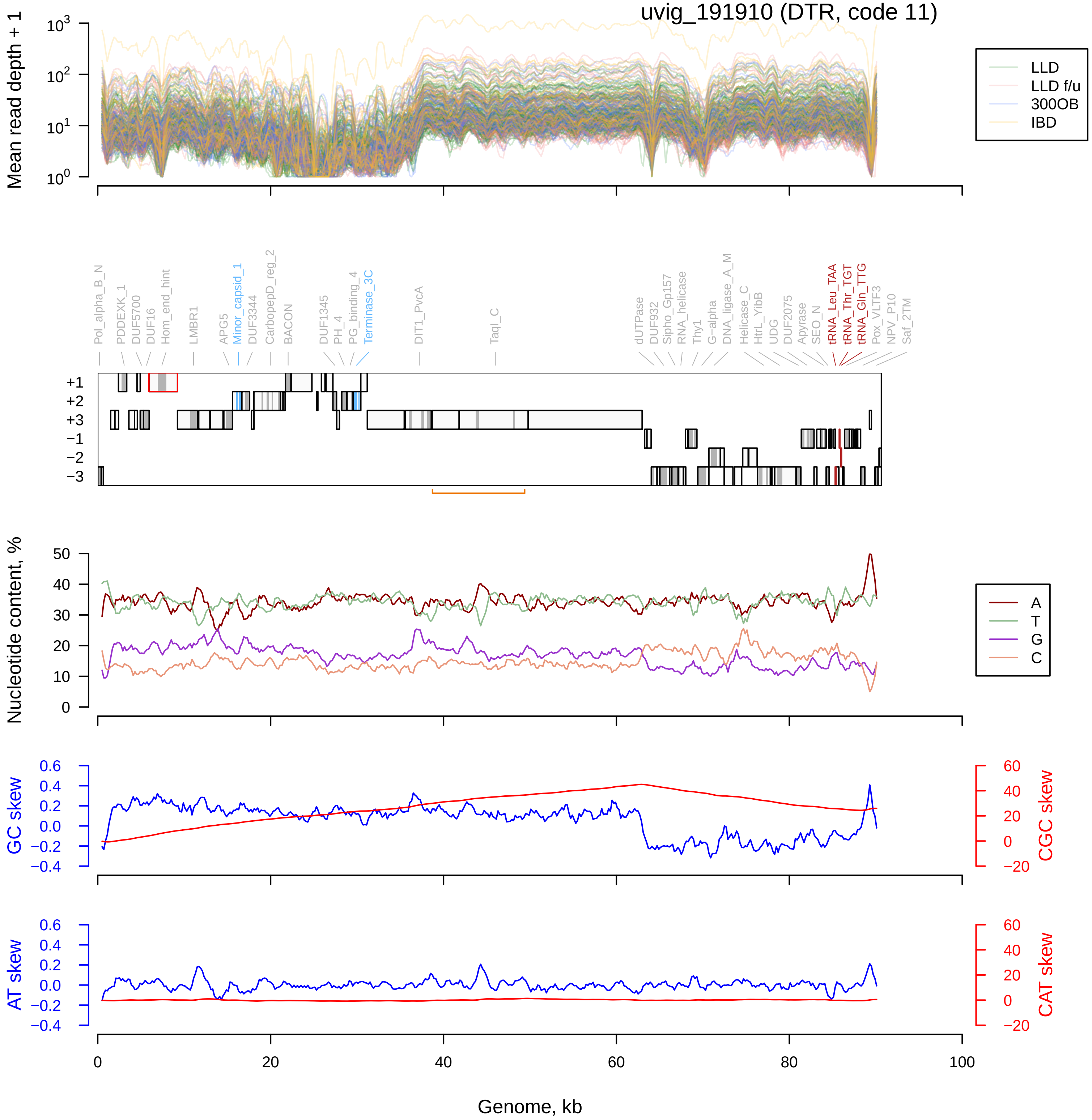
uvig_191910 (DTR, code 11)
Mean read depth + 1
LLD
LLD f/u
300OB
IBD
Pol_alpha_B_N
PDDEXK_1
DUF5700
DUF16
Hom_end_hint
LMBR1
APG5
Minor_capsid_1
DUF3344
CarbopepD_reg_2
BACON
DUF1345
PH_4
PG_binding_4
Terminase_3C
DIT1_PvcA
TaqI_C
dUTPase
DUF932
Sipho_Gp157
RNA_helicase
Thy1
G-alpha
DNA_ligase_A_M
Helicase_C
HtrL_YibB
UDG
DUF2075
Apyrase
SEO_N
tRNA_Leu_TAA
tRNA_Thr_TGT
tRNA_Gln_TTG
Pox_VLTF3
NPV_P10
Saf_2TM
Nucleotide content, %
A
T
G
C
GC skew
CGC skew
AT skew
CAT skew
Genome, kb

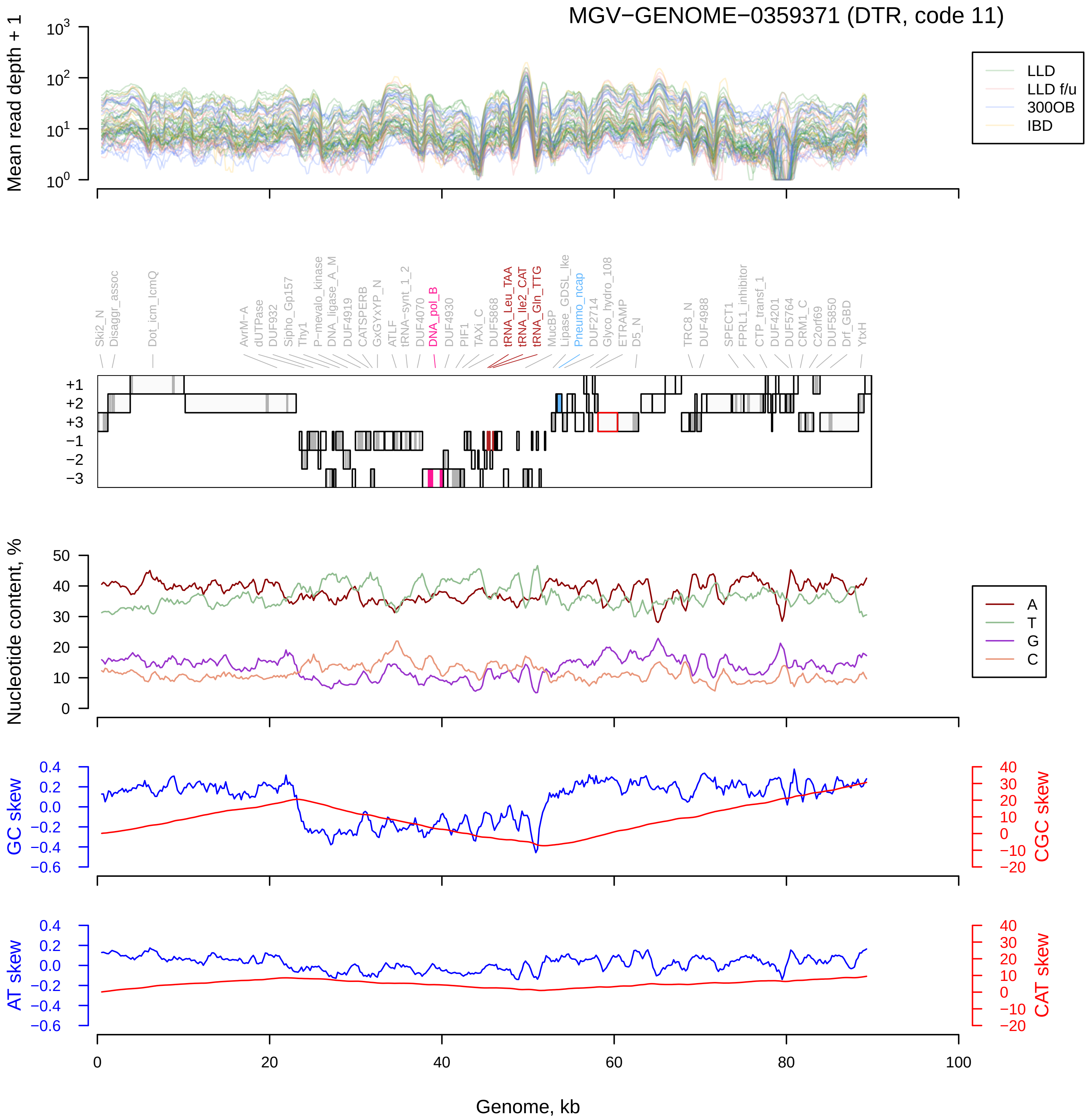
MGV−GENOME−0359371 (DTR, code 11)
Mean read depth + 1
LLD
LLD f/u
300OB
IBD
Ski2_N
Disaggr_assoc
Dot_icm_IcmQ
AvrM−A
dUTPase
DUF932
Sipho_Gp157
Thy1
P−mevalo_kinase
DNA_ligase_A_M
DUF4919
CATSPERB
GxGYxYP_N
ATLF
tRNA−synt_1_2
DUF4070
DNA_pol_B
DUF4930
PIF1
TAXi_C
DUF5868
tRNA_Leu_TAA
tRNA_Ile2_CAT
tRNA_Gln_TTG
MucBP
Lipase_GDSL_lke
Pneumo_ncap
DUF2714
Glyco_hydro_108
ETRAMP
D5_N
TRC8_N
DUF4988
SPECT1
FPRL1_inhibitor
CTP_transf_1
DUF4201
DUF5764
CRM1_C
C2orf69
DUF5850
Drf_GBD
YtxH
+1
+2
+3
−1
−2
−3
Nucleotide content, %
A
T
G
C
GC skew
CGC skew
AT skew
CAT skew
Genome, kb

uvig_540612 (ITR, code 15)

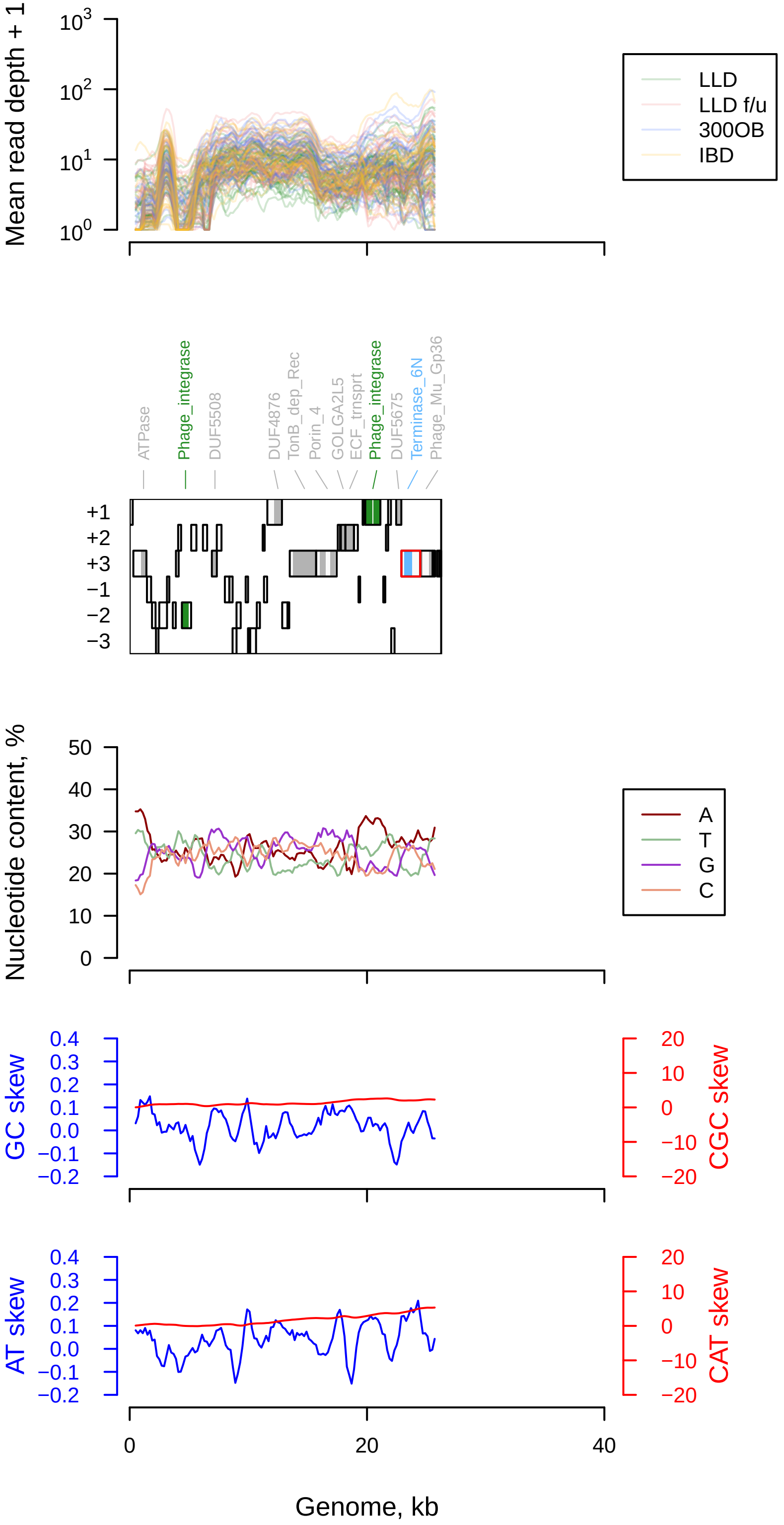

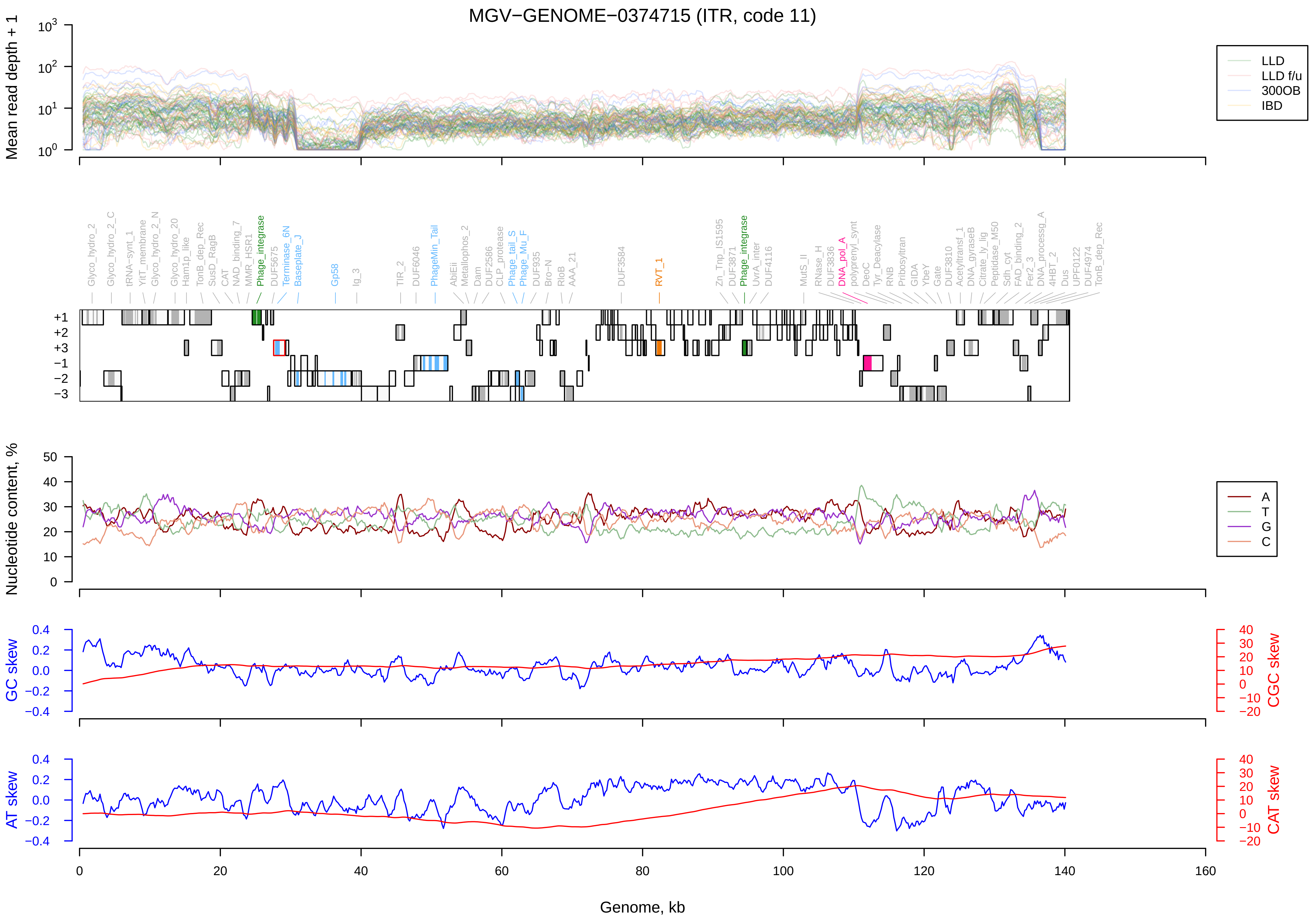
MGV−GENOME−0374715 (ITR, code 11)
Mean read depth + 1
LLD
LLD f/u
300OB
IBD
Glyco_hydro_2
Glyco_hydro_2_C
tRNA-synt_1
YitT_membrane
Glyco_hydro_2_N
Glyco_hydro_20
Ham1p_like
TonB_dep_Rec
SusD_RagB
CAT
NAD_binding_7
MMR_HSR1
Phage_integrase
DUF5675
Terminase_6N
Baseplate_J
Gp58
Ig_3
TIR_2
DUF6046
PhageMin_Tail
AbiEii
Metallophos_2
Dam
DUF2586
CLP_protease
Phage_tail_S
Phage_Mu_F
DUF935
Bro−N
RloB
AAA_21
DUF3584
RVT_1
Zn_Tnp_IS1595
DUF3871
Phage_integrase
UvrA_inter
DUF4116
MutS_II
RNase_H
DUF3836
DNA_pol_A
polyprenyl_synt
DeoC
Tyr_Deacylase
RNB
Pribosyltran
GIDA
YbeY
Gate
DUF3810
Acetyltransf_1
DNA_gyraseB
Citrate_ly_lig
Peptidase_M50
Sdh_cyt
FAD_binding_2
Fer2_3
DNA_processg_A
4HBT_2
Dus
UPF0122
DUF4974
TonB_dep_Rec
+1
+2
+3
−1
−2
−3
Nucleotide content, %
A
T
G
C
GC skew
CGC skew
AT skew
CAT skew
Genome, kb
0
20
40
60
80
100
120
140
160

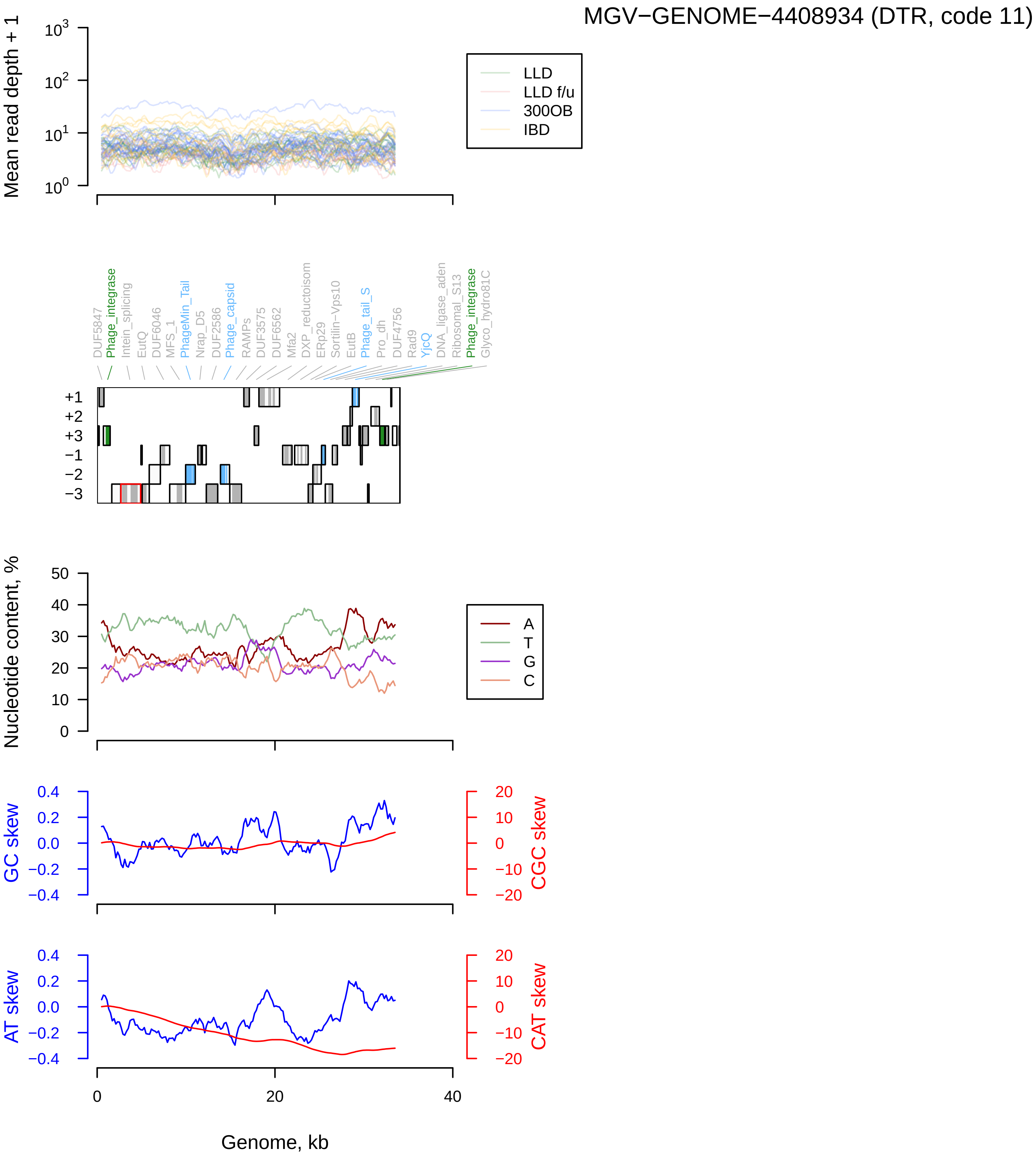
MGV−GENOME−4408934 (DTR, code 11)
Mean read depth + 1
LLD
LLD f/u
300OB
IBD
DUF5847
Phage_integrase
Intein_splicing
EutQ
DUF6046
MFS_1
PhageMin_Tail
Nrap_D5
DUF2586
Phage_capsid
RAMPs
DUF3575
DUF6562
Mfa2
DXP_reductoisom
ERp29
Sortilin−Vps10
EutB
Phage_tail_S
Pro_dh
DUF4756
Rad9
YjcQ
DNA_ligase_aden
Ribosomal_S13
Phage_integrase
Glyco_hydro81C
+1
+2
+3
−1
−2
−3
Nucleotide content, %
A
T
G
C
GC skew
CGC skew
AT skew
CAT skew
Genome, kb

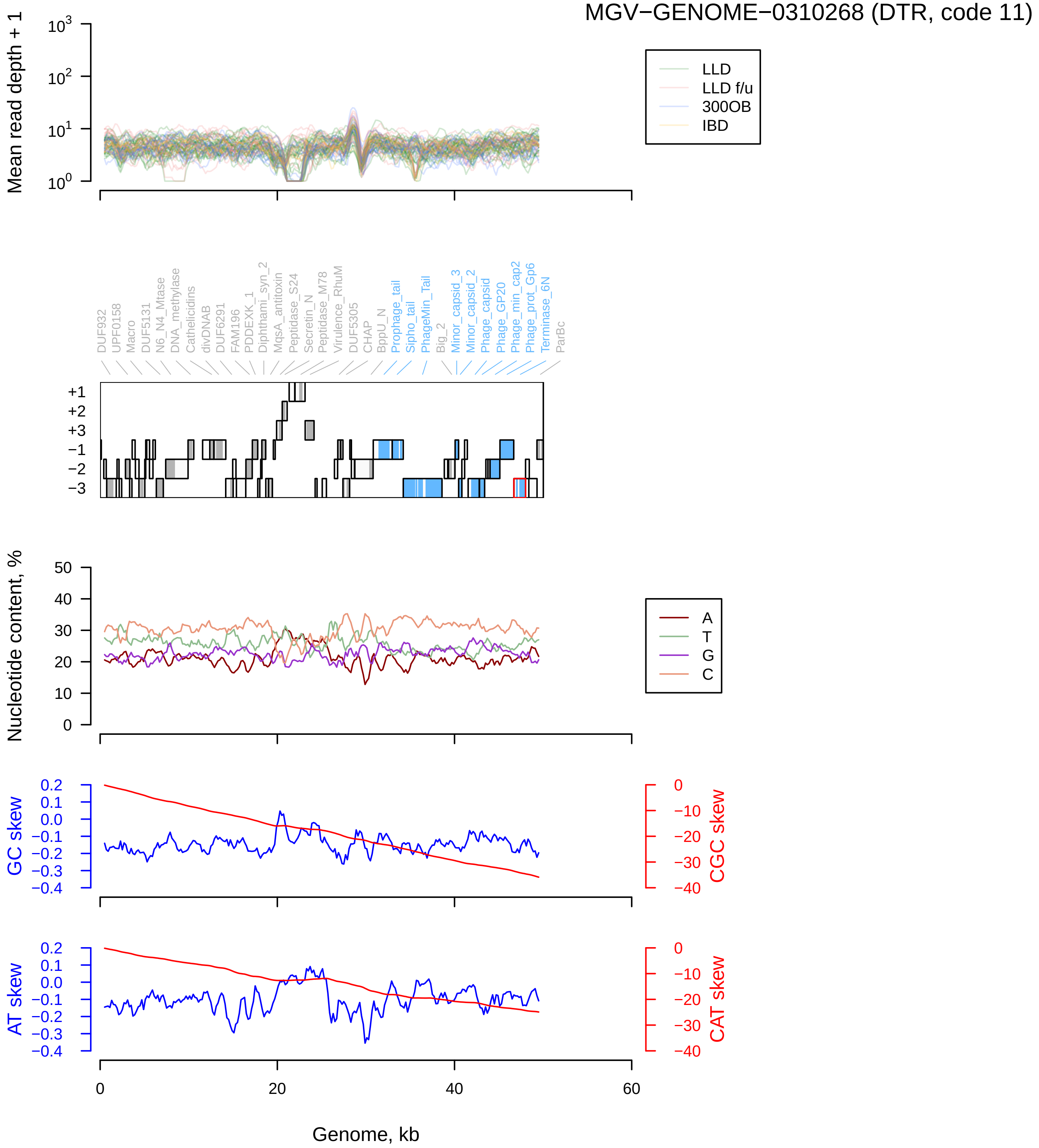
MGV−GENOME−0310268 (DTR, code 11)
Mean read depth + 1
LLD
LLD f/u
300OB
IBD
DUF932
UPF0158
Macro
DUF5131
N6_N4_Mtase
DNA_methylase
Cathelicidins
divDNAB
DUF6291
FAM196
PDDEXK_1
Diphthami_syn_2
MqsA_antitoxin
Peptidase_S24
Secretin_N
Peptidase_M78
Virulence_RhuM
DUF5305
CHAP
BppU_N
Prophage_tail
Sipho_tail
PhageMin_Tail
Big_2
Minor_capsid_3
Minor_capsid_2
Phage_capsid
Phage_GP20
Phage_min_cap2
Phage_prot_Gp6
Terminase_6N
ParBc
Nucleotide content, %
A
T
G
C
GC skew
CGC skew
AT skew
CAT skew
Genome, kb

#### MGV−GENOME−0321287 (DTR, code 11)

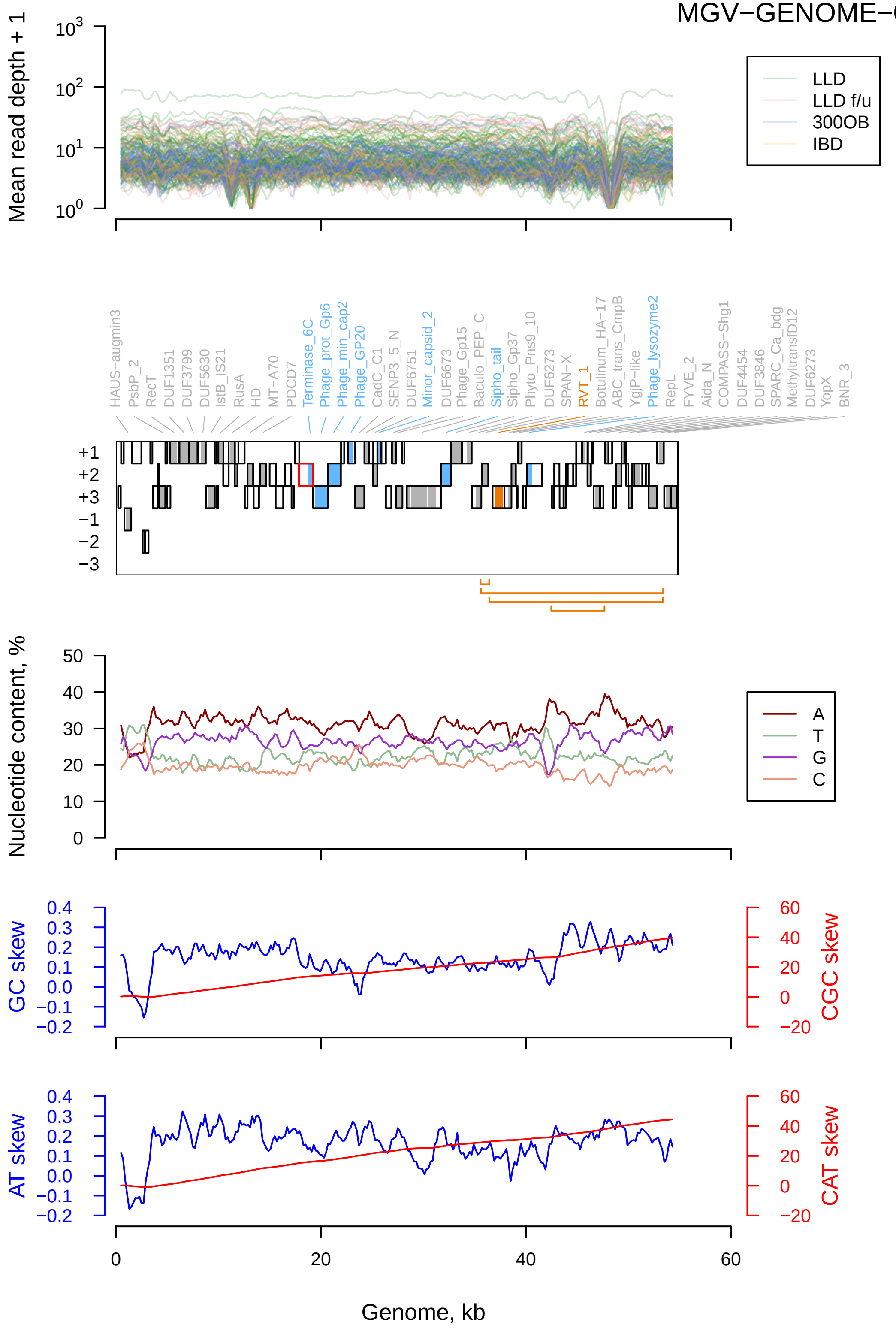

MGV−GENOME−0288753 (DTR, code 11)

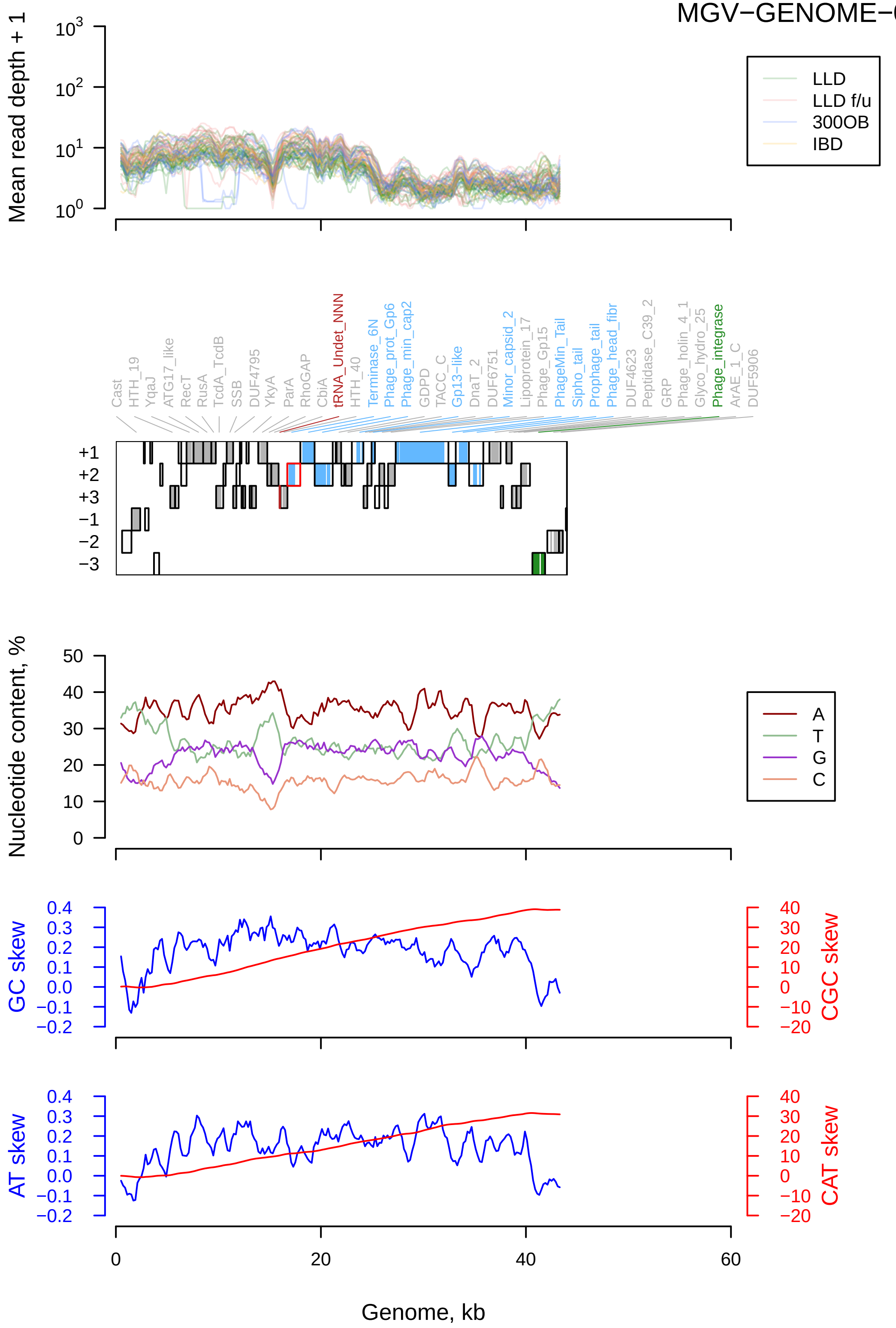

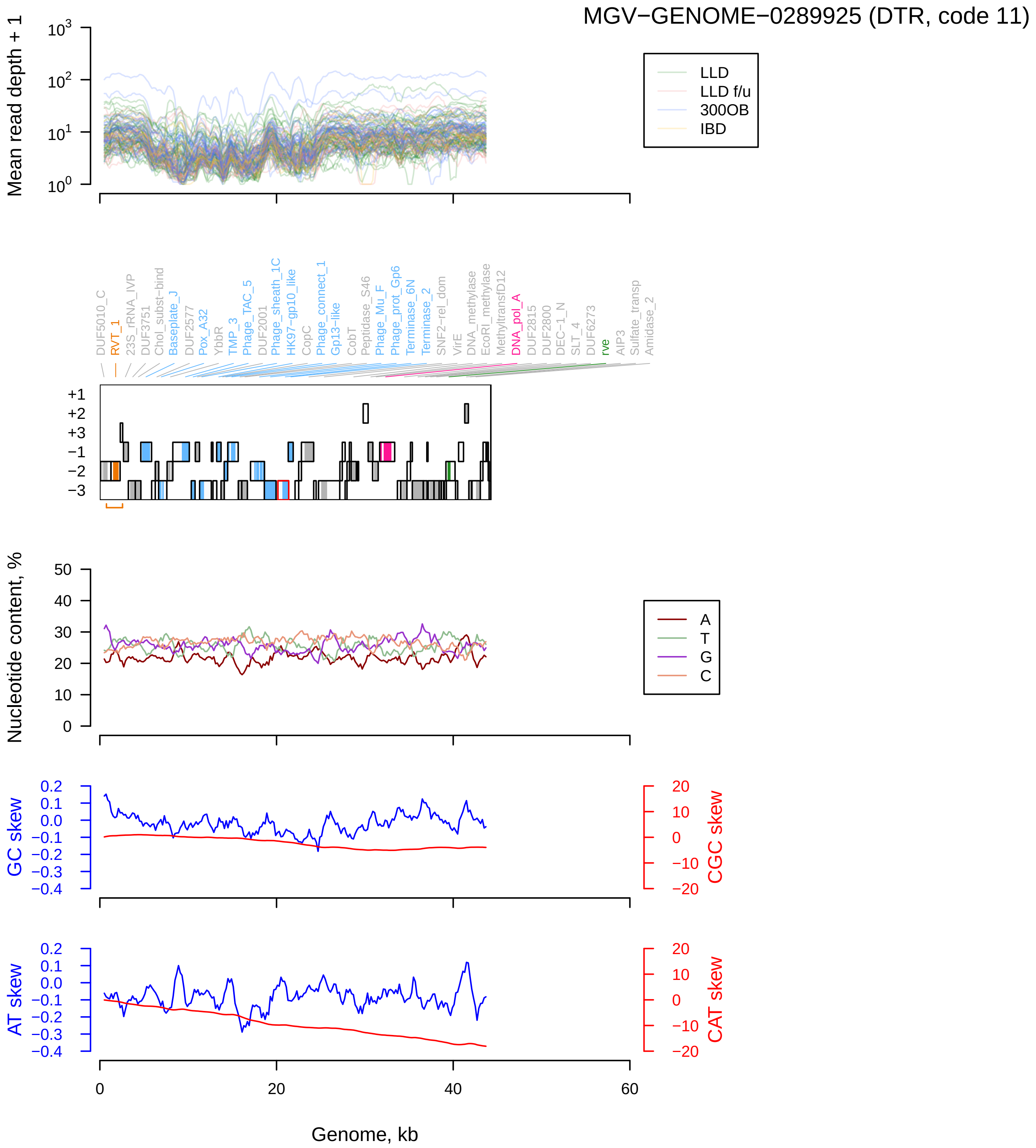

MGV−GENOME−0289925 (DTR, code 11)
Mean read depth + 1
LLD
LLD f/u
300OB
IBD
DUF5010_C
RVT_1
23S_rRNA_IVP
DUF3751
Chol_subst−bind
Baseplate_J
DUF2577
Pox_A32
YbbR
TMP_3
Phage_TAC_5
DUF2001
Phage_sheath_1C
HK97−gp10_like
CopC
Phage_connect_1
Gp13−like
CobT
Peptidase_S46
Phage_Mu_F
Phage_prot_Gp6
Terminase_6N
Terminase_2
SNF2−rel_dom
VirE
DNA_methylase
EcoRI_methylase
MethyltransfD12
DNA_pol_A
DUF2815
DUF2800
DEC−1_N
SLT_4
DUF6273
rve
AIP3
Sulfate_transp
Amidase_2
Nucleotide content, %
A
T
G
C
GC skew
CGC skew
AT skew
CAT skew
Genome, kb

NL_vir036031 (DTR, code 11)

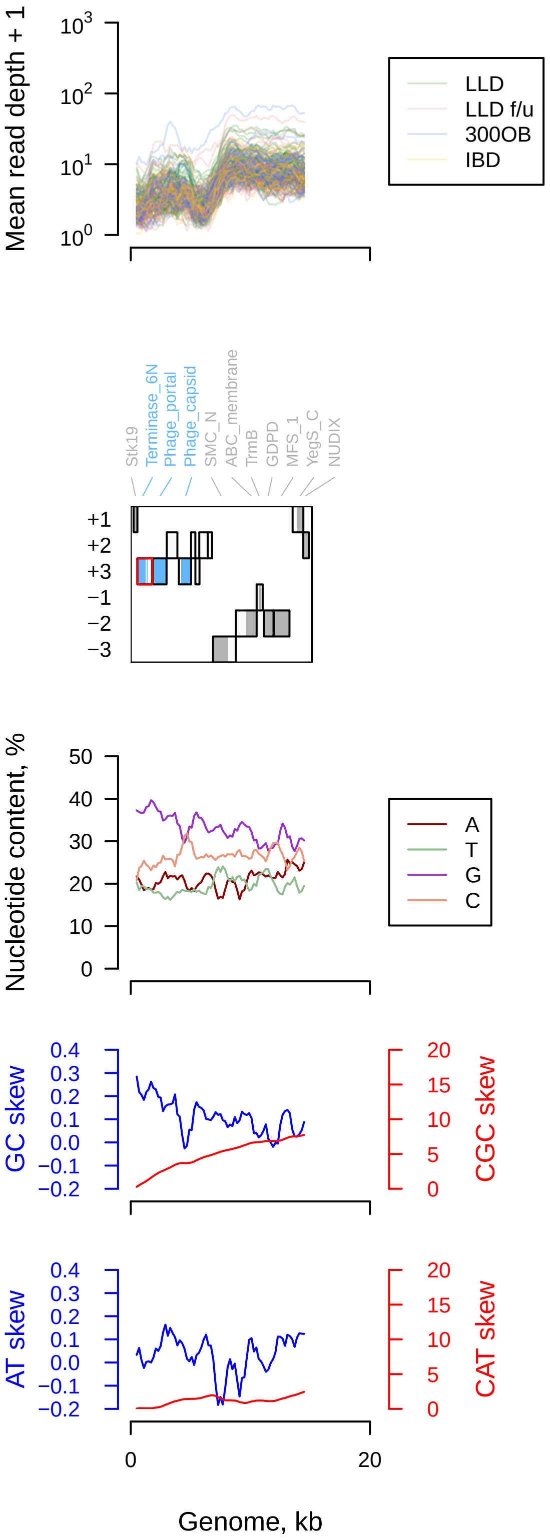

NL_vir053139 (DTR, code 11)

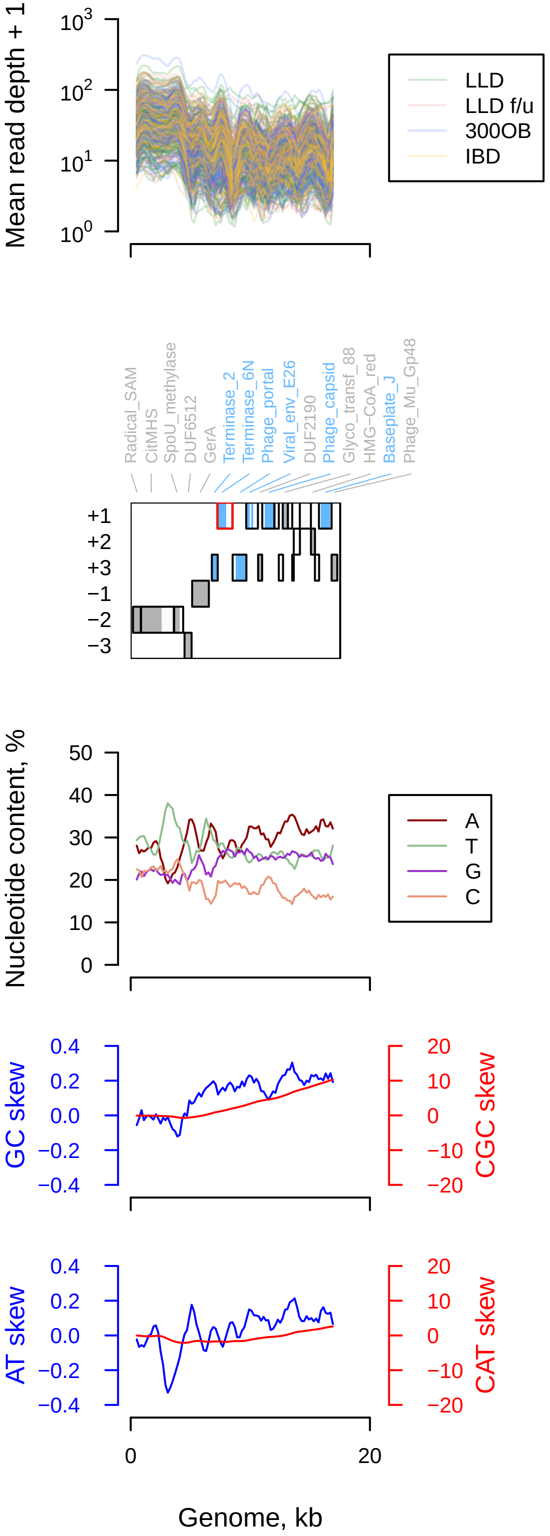

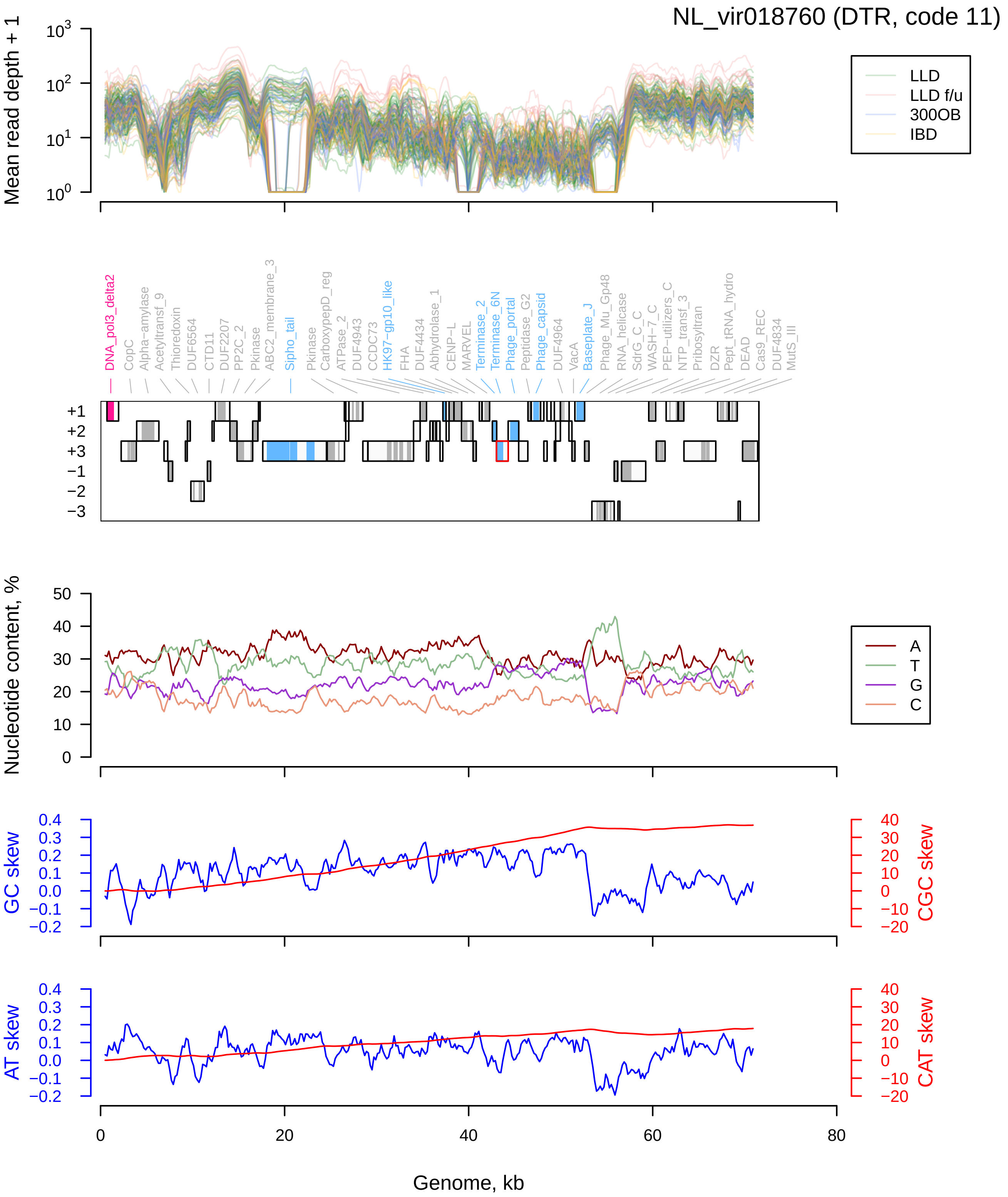
NL_vir018760 (DTR, code 11)
Mean read depth + 1
LLD
LLD f/u
300OB
IBD
DNA_pol3_delta2
CopC
Alpha−amylase
Acetyltransf_9
Thioredoxin
DUF6564
CTD11
DUF2207
PP2C_2
Pkinase
ABC2_membrane_3
Sipho_tail
Pkinase
CarboxypepD_reg
ATPase_2
DUF4943
CCDC73
HK97−gp10_like
FHA
DUF4434
Abhydrolase_1
CENP−L
MARVEL
Terminase_2
Terminase_6N
Phage_portal
Peptidase_G2
Phage_capsid
DUF4964
VacA
Baseplate_J
Phage_Mu_Gp48
RNA_helicase
SdrG_C_C
WASH−7_C
PEP−utilizers_C
NTP_transf_3
Pribosyltran
DZR
Pept_tRNA_hydro
DEAD
Cas9_REC
DUF4834
MutS_III
+1
+2
+3
−1
−2
−3
Nucleotide content, %
A
T
G
C
GC skew
CGC skew
AT skew
CAT skew
Genome, kb

### NL_vir026707 (DTR, code 11)

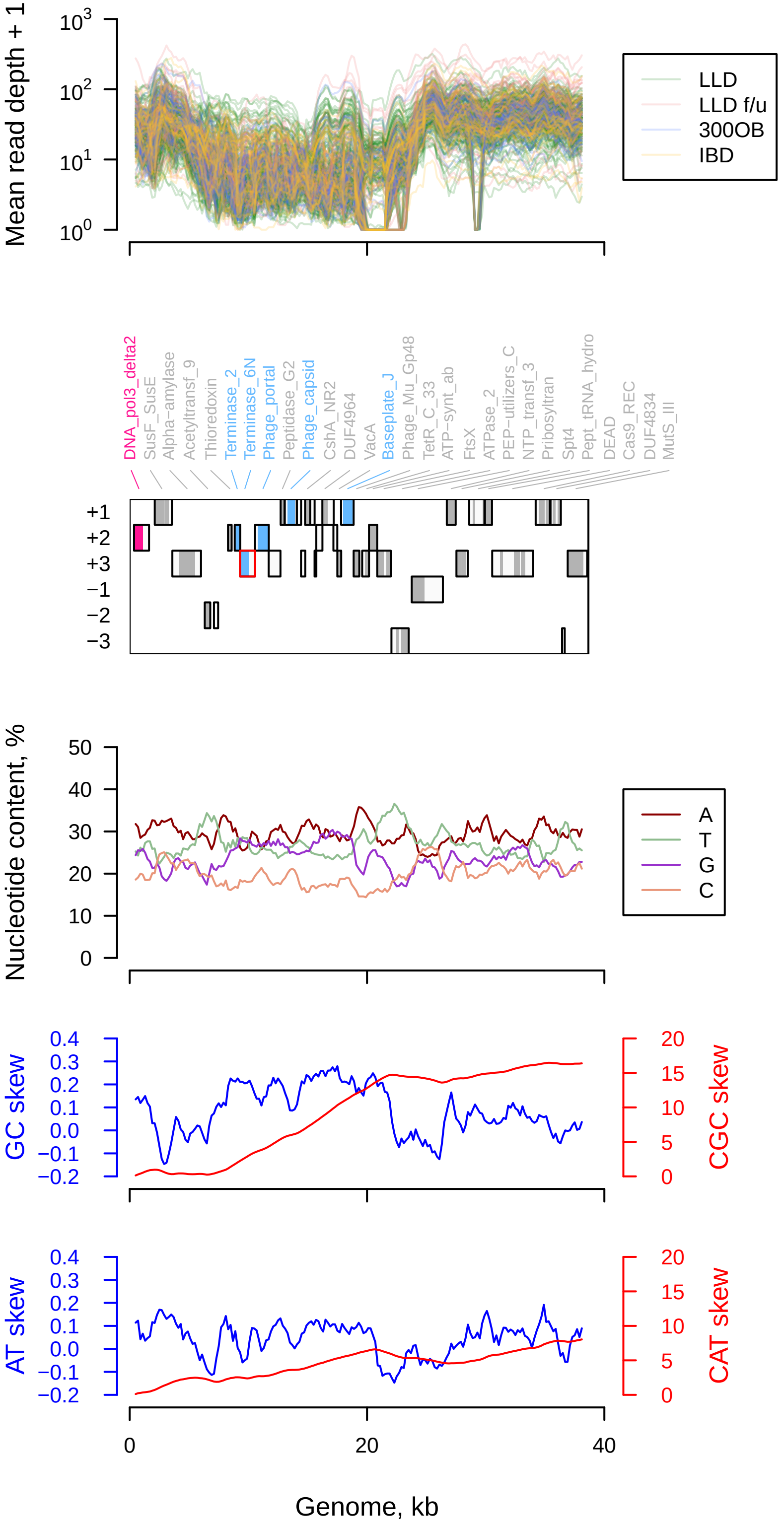

MGV−GENOME−0287672 (DTR, code 11)

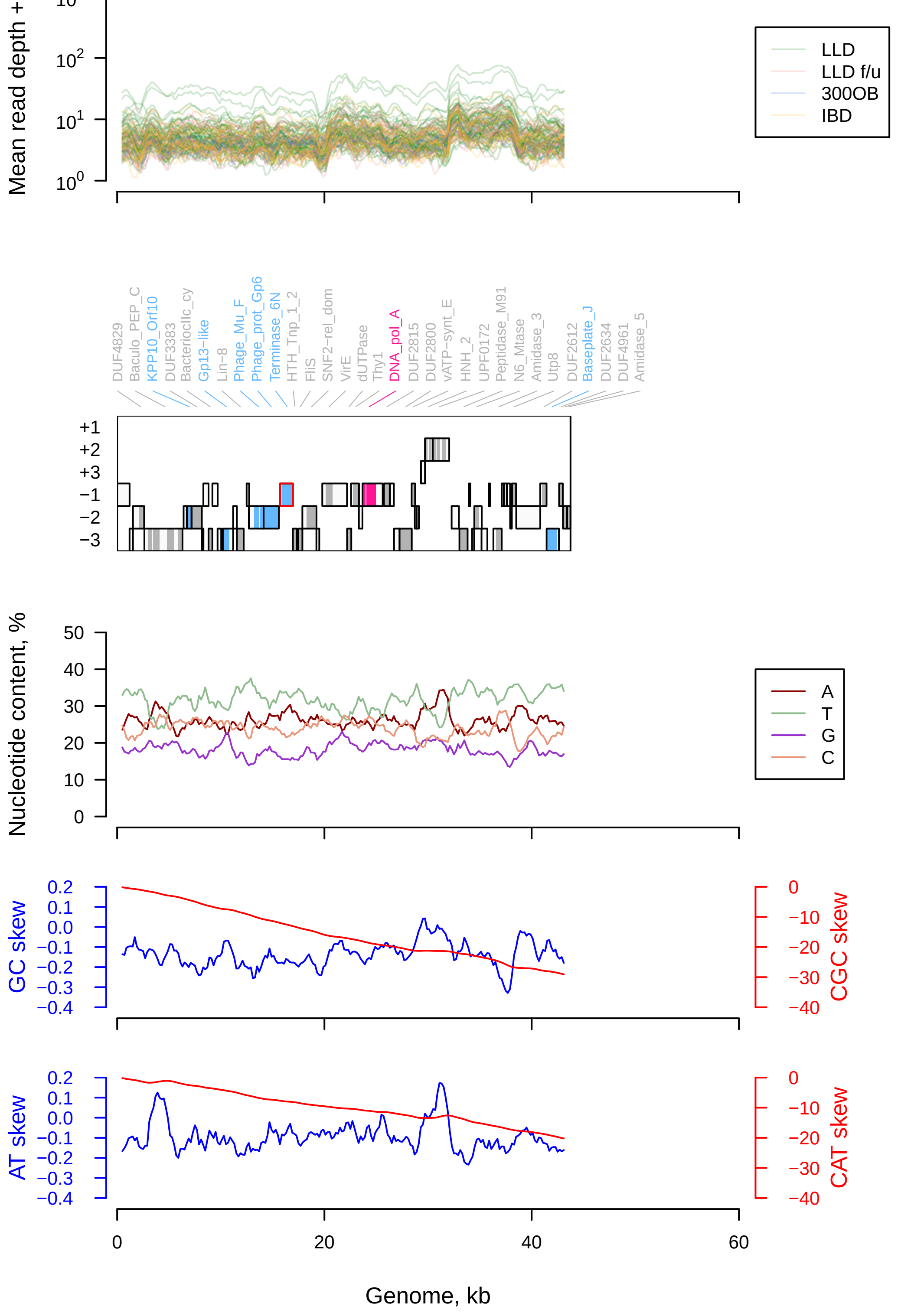

NL_vir014968 (DTR, code 11)

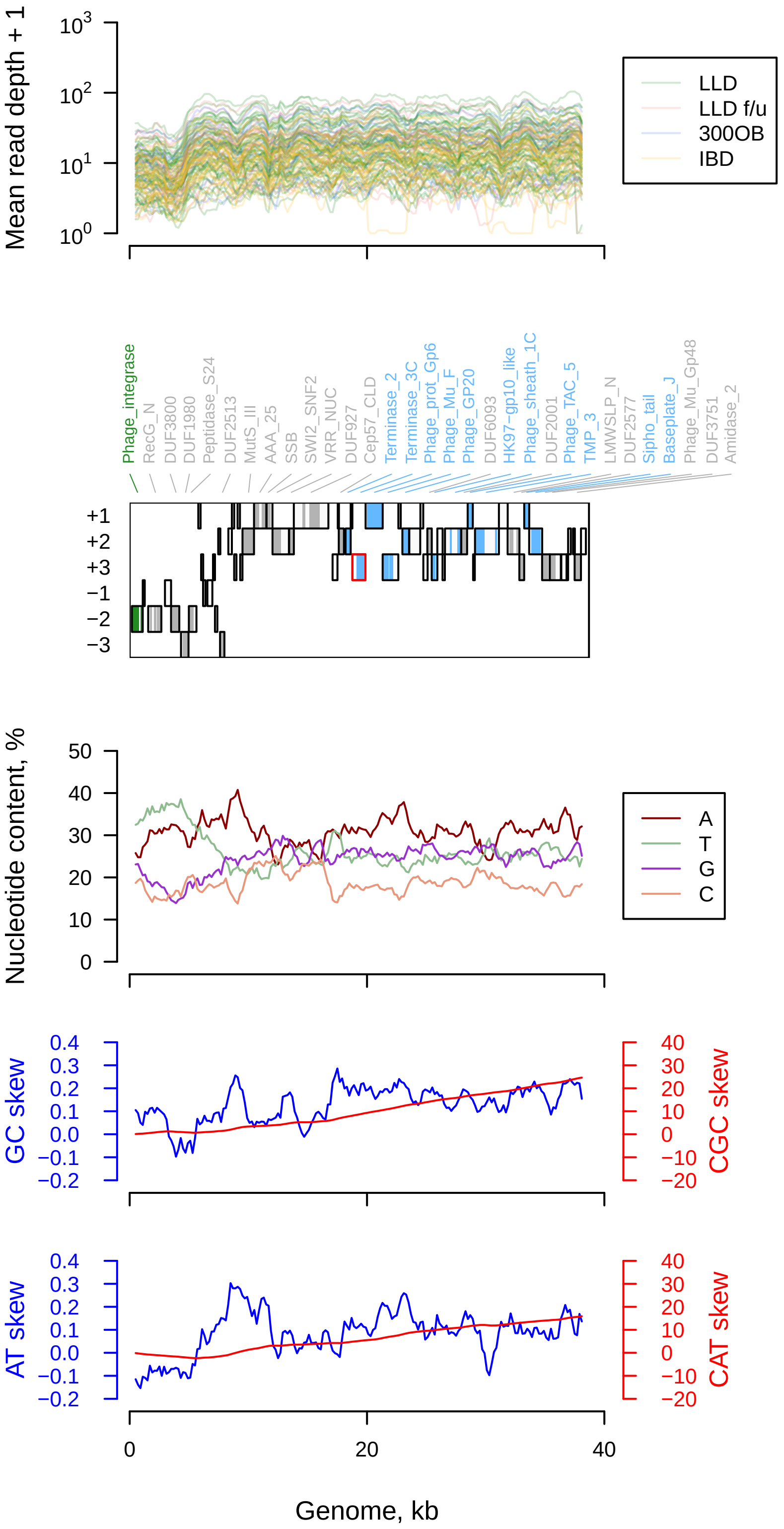

uvig_512347 (DTR, code 11)

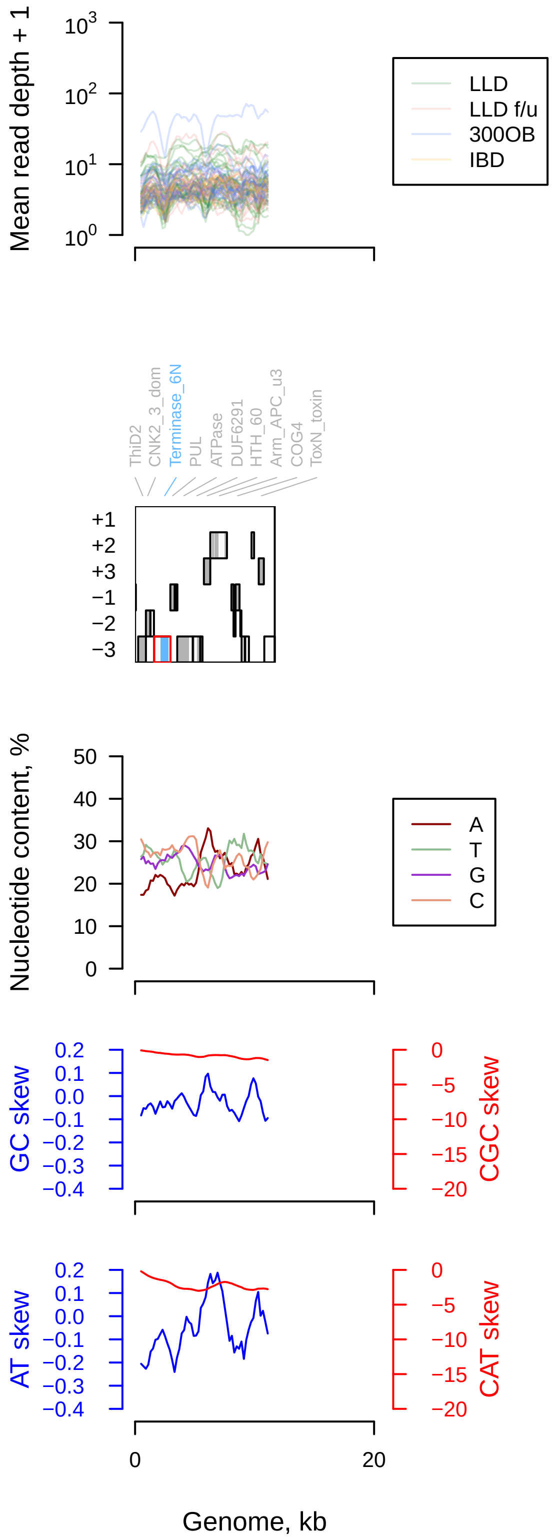

OLXK01000549.1 (DTR, code 11)

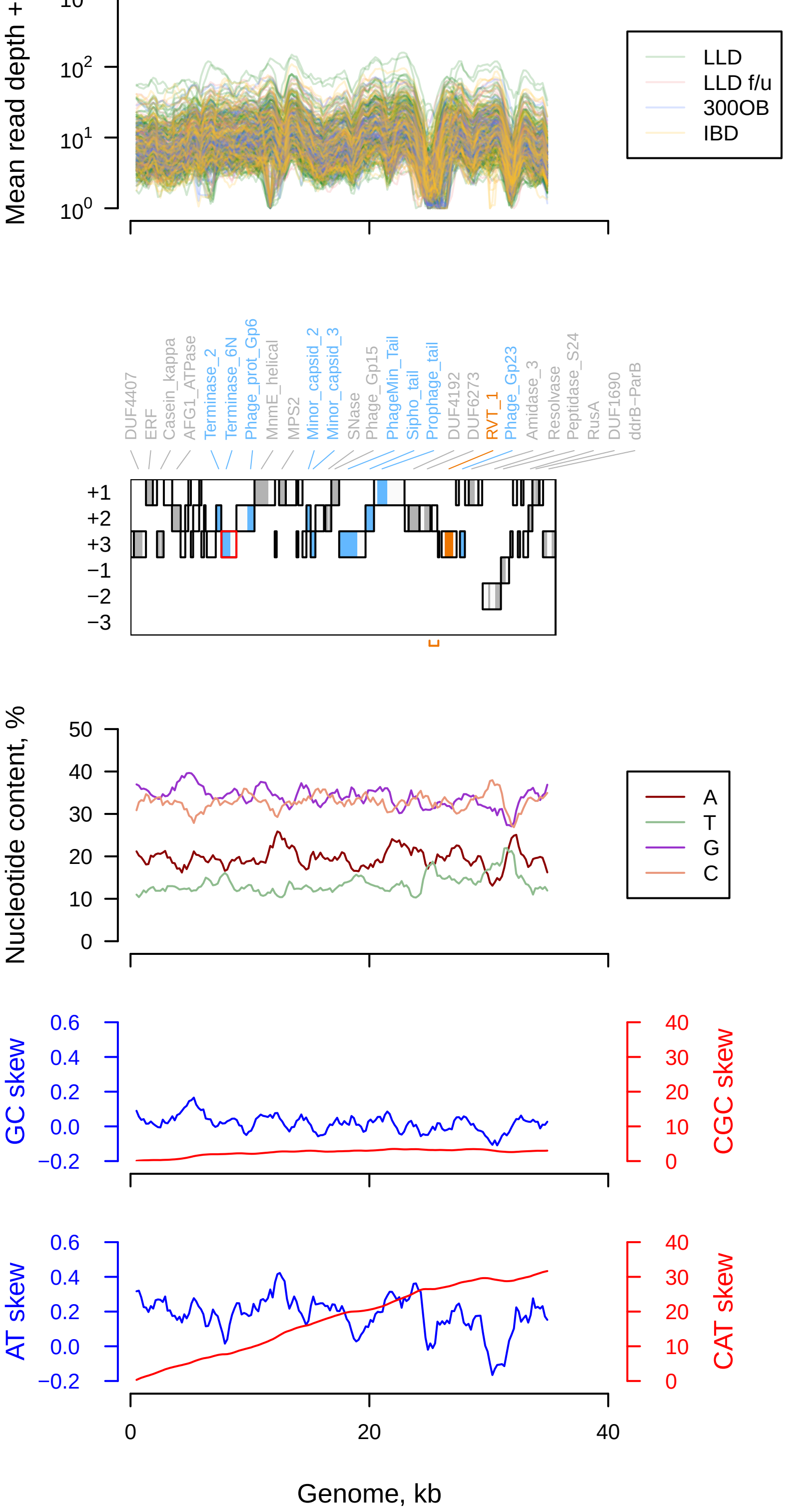

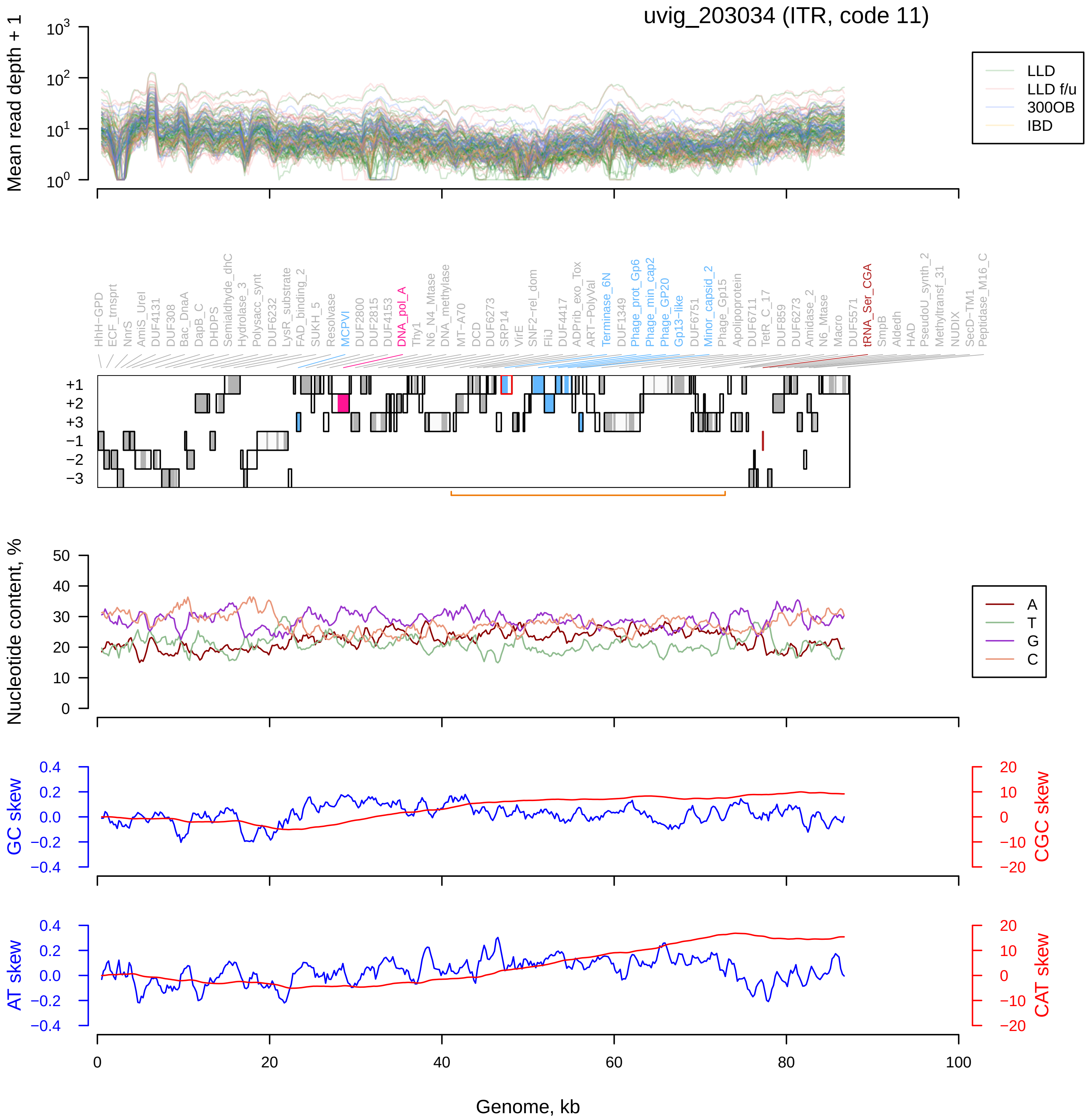
uvig_203034 (ITR, code 11)
Mean read depth + 1
LLD
LLD f/u
300OB
IBD
HhH−GPD
ECF_trnsprt
NnrS
AmiS_UreI
DUF4131
DUF308
Bac_DnaA
DapB_C
DHDPS
Semialdhyde_dhC
Hydrolase_3
Polysacc_synt
DUF6232
LysR_substrate
FAD_binding_2
SUKH_5
Resolvase
MCPVI
DUF2800
DUF2815
DUF4153
DNA_pol_A
Thy1
N6_N4_Mtase
DNA_methylase
MT−A70
DCD
DUF6273
SRP14
VirE
SNF2−rel_dom
FliJ
DUF4417
ADPrib_exo_Tox
ART−PolyVal
Terminase_6N
DUF1349
Phage_prot_Gp6
Phage_min_cap2
Phage_GP20
Gp13−like
DUF6751
Minor_capsid_2
Phage_Gp15
Apolipoprotein
DUF6711
TetR_C_17
DUF859
DUF6273
Amidase_2
N6_Mtase
Macro
DUF5571
tRNA_Ser_CGA
SmpB
Aldedh
HAD
PseudoU_synth_2
Methyltransf_31
NUDIX
SecD−TM1
Peptidase_M16_C
+1
+2
+3
−1
−2
−3
Nucleotide content, %
A
T
G
C
GC skew
CGC skew
AT skew
CAT skew
Genome, kb

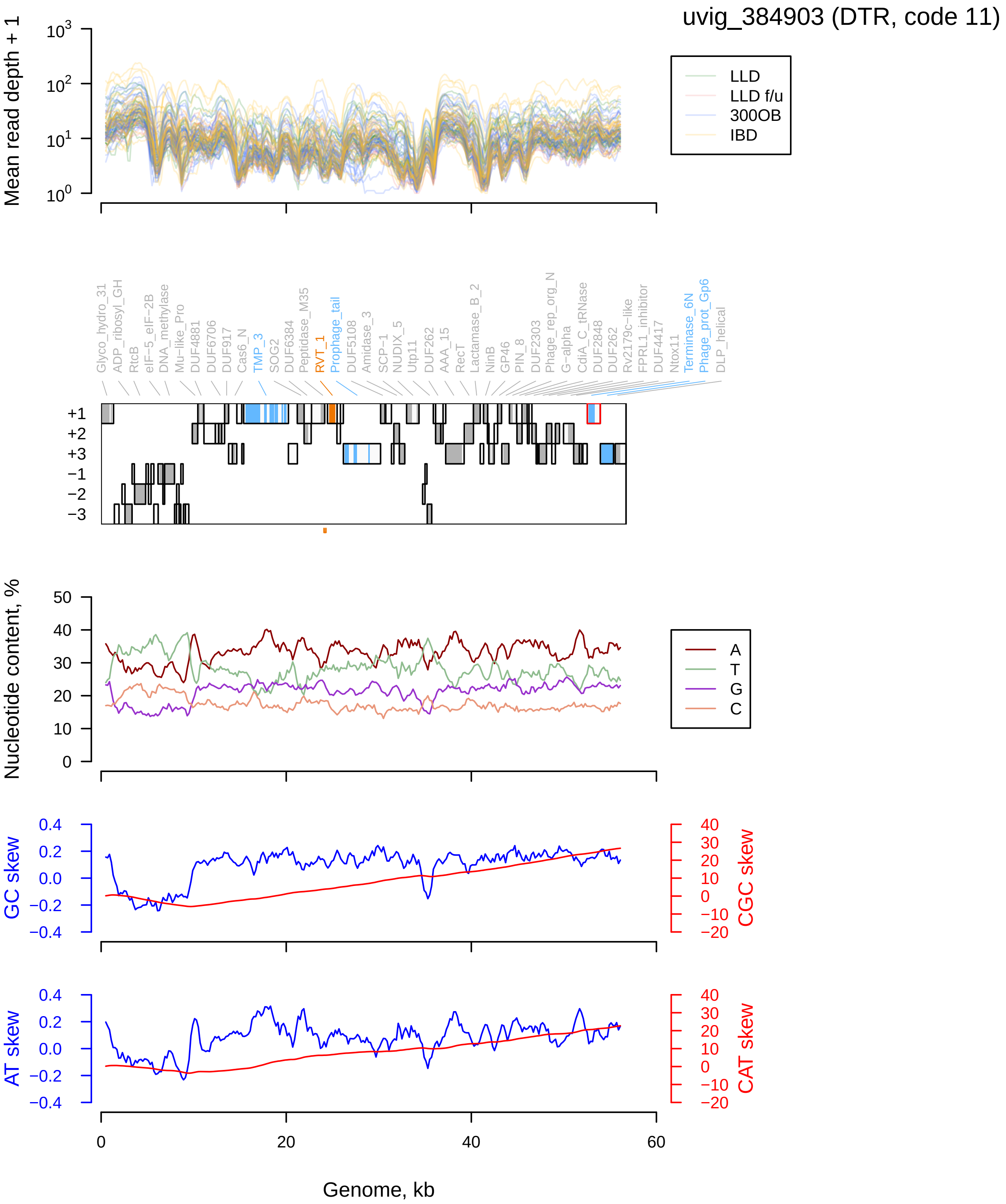
uvig_384903 (DTR, code 11)
Mean read depth + 1
LLD
LLD f/u
300OB
IBD
Glyco_hydro_31
ADP_ribosyl_GH
RtcB
eIF−5_eIF−2B
DNA_methylase
Mu−like_Pro
DUF4881
DUF6706
DUF917
Cas6_N
TMP_3
SOG2
DUF6384
Peptidase_M35
RVT_1
Prophage_tail
DUF5108
Amidase_3
SCP−1
NUDIX_5
Utp11
DUF262
AAA_15
RecT
Lactamase_B_2
NinB
GP46
PIN_8
DUF2303
Phage_rep_org_N
G−alpha
CdiA_C_tRNase
DUF2848
DUF262
Rv2179c−like
FPRL1_inhibitor
DUF4417
Ntox11
Terminase_6N
Phage_prot_Gp6
DLP_helical
Nucleotide content, %
A
T
G
C
GC skew
CGC skew
AT skew
CAT skew
Genome, kb

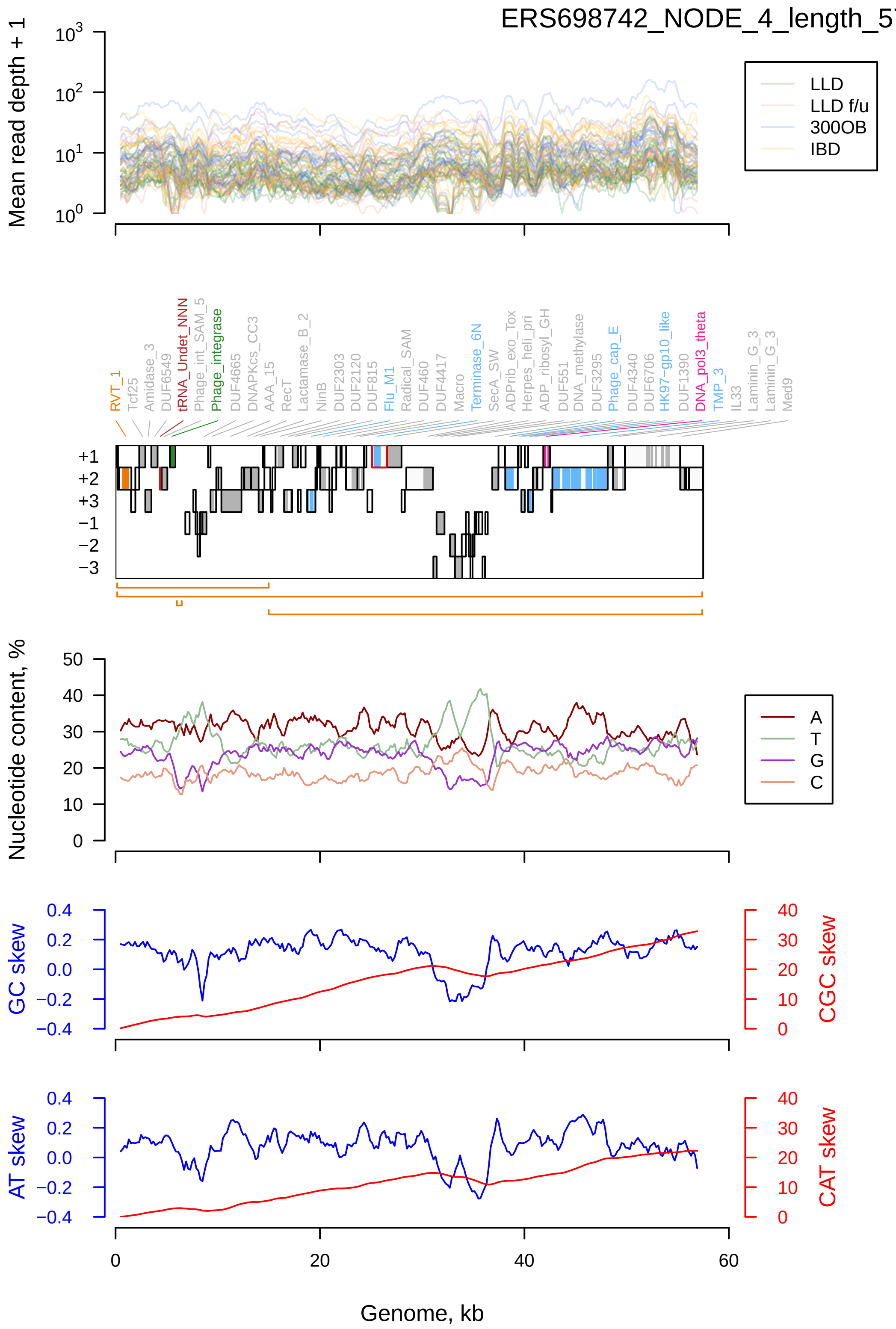

ERS698742_NODE_4_length_57489_cov_840.739161 (DTR, code 11)
Mean read depth + 1
LLD
LLD f/u
300OB
IBD
RVT_1
Tcf25
Amidase_3
DUF6549
tRNA_Undet_NNN
Phage_int_SAM_5
Phage_integrase
DUF4665
DNAPKcs_CC3
AAA_15
RecT
Lactamase_B_2
NinB
DUF2303
DUF2120
DUF815
Flu_M1
Radical_SAM
DUF460
DUF4417
Macro
Terminase_6N
SecA_SW
ADPrib_exo_Tox
Herpes_heli_pri
ADP_ribosyl_GH
DUF551
DNA_methylase
DUF3295
Phage_cap_E
DUF4340
DUF6706
HK97-gp10_like
DUF1390
DNA_pol3_theta
TMP_3
IL33
Laminin_G_3
Laminin_G_3
Med9
+1
+2
+3
−1
−2
−3
Nucleotide content, %
A
T
G
C
GC skew
CGC skew
AT skew
CAT skew
Genome, kb

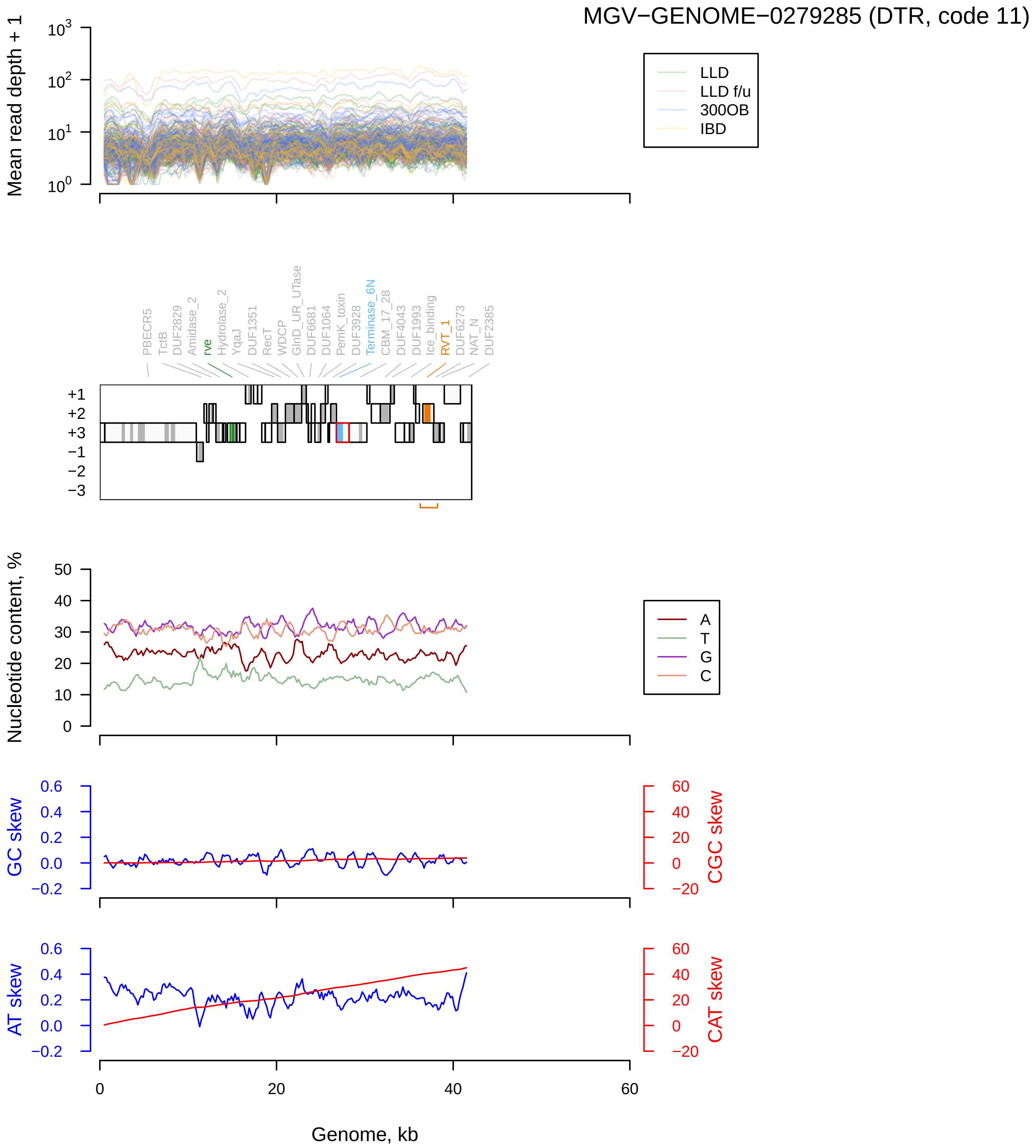

MGV−GENOME−0279285 (DTR, code 11)
Mean read depth + 1
LLD
LLD f/u
300OB
IBD
PBECR5
TctB
DUF2829
Amidase_2
rve
Hydrolase_2
YqaJ
DUF1351
RecT
WDCP
GlnD_UR_UTase
DUF6681
DUF1064
PemK_toxin
DUF3928
Terminase_6N
CBM_17_28
DUF4043
DUF1993
Ice_binding
RVT_1
DUF6273
NAT_N
DUF2385
Nucleotide content, %
A
T
G
C
GC skew
CGC skew
AT skew
CAT skew
Genome, kb

MGV−GENOME−0248858 (DTR, code 11)

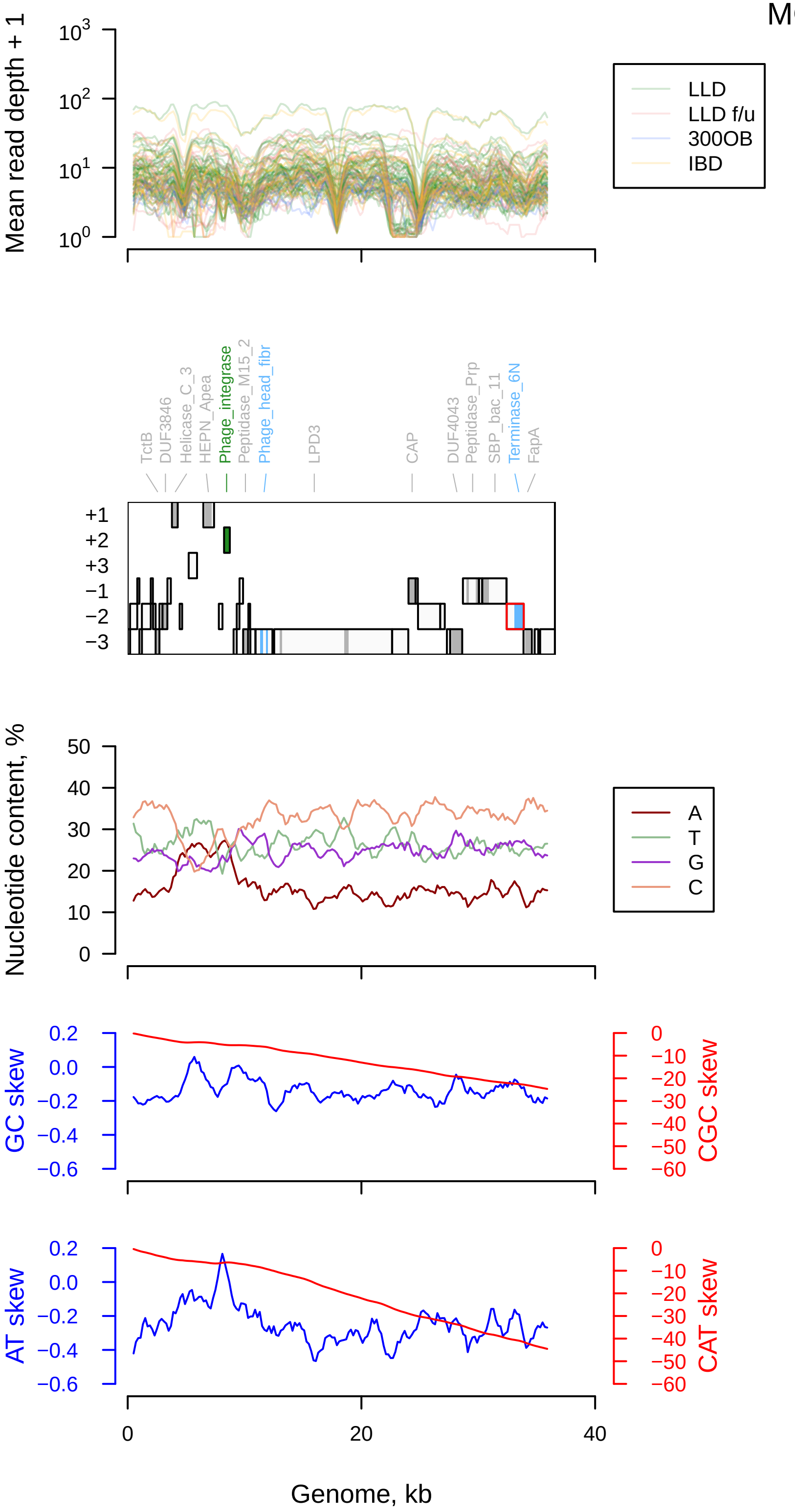

NL_vir049144 (DTR, code 11)

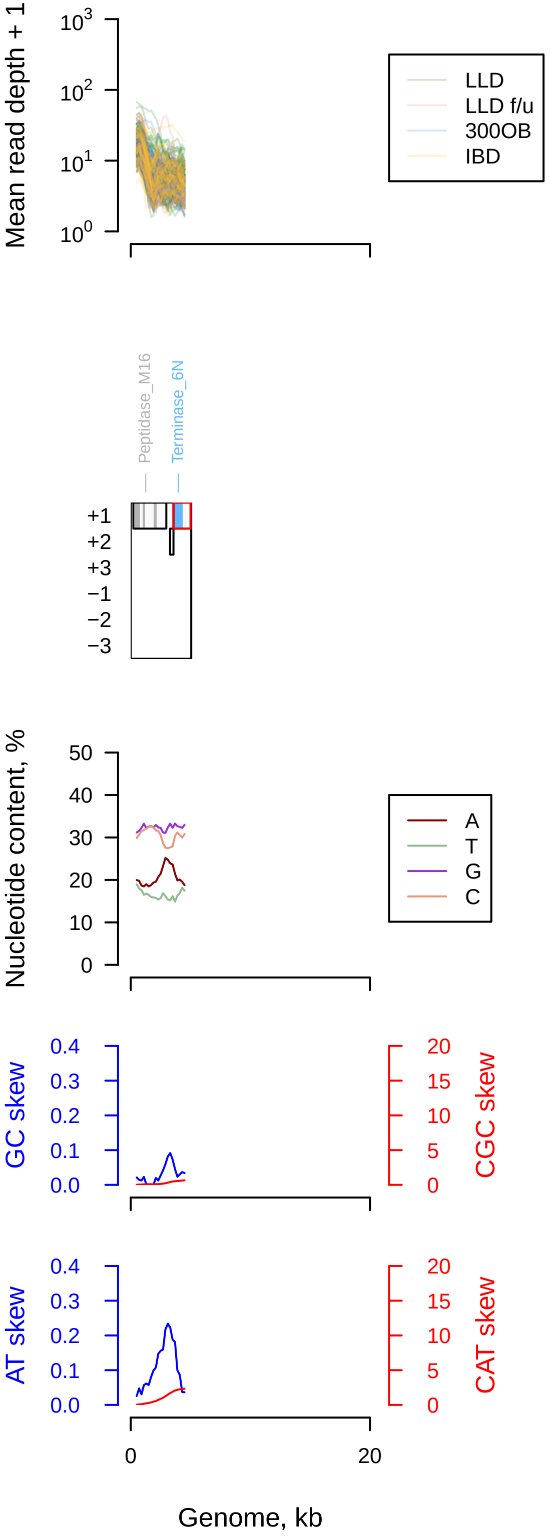

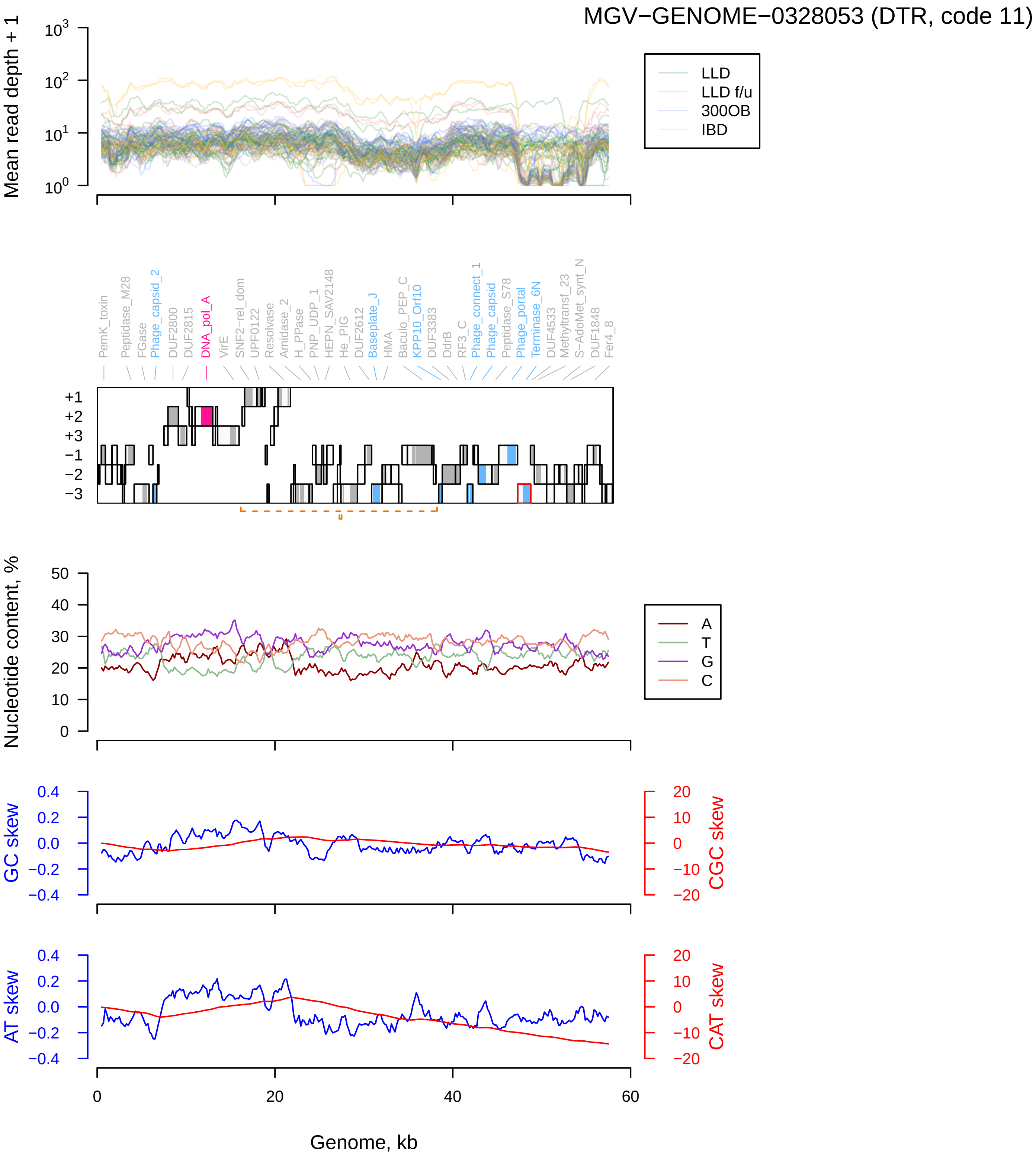
MGV−GENOME−0328053 (DTR, code 11)
Mean read depth + 1
LLD
LLD f/u
300OB
IBD
PemK_toxin
Peptidase_M28
FGase
Phage_capsid_2
DUF2800
DUF2815
DNA_pol_A
VirE
SNF2−rel_dom
UPF0122
Resolvase
Amidase_2
H_PPase
PNP_UDP_1
HEPN_SAV2148
He_PIG
DUF2612
Baseplate_J
HMA
Baculo_PEP_C
KPP10_Orf10
DUF3383
DdrB
RF3_C
Phage_connect_1
Phage_capsid
Peptidase_S78
Phage_portal
Terminase_6N
DUF4533
Methyltransf_23
S−AdoMet_synt_N
DUF1848
Fer4_8
+1
+2
+3
−1
−2
−3
Nucleotide content, %
A
T
G
C
GC skew
CGC skew
AT skew
CAT skew
Genome, kb

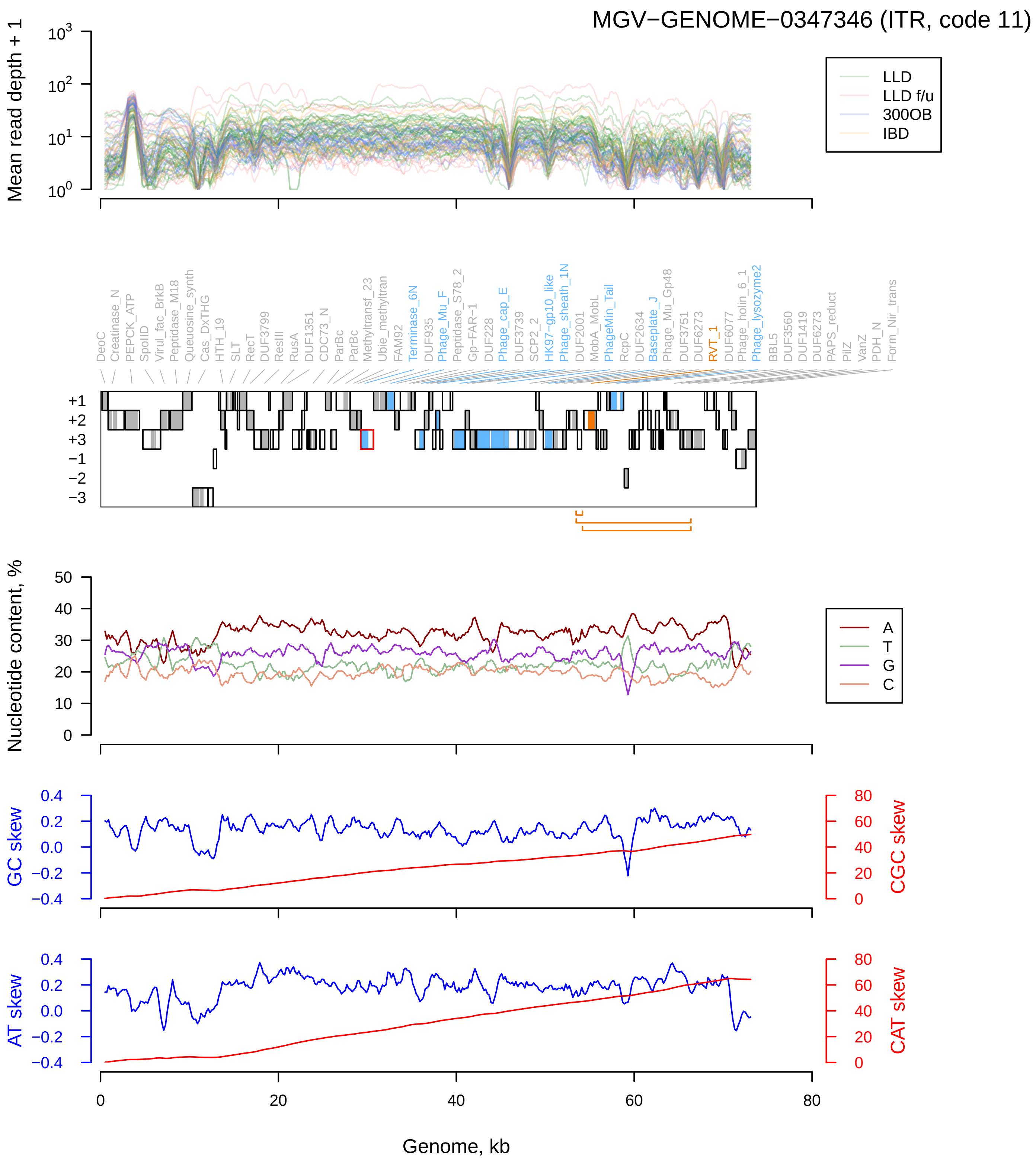
MGV−GENOME−0347346 (ITR, code 11)
Mean read depth + 1
LLD
LLD f/u
300OB
IBD
DeoC
Creatinase_N
PEPCK_ATP
SpoIID
Virul_fac_BrkB
Peptidase_M18
Queuosine_synth
Cas_DxTHG
HTH_19
SLT
RecT
DUF3799
ResIII
RusA
DUF1351
CDC73_N
ParBc
ParBc
Methyltransf_23
Ubie_methyltran
FAM92
Terminase_6N
DUF935
Phage_Mu_F
Peptidase_S78_2
Gp-FAR-1
DUF228
Phage_cap_E
DUF3739
SCP2_2
HK97-gp10_like
Phage_sheath_1N
DUF2001
MobA_MobL
PhageMin_Tail
RcpC
DUF2634
Baseplate_J
Phage_Mu_Gp48
DUF3751
DUF6273
RVT_1
DUF6077
Phage_holin_6_1
Phage_lysozyme2
BBL5
DUF3560
DUF1419
DUF6273
PAPS_reduct
PilZ
VanZ
PDH_N
Form_Nir_trans
+1
+2
+3
−1
−2
−3
Nucleotide content, %
A
T
G
C
GC skew
CGC skew
AT skew
CAT skew
Genome, kb
