## Supplementary material for "Diversity and ecology of *Caudoviricetes* phages with genome terminal repeats in fecal metagenomes from four Dutch cohorts": Material S2

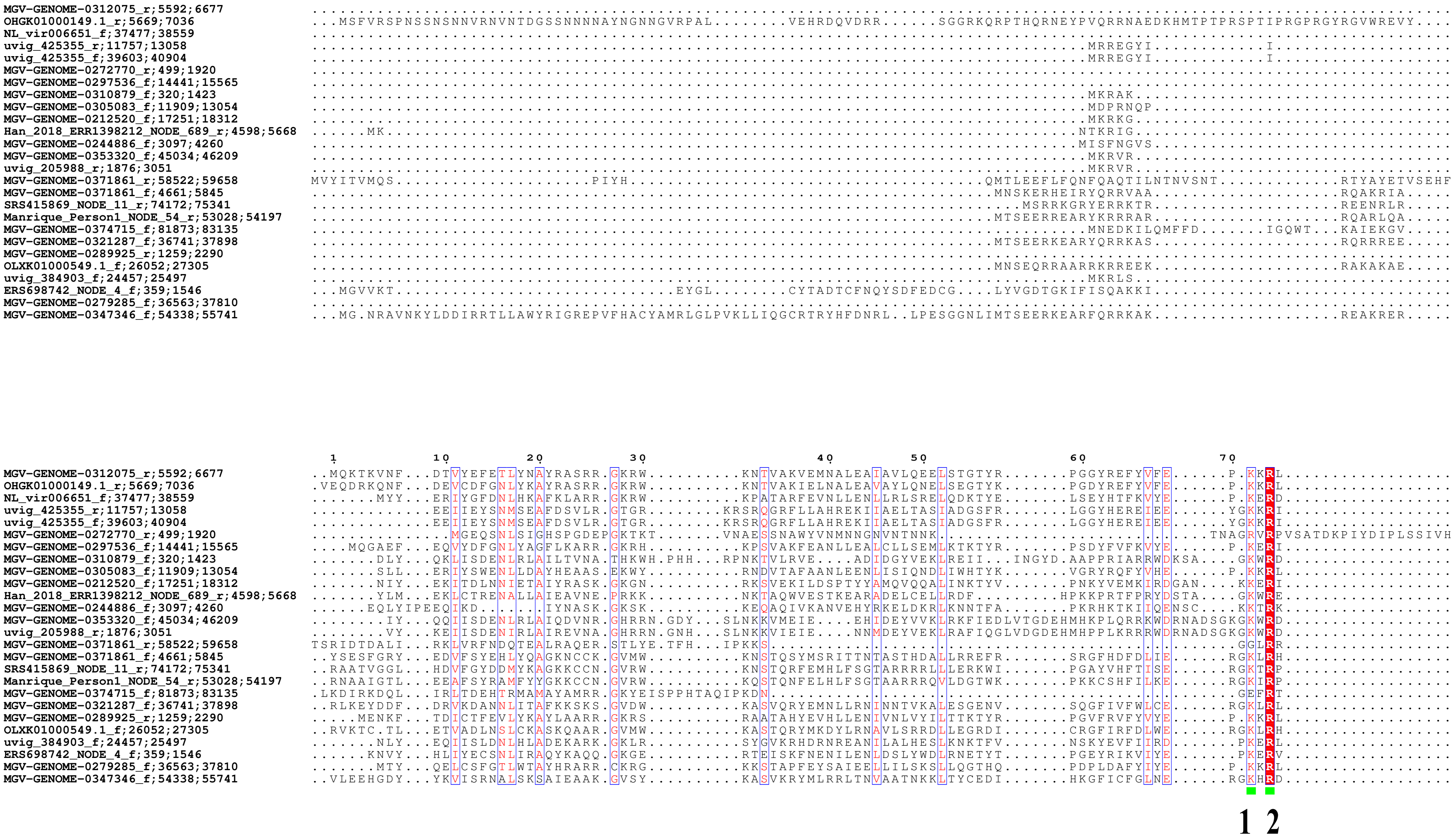
MGV-GENOME-0312075_r;5592;6677
OHGK01000149.1_r;5669;7036
NL_vir006651_f;37477;38559
uvig_425355_r;11757;13058
uvig_425355_f;39603;40904
MGV-GENOME-0272770_r;499;1920
MGV-GENOME-0297536_f;14441;15565
MGV-GENOME-0310879_f;320;1423
MGV-GENOME-0305083_f;11909;13054
MGV-GENOME-0212520_f;17251;18312
Han_2018_ERR1398212_NODE_689_r;4598;5668
MGV-GENOME-0244886_f;3097;4260
MGV-GENOME-0353320_f;45034;46209
uvig_205988_r;1876;3051
MGV-GENOME-0371861_r;58522;59658
MGV-GENOME-0371861_f;4661;5845
SRS415869_NODE_11_r;74172;75341
Manrique_Person1_NODE_54_r;53028;54197
MGV-GENOME-0374715_f;81873;83135
MGV-GENOME-0321287_f;36741;37898
MGV-GENOME-0289925_r;1259;2290
OLXK01000549.1_f;26052;27305
uvig_384903_f;24457;25497
ERS698742_NODE_4_f;359;1546
MGV-GENOME-0279285_f;36563;37810
MGV-GENOME-0347346_f;54338;55741
1 10 20 30 40 50 60 70
1 2

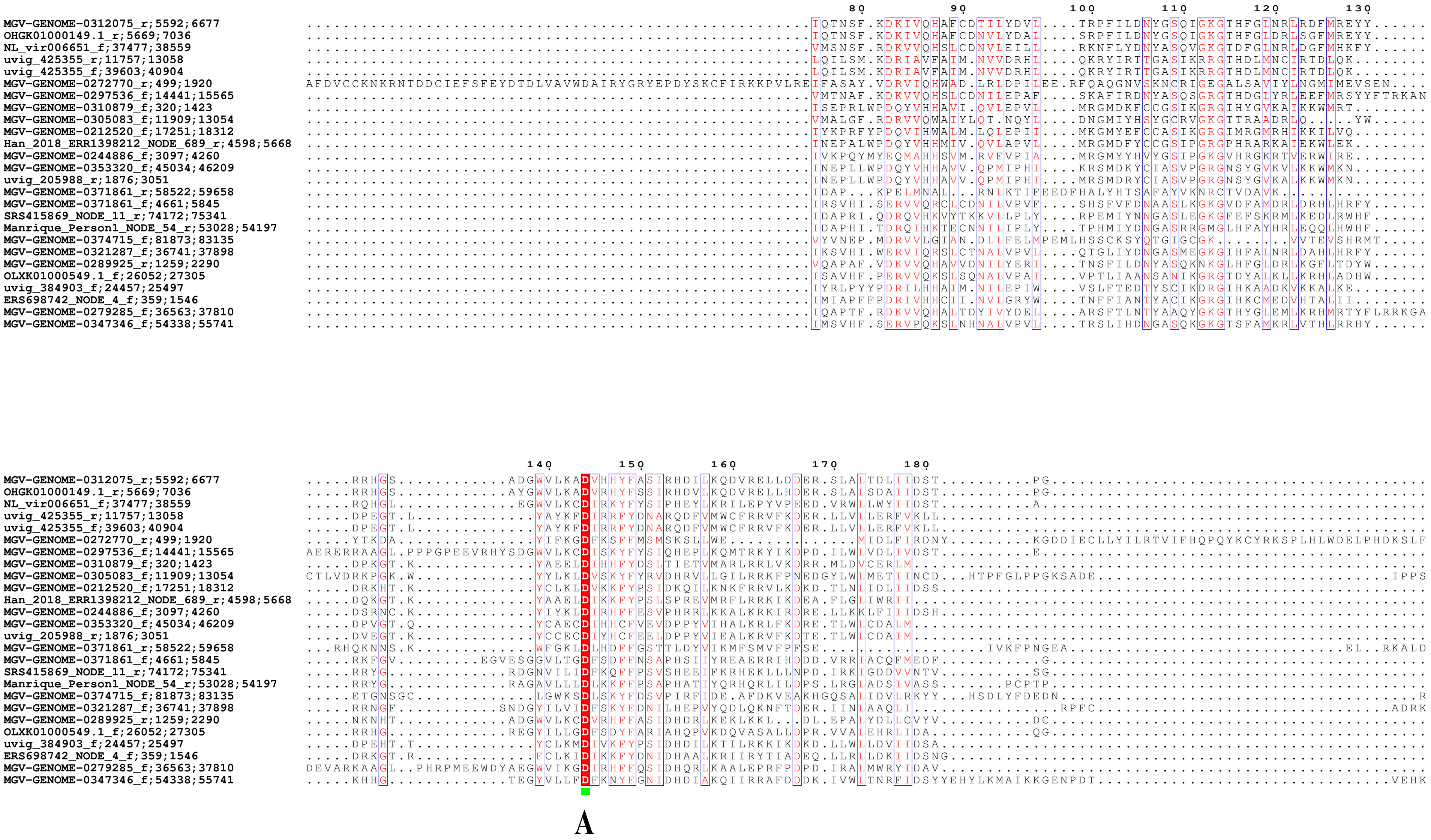
80
90
100
110
120
130
140
150
160
170
180
MGV-GENOME-0312075_r;5592;6677
OHGK01000149.1_r;5669;7036
NL_vir006651_f;37477;38559
uvig_425355_r;11757;13058
uvig_425355_f;39603;40904
MGV-GENOME-0272770_r;499;1920
MGV-GENOME-0297536_f;14441;15565
MGV-GENOME-0310879_f;320;1423
MGV-GENOME-0305083_f;11909;13054
MGV-GENOME-0212520_f;17251;18312
Han_2018_ERR1398212_NODE_689_r;4598;5668
MGV-GENOME-0244886_f;3097;4260
MGV-GENOME-0353320_f;45034;46209
uvig_205988_r;1876;3051
MGV-GENOME-0371861_r;58522;59658
MGV-GENOME-0371861_f;4661;5845
SRS415869_NODE_11_r;74172;75341
Manrique_Person1_NODE_54_r;53028;54197
MGV-GENOME-0374715_f;81873;83135
MGV-GENOME-0321287_f;36741;37898
MGV-GENOME-0289925_r;1259;2290
OLXK01000549.1_f;26052;27305
uvig_384903_f;24457;25497
ERS698742_NODE_4_f;359;1546
MGV-GENOME-0279285_f;36563;37810
MGV-GENOME-0347346_f;54338;55741
A

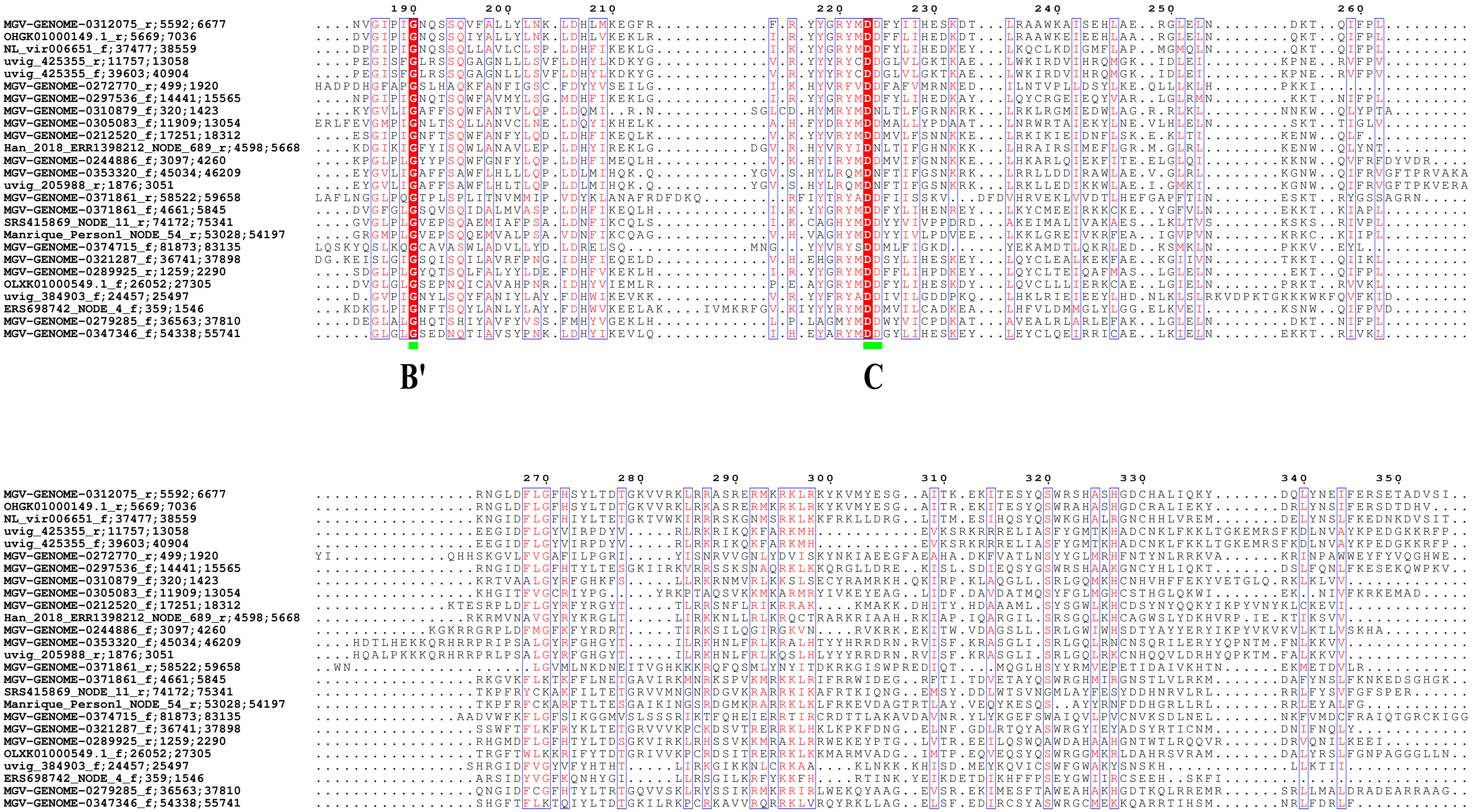
B'
C

```
                                                             360
MGV-GENOME-0312075_r;5592;6677           ...................NQRPAR..............................................................................................
OHGK01000149.1_r;5669;7036               .......................................................................................................................
NL_vir006651_f;37477;38559               ...................VESADRGESQVR........................................................................................
uvig_425355_r;11757;13058                ..................GVVV.....SIRE..LVNLPIVVK.DFETGI.....KTEQGEDRCIVAIEVNGEAKKFFTNSEEMKNILAQVKEMPDGFPFETTIKTETFGKGRTKYV.FT
uvig_425355_f;39603;40904                ..................GVVV.....SIRE..LVNLPIVVK.DFETGI.....KTEQGEDRCIVAIEVNGEAKKFFTNSEEMKNILAQVKEMPDGFPFETTIKTETFGKGRTKYV.FT
MGV-GENOME-0272770_r;499;1920            ...................VFVLKNEYNFKKQ..................................................................LKKQIRKGNAKKYLTPEIG..
MGV-GENOME-0297536_f;14441;15565         .......................................................................................................................
MGV-GENOME-0310879_f;320;1423            .............................RE.......................HTRKEQARWNM.................................CTEPLKSTA............
MGV-GENOME-0305083_f;11909;13054         ...................IELPQEVTQWELNQF.....................................................................................
MGV-GENOME-0212520_f;17251;18312         .............................RN.......................ESKKSRKH..............................................NKT........
Han_2018_ERR1398212_NODE_689_r;4598;5668 .............................RE.......................ASRKERILC........................................SST.............
MGV-GENOME-0244886_f;3097;4260           ........................RKENERN.......................GMVRSRKHC....................................RRETKRAGYNKQPAPCI...
MGV-GENOME-0353320_f;45034;46209         .............................RR.......................ECRRLQRLY.....................................PPYRAA.............
uvig_205988_r;1876;3051                  .............................RK.......................ECRRLQALY.....................................PPYQAA.............
MGV-GENOME-0371861_r;58522;59658         ..................LIK.....DDLR.........................................................................................
MGV-GENOME-0371861_f;4661;5845           ...................VS......EERQ.......................................................................LARAC............
SRS415869_NODE_11_r;74172;75341          ...................IE......NFRE.......................RGKKDEICC.....................................A..................
Manrique_Person1_NODE_54_r;53028;54197   .......................................................GLK....C.....................................SELSKTAAASA........
MGV-GENOME-0374715_f;81873;83135         LGYVRTKPDGCIVRGRGRNVKA.NRDKTDRD.....IPGYLTV...........GCMRNALLT........................................SRAVYNTLVASL....
MGV-GENOME-0321287_f;36741;37898         ...................VPA....GEWR.........................................................................................
MGV-GENOME-0289925_r;1259;2290           .......................................................................................................................
OLXK01000549.1_f;26052;27305             ..................AIQ......TWRRHGWEHAPTLAT....EGATHDG.TGNRRG..D.....................................QRLQTAA............
uvig_384903_f;24457;25497                ............................................................................................................KKQVYDTLRF.
ERS698742_NODE_4_f;359;1546              .........................FNNCLN.......................DGSKCFDY.........................................RAAG............
MGV-GENOME-0279285_f;36563;37810         ...................IPAARPGPDERR.....TKRYGTVTFQSGKRQPGEAGGKQQ..A.....................................HQVHQAGQ...........
MGV-GENOME-0347346_f;54338;55741         ...................IE......NWQR...EEVPLYG..............................................................................
```
