## Supplementary material for "Diversity and ecology of *Caudoviricetes* phages with genome terminal repeats in fecal metagenomes from four Dutch cohorts": Text S1

**Benchmarking of virus detection and taxonomic assignment**

To benchmark virus detection, we applied the procedure relying on Cenote-Taker 2 to detect viral genomes (see Methods) to the Viral RefSeq 209 genomes. The percentage of detected viral genomes was noted (Table 1) for seven dsDNA viral groups that could potentially be found in the human gut: phylum *Nucleocytoviricota*, class *Caudoviricetes*, class *Tectiliviricetes* excluding family *Adenoviridae*, and families *Adenoviridae*, *Herpesviridae*, *Papillomaviridae* and *Polyomaviridae*. For five of the groups, more than 70% of the genomes were detected, reaching 99.61% for class *Caudoviricetes*. At the same time, almost no genomes belonging to families *Papillomaviridae* and *Polyomaviridae* were detected, possibly because almost all the Viral RefSeq 209 genome sequences of these viruses lack terminal redundancy, preventing bioinformatics tools from recognizing them as circular and thus from identifying them as viral.

To benchmark virus taxonomic assignment, we applied the procedure relying on marker genes to assign viral genomes to the seven taxonomic groups (see Methods, Figure S2) to the Viral RefSeq 209 proteomes. Sensitivity and specificity of the assignment to each group was noted (Table 1). Specificity exceeded 99% and sensitivity exceeded 70% for all groups.

It is important to note that the benchmarked procedures rely on marker protein alignments that may contain sequences included in the Viral RefSeq database. Consequently, the robustness of the procedures may be overestimated during the benchmarking. On the other hand, Viral RefSeq contains incomplete genomes missing the marker genes, which may explain why some of the genomes belonging to the selected groups remained undetected.
